# Chemoreceptor Co-Expression in *Drosophila* Olfactory Neurons

**DOI:** 10.1101/2020.11.07.355651

**Authors:** Darya Task, Chun-Chieh Lin, Alina Vulpe, Ali Afify, Sydney Ballou, Maria Brbić, Philipp Schlegel, Gregory S. X. E. Jefferis, Hongjie Li, Karen Menuz, Christopher J. Potter

## Abstract

*Drosophila melanogaster* olfactory neurons have long been thought to express only one chemosensory receptor gene family. There are two main olfactory receptor gene families in *Drosophila*, the Odorant Receptors (ORs) and the Ionotropic Receptors (IRs). The dozens of odorant binding receptors in each family require at least one co-receptor gene in order to function: *Orco* for ORs, and *Ir25a*, *Ir8a*, and *Ir76b* for IRs. Using a new genetic knock-in strategy, we targeted the four co-receptors representing the main chemosensory families in *Drosophila* (*Orco, Ir8a, Ir76b, Ir25a*). Co-receptor knock-in expression patterns were verified as accurate representations of endogenous expression. We find extensive overlap in expression among the different co-receptors. As defined by innervation into antennal lobe glomeruli, *Ir25a* is broadly expressed in 88% of all olfactory sensory neuron classes and is co-expressed in 82% of Orco+ neuron classes, including all neuron classes in the maxillary palp. *Orco*, *Ir8a*, and *Ir76b* expression patterns are also more expansive than previously assumed. Single sensillum recordings from Orco-expressing *Ir25a* mutant antennal and palpal neurons identify changes in olfactory responses. These results suggest co-expression of chemosensory receptors is common in olfactory neurons. Together, our data present the first comprehensive map of chemosensory co-receptor expression and reveal their unexpected widespread co-expression in the fly olfactory system.

## Introduction

The sense of smell is crucial for many animal behaviors, from conspecific recognition and mate choice (Dweck et al., 2015; Stengl, 2010), to location of a food source (Auer et al., 2020; Hansson and Stensmyr, 2011), to avoidance of predators (Kondoh et al., 2016; Papes et al., 2010) and environmental dangers (Mansourian et al., 2016; Stensmyr et al., 2012). Peripheral sensory organs detect odors in the environment using a variety of chemosensory receptors (Carey and Carlson, 2011; Su et al., 2009). The molecular repertoire of chemosensory receptors expressed by the animal, and the particular receptor expressed by any individual olfactory neuron, define the rules by which an animal interfaces with its odor environment. Investigating this initial step in odor detection is critical to understanding how odor signals first enter the brain to guide behaviors.

The olfactory system of the vinegar fly, *Drosophila melanogaster*, is one of the most extensively studied and well understood (Depetris-Chauvin et al., 2015). *Drosophila* is an attractive model for studying olfaction due to its genetic tractability, numerically simpler nervous system (compared to mammals), complex olfactory-driven behaviors, and similar organizational principles to vertebrate olfactory systems (Ache and Young, 2005; Wilson, 2013). Over sixty years of research have elucidated many of the anatomical, molecular, and genetic principles underpinning fly olfactory behaviors (Gomez-Diaz et al., 2018; Harris, 1972; Pask and Ray, 2016; Siddiqi, 1987; Stocker, 2001; Venkatesh and Singh, 1984; Vosshall and Stocker, 2007; Yan et al., 2020). Recent advances in electron microscopy and connectomics are revealing higher brain circuits involved in processing of olfactory information (Bates et al., 2020; Berck et al., 2016; Frechter et al., 2019; Horne et al., 2018; Marin et al., 2020; Zheng et al., 2018); such endeavors will aid the full mapping of neuronal circuits from sensory inputs to behavioral outputs.

The fly uses two olfactory appendages to detect odorants: the antennae and maxillary palps (Figure 1A) (Stocker, 1994). Each of these is covered by sensory hairs called sensilla, and each sensillum houses between one and four olfactory sensory neurons or OSNs (Figure 1B) (de Bruyne et al., 2001; Venkatesh and Singh, 1984). The dendrites of these neurons are found within the sensillar lymph, and they express chemosensory receptors from three gene families: *Odorant Receptors* (*ORs*), *Ionotropic Receptors* (*IRs*), and *Gustatory Receptors* (*GRs*) (Figure 1C, left) (Benton et al., 2009; Clyne et al., 1999; Gao and Chess, 1999; Jones et al., 2007; Kwon et al., 2007; Vosshall et al., 1999; Vosshall et al., 2000). These receptors bind odorant molecules that enter the sensilla from the environment, leading to the activation of the OSNs, which then send this olfactory information to the fly brain (Figure 1D), to the first olfactory processing center – the antennal lobes (ALs) (Figure 1E) (reviewed in: Depetris-Chauvin et al., 2015; Gomez-Diaz et al., 2018; Pask and Ray, 2016). The standard view regarding the organization of the olfactory system in *Drosophila* is that olfactory neurons express receptors from only one of the chemosensory gene families (either *ORs*, *IRs*, or *GRs*), and all neurons expressing the same receptor (which can be considered an OSN class) project their axons to one specific region in the AL called a glomerulus (Figure 1C, right) (Couto et al., 2005; Fishilevich and Vosshall, 2005; Gao et al., 2000; Laissue et al., 1999; Pinto et al., 1988; Vosshall et al., 2000). This pattern of projections creates a map in which the OR+ (Figure 1E, teal), IR+ (Figure 1E, purple), and GR+ (Figure 1E, dark blue) domains are segregated from each other in the AL. The OR+ domains innervate 38 anterior glomeruli, while the IR+ (19 glomeruli) and GR+ (1 glomerulus) domains occupy more posterior portions of the AL. One exception is the Or35a+ OSN class, which expresses an *IR* (*Ir76b*) in addition to the *OR* and *Orco*, and innervates the VC3 glomerulus (Figure 1E, striped glomerulus) (Benton et al., 2009; Couto et al., 2005; Fishilevich and Vosshall, 2005). Different OSN classes send their information to different glomeruli, and the specific combination of OSN classes and glomeruli that are activated by a given smell (usually a blend of different odorants) constitutes an olfactory ‘code’ which the fly brain translates into an appropriate behavior (Grabe and Sachse, 2018; Haverkamp et al., 2018; Seki et al., 2017).

**Figure 1.**
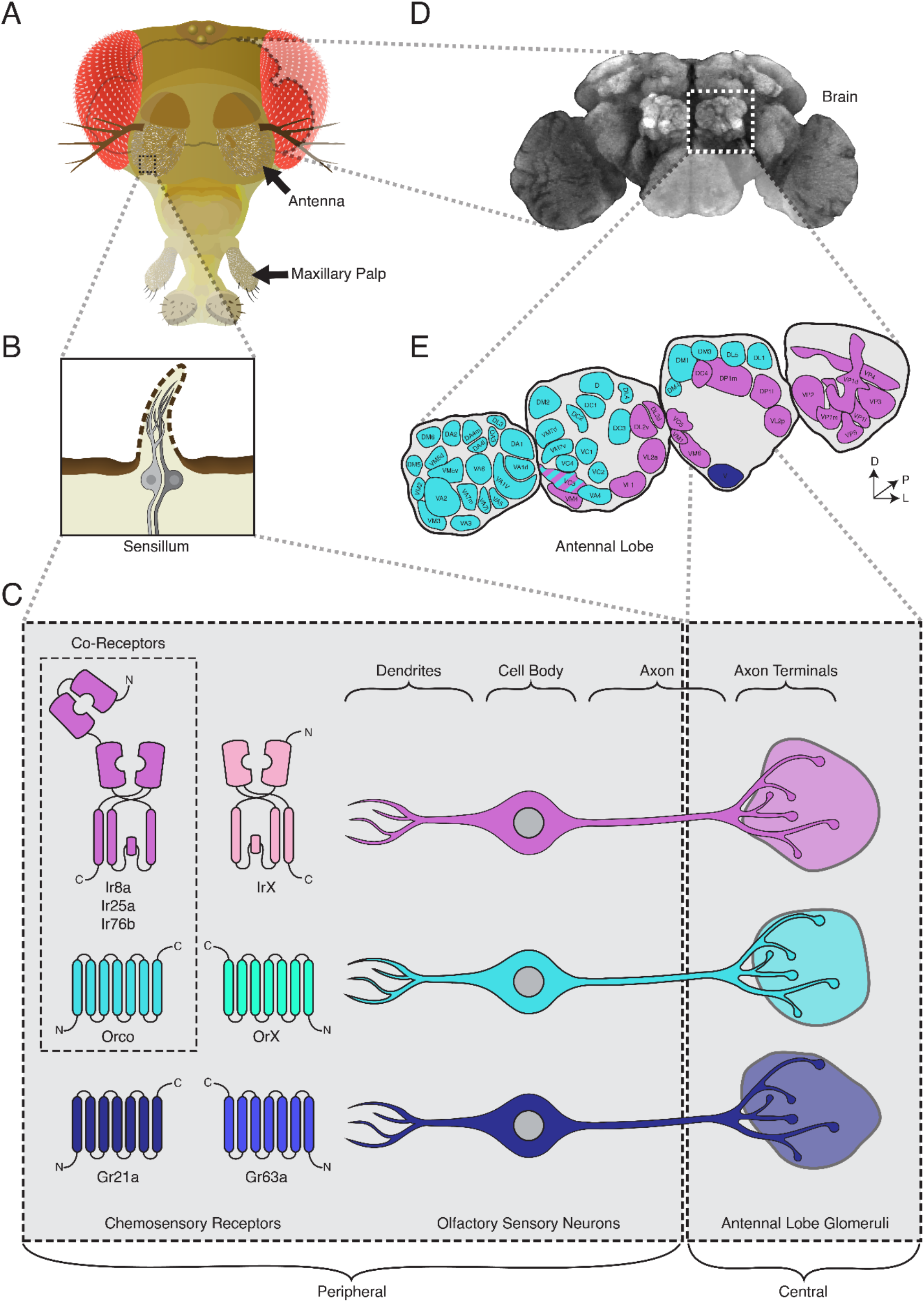
The Standard View of Olfactory Receptor Expression in *Drosophila melanogaster*. **A.** The adult fly head (left) has two olfactory organs: the antennae and the maxillary palps (arrows). Olfactory neurons from these organs project to the fly brain (D), to the first center involved in processing of olfactory information, the antennal lobes (E). **B.** The olfactory organs are covered by sensory hairs called sensilla (left). Each sensillum contains between one and four olfactory sensory neurons (two example neurons are shown in grey). The dendrites of these neurons extend into the sensilla, and the axons target discrete regions of the antennal lobes called glomeruli (E). Neuronal compartments (dendrites, cell body, axon, axon terminals) are labeled in (C). **C.** Left: in the periphery, each olfactory sensory neuron is traditionally thought to express chemosensory receptors from only one of three gene families on its dendrites: Ionotropic Receptors (IRs, pink and purple), Odorant Receptors (ORs, teal and green), or Gustatory Receptors (GRs, light and dark blue). IRs and ORs require obligate co-receptors (dotted box outline) to form functional ion channels. All ORs utilize a single co-receptor, Orco (teal), while IRs can utilize one (or a combination) of three possible co-receptors (purple): Ir8a, Ir25a, or Ir76b. The two GRs form a functional carbon dioxide detecting channel expressed in only one class of neurons. All other olfactory neurons express one of the four co-receptors. Right: olfactory sensory neurons expressing ORs, IRs, and GRs are thought to project to mutually exclusive glomeruli in the AL of the central brain, forming the olfactory map shown in (E). **D.** Fly brain stained with anti-brp synaptic marker (nc82), with left antennal lobe (AL) outlined by dotted white box. **E.** AL map with glomeruli color-coded by the chemosensory receptors (ORs, IRs, or GRs) expressed in the olfactory sensory neurons projecting to them. Only one glomerulus (VC3, striped) receives inputs from neurons expressing chemoreceptors from multiple gene families (ORs and IRs). Compass: D = dorsal, L = lateral, P = posterior.

The receptors within each chemosensory gene family form heteromeric ion channels (receptor complexes) (Abuin et al., 2011; Butterwick et al., 2018; Sato et al., 2008). The *ORs* require a single co-receptor, *Orco*, to function (Figure 1C, middle row) (Benton et al., 2006; Larsson et al., 2004; Vosshall and Hansson, 2011). The ligand-binding *OrX* confers odorant specificity upon the receptor complex, while the co-receptor *Orco* is necessary for trafficking of the *OrX* to the dendritic membrane and formation of a functional ion channel (Benton et al., 2006; Larsson et al., 2004). Likewise, the ligand-binding *IrXs* require one or more *IR* co-receptors: *Ir8a*, *Ir25a*, and/or *Ir76b* (Figure 1C, top row). The *IR* co-receptors are similarly required for trafficking and ion channel function (Abuin et al., 2011; Abuin et al., 2019; Ai et al., 2013; Vulpe et al., 2021b). The *GR* gene family generally encodes receptors involved in taste, which are typically expressed outside the olfactory system (such as in the labella or the legs) (Dunipace et al., 2001; Park and Kwon, 2011; Scott, 2018; Scott et al., 2001); however, *Gr21a* and *Gr63a* are expressed in one antennal OSN neuron class, and form a complex sensitive to carbon dioxide (Figure 1C, bottom row) (Jones et al., 2007; Kwon et al., 2007).

The majority of receptors have been mapped to their corresponding OSNs, sensilla, and glomeruli in the fly brain (Bhalerao et al., 2003; Couto et al., 2005; Fishilevich and Vosshall, 2005; Frank et al., 2017; Grabe et al., 2016; Hallem and Carlson, 2006; Hallem et al., 2004; Knecht et al., 2017; Marin et al., 2020; Ray et al., 2008; Silbering et al., 2011). This detailed map has allowed for exquisite investigations into the developmental, molecular, electrophysiological, and circuit/computational bases of olfactory neurobiology. This work has relied on transgenic lines to identify and manipulate OSN classes (Ai et al., 2013; Brand and Perrimon, 1993; Couto et al., 2005; Fishilevich and Vosshall, 2005; Kwon et al., 2007; Lai and Lee, 2006; Larsson et al., 2004; Menuz et al., 2014; Potter et al., 2010; Silbering et al., 2011). These transgenic lines use segments of DNA upstream of the chemosensory genes that are assumed to reflect the enhancers and promoters driving expression of these genes. While a powerful tool, transgenic lines may not contain all of the necessary regulatory elements to faithfully recapitulate the expression patterns of the endogenous genes. Some transgenic lines label a subset of the cells of a given olfactory class, while others label additional cells: for example, the transgenic *Ir25a-Gal4* line is known to label only a portion of cells expressing Ir25a protein (as revealed by antibody staining) (Abuin et al., 2011); conversely, *Or67d-Gal4* transgenes incorrectly label two glomeruli, whereas a *Gal4* knock-in at the *Or67d* genetic locus labels a single glomerulus (Couto et al., 2005; Fishilevich and Vosshall, 2005; Kurtovic et al., 2007). While knock-ins provide a faithful method to capture a gene’s expression pattern, generating these lines has traditionally been cumbersome.

In this paper, we implement an efficient knock-in strategy to target the four main chemosensory co-receptor genes in *Drosophila melanogaster (Orco, Ir8a, Ir76b, Ir25a)*. We find broad co-expression of these co-receptor genes in various combinations in olfactory neurons, challenging the current view of segregated olfactory families in the fly. In particular, *Ir25a* is expressed in the majority of olfactory neurons, including most Orco+ OSNs. In addition, the *Ir8a* and *Ir25a* knock-in lines help to distinguish two new OSN classes in the sacculus that target previously un-identified glomerular subdivisions in the posterior AL. Recordings in *Ir25a* mutant sensilla in Orco+ neurons reveal subtle changes in odor responses, suggesting that multiple chemoreceptor gene families could be involved in the signaling or development of a given OSN class. These data invite a re-examination of odor coding in *Drosophila*. We present a comprehensive model of co-receptor expression, which will inform future investigations of combinatorial chemosensory processing.

## Results

### Generation and Validation of Co-Receptor Knock-in Lines

We previously developed the HACK technique for CRISPR/Cas9-mediated *in vivo* gene conversion of binary expression system components, such as the conversion of transgenic *Gal4* to *QF2* (Brand and Perrimon, 1993; Jinek et al., 2012; Lin and Potter, 2016a, b; Potter et al., 2010; Riabinina et al., 2015; Xie et al., 2018). Here, we adapt this strategy for the efficient generation of targeted knock-ins (see Table 1 and Table 1-Source Data 1 for details). We chose to target the four chemosensory co-receptor genes to examine unmapped patterns of co-receptor expression in *Drosophila melanogaster*. We inserted a *T2A-QF2* cassette and mCherry selection marker before the stop codon of the four genes of interest (Figure 2A and Figure 2-Figure Supplement 1). By introducing the T2A ribosomal skipping peptide, the knock-in will produce the full-length protein of the gene being targeted as well as a functional QF2 transcription factor (Figure 2A, Protein Products). This approach should capture the endogenous expression pattern of the gene under the control of the gene’s native regulatory elements, while retaining the gene’s normal function (Baena-Lopez et al., 2013; Bosch et al., 2020; Chen et al., 2020; Diao et al., 2015; Diao and White, 2012; Du et al., 2018; Gnerer et al., 2015; Gratz et al., 2014; Kanca et al., 2019; Lee et al., 2018; Li-Kroeger et al., 2018; Lin and Potter, 2016a; Vilain et al., 2014; Xue et al., 2014). We found that *T2A-QF2* knock-ins were functional with some exceptions (See Figure 2-Figure Supplement 2 and Figure 2-Source Data 1). For example, *Orco-T2A-QF2* knock-in physiology was normal, while a homozygous *Ir25a-T2A-QF2* knock-in exhibited a mutant phenotype. This suggests that the addition of the T2A peptide onto the C-terminus of Ir25a might interfere with its co-receptor function.

**Figure 2.**
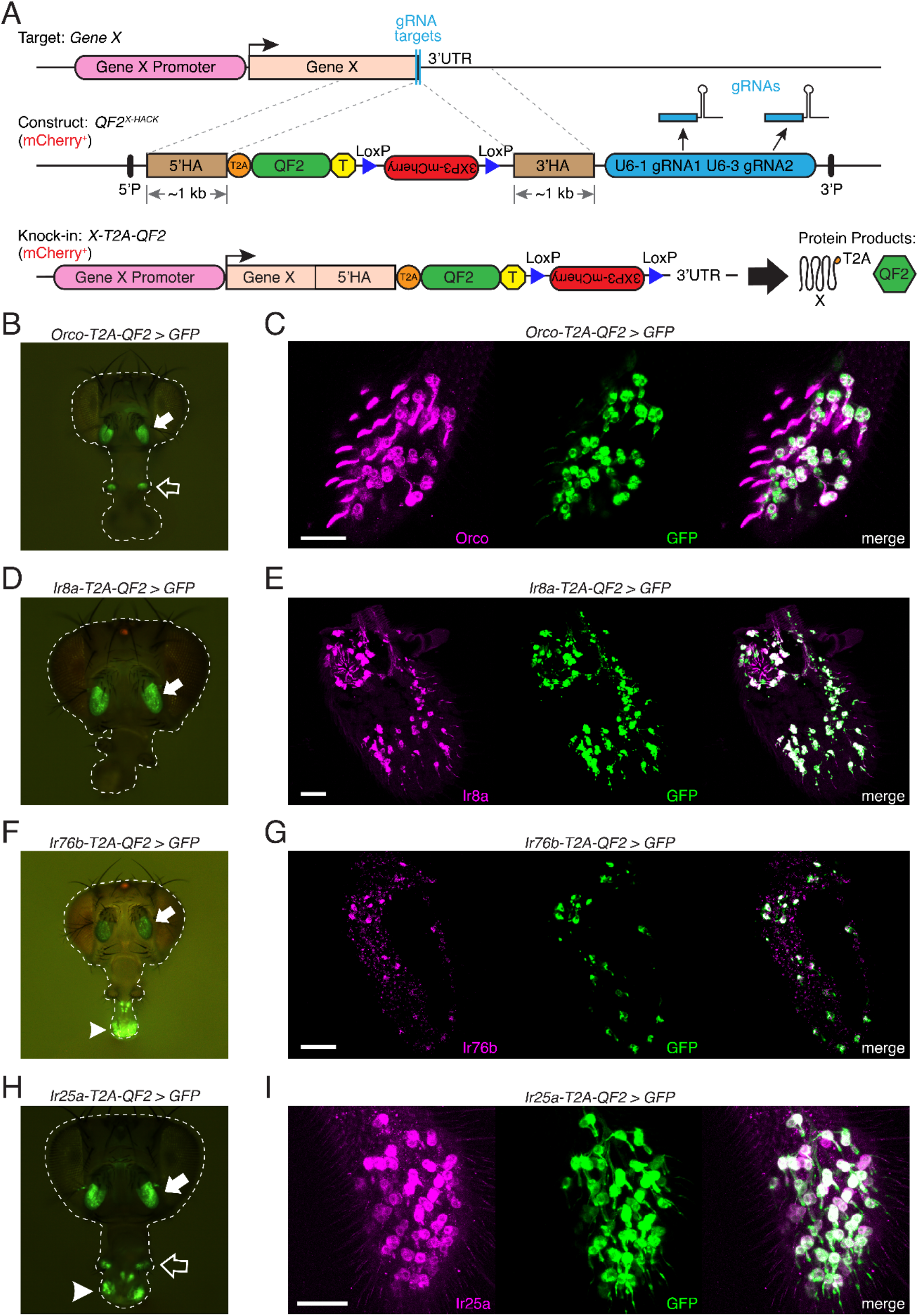
Generation and Validation of Chemosensory Co-Receptor Knock-in Lines. **A.** Schematic of HACK knock-in approach. Top: two double-stranded breaks are induced on either side of the target gene stop codon with gRNAs (blue) expressed from the *QF2^X-HACK^* construct (middle) in the presence of Cas9. The construct includes *T2A-QF2* and a floxed *3XP3-mCherry* marker. The knock-in introduces a transcriptional stop (yellow T) after QF2. Bottom: the knock-in produces two protein products (right) from the targeted mRNA: target X and the *QF2* transcription factor (Diao and White, 2012). The *X-T2A-QF2* knock-in can be crossed to a reporter (e.g. *QUAS-GFP*) to examine the endogenous expression pattern of the target gene. **B.** *Orco-T2A-QF2* driving *QUAS-GFP* in adult fly head. GFP expression is found in the antennae (filled arrow) and maxillary palps (hollow arrow), as previously reported (Larsson et al., 2004). **C.** Whole-mount anti-Orco antibody staining in *Orco-T2A-QF2*>*GFP* maxillary palps reveals a high degree of overlap of Orco+ and GFP+ cells. N = 3. **D.** *Ir8a-T2A-QF2* drives GFP in the antennae (arrow), as previously reported (Abuin et al., 2011). **E.** Anti-Ir8a antibody staining of *Ir8a-T2A-QF2*>*GFP* antennal cryosections shows high correspondence between Ir8a+ and GFP+ cells. N = 7. **F.** *Ir76b-T2A-QF2* drives GFP expression in the antennae (filled arrow) and labella (hollow arrow), reflecting *Ir76b*’s role in olfaction and gustation, respectively (Benton et al., 2009; Zhang et al., 2013). **G.** *In situs* on *Ir76b-T2A-QF2*>*GFP* antennal cryosections to validate that the knock-in faithfully recapitulates the endogenous expression pattern. N = 3. **H.** *Ir25a-T2A-QF2* drives GFP in the antennae (filled arrow) and labella (hollow arrow), which has been reported previously (Benton et al., 2009; Croset et al., 2010). Expression in the maxillary palps (arrowhead) has not been previously reported. **I.** Whole-mount maxillary palp staining with an anti-Ir25a antibody in *Ir25a-T2A-QF2>GFP* flies. The knock-in and Ir25a antibody co-labeled the majority of olfactory neurons in the palps. N = 5. Scale bars = 25 µm. In (D) and (F), the *3XP3-mCherry* knock-in marker can be weakly detected in the eyes and ocelli (red spot) of both *Ir8a-T2A-QF2* and *Ir76b-T2A-QF2*. See also Figure 2-Figure Supplements 1-4, Tables 1 and 2, Table 1-Source Data 1, Figure 2-Source Data 1, and Methods.

**Table 1.** Summary of HACK Knock-in Efficiency. Related to Figure 2. There are two ways to generate knock-ins via the HACK technique: by direct injection or by genetic cross (see Figure 2-Figure Supplement 1 and Methods for details) (Lin and Potter, 2016a). All four co-receptor genes were targeted using the direct injection approach; additionally, the crossing approach was tested with *Orco* and *Ir25a*. Knock-in efficiency, as measured by the number of flies having the mCherry+ marker divided by the total number of potentially HACKed flies, was high for all genes tested and both approaches. Efficiency appears to depend on the genetic locus, as has been previously demonstrated (Lin and Potter, 2016a). To further estimate the effort required to generate a HACK knock-in, we calculated the percentage of founder flies producing knock-in lines; this gives an indication of the number of independent crosses needed to successfully create a knock-in line. Two to five individual G_0_ starting crosses were sufficient to produce a knock-in. For each gene, a sample of individual knock-in lines were tested via PCR genotyping, sequencing, and by crossing to a reporter line to confirm brain expression (knock-ins sampled). For all knock-ins generated via the direct injection method, every fly tested represented a correctly targeted knock-in. However, for the cross method, some lines had the mCherry+ marker and yet did not drive GFP expression in the brain when crossed to a reporter line (labeled here as false positives). See also Table 1-Source Data 1 and Figure 2-Figure Supplement 1.

| Gene | Approach | mCherry+ | mCherry- | Total | Efficiency | Founders<br>Producing<br>Knock-in<br>(#/Total) | Knock-<br>ins<br>Sampled | False<br>Positives | Confirmed | Correct |
| --- | --- | --- | --- | --- | --- | --- | --- | --- | --- | --- |
| *Orco* | Direct<br>Injection | 180 | 365 | 545 | 33% | 43% (3/7) | 30 | 0 | 30 | 100% |
| *Ir8a* | Direct<br>Injection | 53 | 609 | 662 | 8% | 20% (4/20) | 5 | 0 | 5 | 100% |
| *Ir76b* | Direct<br>Injection | 79 | 184 | 263 | 30% | 100% (2/2) | 10 | 0 | 10 | 100% |
| *Ir25a* | Direct<br>Injection | 82 | 268 | 350 | 23% | 40% (2/5) | 6 | 0 | 6 | 100% |
| *Orco* | Cross | 37 | 96 | 133 | 28% | 100% (3/3) | 2 | 1 | 1 | 50% |
| *Ir25a* | Cross | 30 | 95 | 125 | 24% | 100% (2/2) | 30 | 5 | 25 | 83% |

We examined the expression of the co-receptor knock-in lines in the adult olfactory organs by crossing each line to the same *10XQUAS-6XGFP* reporter (Figure 2B – I). *Orco-T2A-QF2* driven GFP expression was detected in the adult antennae and maxillary palps (Figure 2B), as previously described (Larsson et al., 2004). We validated the *Orco-T2A-QF2* knock-in line with whole-mount antibody staining of maxillary palps (Figure 2C) and found a high degree of correspondence between anti-Orco antibody staining and knock-in driven GFP in palpal olfactory neurons (quantified in Table 2; see also Figure 2-Figure Supplement 3A – D for PCR and sequencing validation of all knock-in lines). We confirmed the specificity of the anti-Orco antibody by staining *Orco^2^* mutant palps and found no labeling of olfactory neurons (Figure 2-Figure Supplement 3E).

**Table 2.** Validation of T2A-QF2 Knock-in Expression in the Antennae and Maxillary Palps. Related to Figure 2. To verify that the knock-in lines recapitulate the endogenous expression patterns of the target genes, antennae or maxillary palps of flies containing the knock-ins driving GFP expression were co-stained with the corresponding antibody (Ab) (anti-Orco, anti-Ir8a, or anti-Ir25a). The overlap of Ab+ and GFP+ cells was examined, and a high correspondence between antibody staining and knock-in driven GFP was found. *WM = whole-mount, Cryo = cryosection. See also Figure 2-Figure Supplement 3.

| Knock-in | Sample* | Antibody<sup>†</sup> (Ab) | Ab+ cells | GFP+ cells | Double-labeled cells | Total cells |
| --- | --- | --- | --- | --- | --- | --- |
| ***Orco*** | Palp 1 (WM) | anti-Orco | 125 | 127 | 125 | 127 |
| ***Orco*** | Palp 2 (WM) | anti-Orco | 112 | 111 | 108 | 115 |
| ***Orco*** | Palp 6 (WM) | anti-Orco | 125 | 126 | 123 | 128 |
|  |  | Total across samples: | 362 | 364 | 356 | 370 |
|  |  |  | Proportion of Ab+ cells that are GFP+: | Proportion of GFP+ cells that are Ab+: | Proportion of all cells that are double labeled: |  |
|  |  |  | **0.98** | **0.98** | **0.96** |  |
| ***Ir8a*** | Antenna 1 (Cryo) | anti-Ir8a | 20 | 21 | 20 | 21 |
| ***Ir8a*** | Antenna 2 (Cryo) | anti-Ir8a | 24 | 24 | 24 | 24 |
| ***Ir8a*** | Antenna 6 (Cryo) | anti-Ir8a | 40 | 43 | 40 | 43 |
| ***Ir8a*** | Antenna 7 (Cryo) | anti-Ir8a | 12 | 13 | 12 | 13 |
| ***Ir8a*** | Antenna 8 (Cryo) | anti-Ir8a | 16 | 16 | 16 | 16 |
| ***Ir8a*** | Antenna 9 (Cryo) | anti-Ir8a | 42 | 42 | 41 | 43 |
| ***Ir8a*** | Antenna 10 (Cryo) | anti-Ir8a | 41 | 40 | 40 | 41 |
|  |  | Total across samples: | 195 | 199 | 193 | 201 |
|  |  |  | Proportion of Ab+ cells that are GFP+: | Proportion of GFP+ cells that are Ab+: | Proportion of all cells that are double labeled: |  |
|  |  |  | **0.99** | **0.97** | **0.96** |  |
| ***Ir25a*** | Palp 1 (WM) | anti-Ir25a | 107 | 105 | 104 | 108 |
| ***Ir25a*** | Palp 2 (WM) | anti-Ir25a | 86 | 85 | 85 | 86 |
| ***Ir25a*** | Palp 3 (WM) | anti-Ir25a | 111 | 111 | 110 | 112 |
| ***Ir25a*** | Palp 4 (WM) | anti-Ir25a | 94 | 94 | 94 | 94 |
| ***Ir25a*** | Palp 5 (WM) | anti-Ir25a | 83 | 83 | 81 | 85 |
|  |  | Total across samples: | 481 | 478 | 474 | 485 |
|  |  |  | Proportion of Ab+ cells that are GFP+: | Proportion of GFP+ cells that are Ab+: | Proportion of all cells that are double labeled: |  |
|  |  |  | **0.99** | **0.99** | **0.98** |  |

Unlike *Orco*, *Ir8a* expression has previously been localized only to the antenna, to olfactory neurons found in coeloconic sensilla and in the sacculus (Abuin et al., 2011). As expected, the knock-in line drove GFP expression only in the antenna (Figure 2D). To validate the *Ir8a-T2A-QF2* knock-in line, we performed antibody staining on antennal cryosections and found the majority of cells to be double-labeled (Figure 2E, Table 2). There was no anti-Ir8a staining in control *Ir8a^1^* mutant antennae (Figure 2-Figure Supplement 3F).

The *Ir76b* gene has previously been implicated in both olfaction and gustation and has been shown to be expressed in adult fly antennae, labella (mouthparts), legs, and wings (Abuin et al., 2011; Chen and Amrein, 2017; Croset et al., 2010; Ganguly et al., 2017; Hussain et al., 2016; Sanchez-Alcaniz et al., 2018; Zhang et al., 2013). We examined the *Ir76b-T2A-QF2* knock-in line and found a similar pattern of expression in the periphery, with GFP expression in the antennae and labella (Figure 2F). Because an anti-Ir76b antibody has not previously been tested in fly antennae, we performed *in situs* on *Ir76b-T2A-QF2 > GFP* antennal cryosections to validate knock-in expression (Figure 2G) and confirmed the specificity of the probe in *Ir76b^1^* mutant antennae (Figure 2-Figure Supplement 3G).

Of the four *Drosophila* co-receptor genes, *Ir25a* has been implicated in the broadest array of cellular and sensory functions, from olfaction (Abuin et al., 2011; Benton et al., 2009; Silbering et al., 2011) and gustation (Chen and Amrein, 2017; Chen and Dahanukar, 2017; Jaeger et al., 2018), to thermo- and hygro-sensation (Budelli et al., 2019; Enjin et al., 2016; Knecht et al., 2017; Knecht et al., 2016), to circadian rhythm modulation (Chen et al., 2015). In the adult olfactory system, *Ir25a* expression has previously been reported in three types of structures in the antenna: coeloconic sensilla, the arista, and the sacculus (Abuin et al., 2011; Benton et al., 2009). We examined the *Ir25a-T2A-QF2* knock-in line and found GFP expression in the adult antennae, labella, and maxillary palps (Figure 2H). This was surprising because no Ionotropic Receptor expression has previously been reported in fly palps. To verify Ir25a protein expression in the maxillary palps, we performed whole-mount anti-Ir25a antibody staining in *Ir25a-T2A-QF2 > GFP* flies. We found broad *Ir25a* expression in palpal olfactory neurons (Figure 2I), and a high degree of overlap between knock-in driven GFP expression and antibody staining (Table 2). As expected, there was no anti-Ir25a staining in *Ir25a^2^* mutant palps (Figure 2-Figure Supplement 3H).

We also examined co-receptor knock-in expression in *Drosophila* larvae. As in the adult stage, larval GFP expression was broadest in the *Ir25a-T2A-QF2* and *Ir76b-T2A-QF2* knock-in lines, with GFP labeling of neurons in the head and throughout the body wall (Figure 2-Figure Supplement 4). The *Orco-T2A-QF2* knock-in line labeled only the olfactory dorsal organs in the larva, while the *Ir8a-T2A-QF2* knock-in line did not have obvious expression in the larval stage (Figure 2-Figure Supplement 4). All subsequent analyses focused on the adult olfactory system.

### Expanded Expression of Olfactory Co-Receptors

We next examined the innervation patterns of the four co-receptor knock-in lines in the adult central nervous system: the brain and ventral nerve cord (VNC) (Figure 3). Only two of the four lines (*Ir25a* and *Ir76b*) showed innervation in the VNC, consistent with the role of these genes in gustation in addition to olfaction (Figure 3-Figure Supplement 1A). In the brain, we compared the expression of each knock-in line (Figure 3A – D, green) to the corresponding transgenic *Gal4* line (Figure 3A – D, orange) to examine differences in expression to what has previously been reported. Reporter alone controls for these experiments are shown in Figure 3-Figure Supplement 1B. All four knock-in lines innervated the ALs, and the *Ir25a-T2A-QF2* and *Ir76b-T2A-QF2* lines additionally labeled the suboesophageal zone (SEZ), corresponding to gustatory axons from the labella (Figure 3C and D, arrowheads) (Hussain et al., 2016; Zhang et al., 2013). The co-labeling experiments revealed that all four knock-ins label more glomeruli than previously reported (see Figure 3-Source Data 1 for AL analyses, Figure 3-Source Data 2 for traced examples of newly identified glomeruli in each knock-in line, and Table 3 for a summary of glomerular expression across all knock-in lines). Some glomeruli were not labeled consistently in all flies, which we define as variable expression (found in <50% of brains examined).

**Figure 3.**
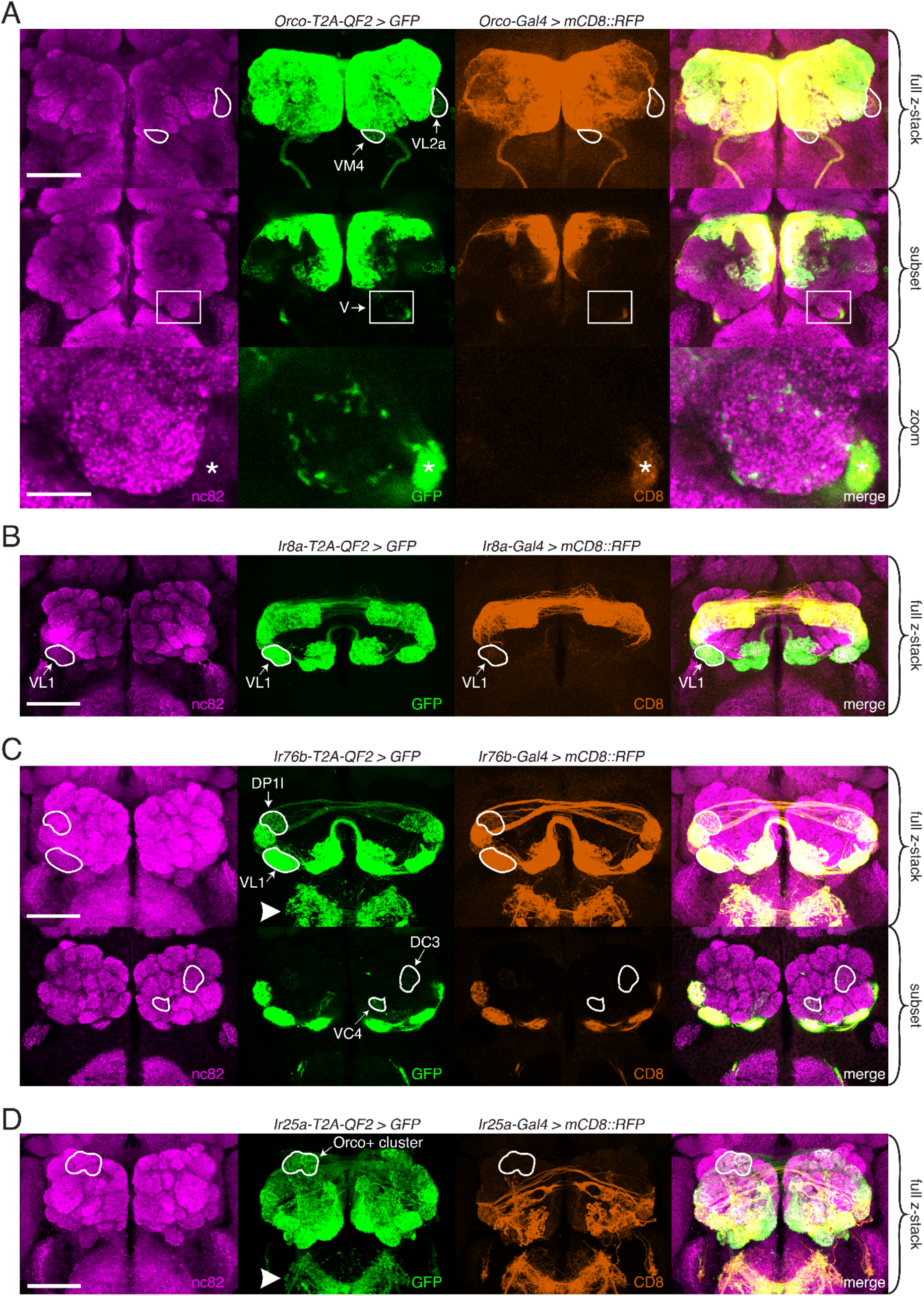
Expanded Expression of Olfactory Co-Receptors. **A-D.** Comparing knock-in innervation patterns of the AL with what has previously been reported for each co-receptor. Co-labeling experiments with each co-receptor knock-in line driving *QUAS-GFP* (green) and the corresponding transgenic co-receptor *Gal4* line driving *UAS-mCD8::RFP* (anti-CD8, orange). The nc82 antibody labels synapses (magenta) and is used as a brain counterstain in these and all subsequent brain images. **A.** The *Orco-T2A-QF2* knock-in labels more glomeruli than the *Orco-Gal4* line. Top: maximum intensity projection of full z-stack showing two additional glomeruli labeled by the knock-in, VM4 (Ir8a+/Ir76b+/Ir25a+) and VL2a (Ir8a+). Middle: subset of z-stack with a box around the V glomerulus. Bottom: zoom of boxed region showing sparse innervation of the V glomerulus (Gr21a+/Gr63a+) by the knock-in but not the *Gal4* line. Asterisk indicates antennal nerve which is outside the V glomerulus. In the sub z-stack and zoom panel, gain has been increased in the GFP channel to visualize weak labeling more clearly. **B.** The *Ir8a-T2A-QF2* knock-in also drives GFP expression in more glomeruli than previously reported, including the outlined VL1 glomerulus (Ir25a+). **C.** In the brain, *Ir76b-T2A-QF2 > GFP* olfactory neurons innervate the antennal lobes (AL), while gustatory neurons from the labella innervate the subesophageal zone (SEZ, arrowhead). Top: Both the *Ir76b* knock-in and transgenic *Gal4* line label more glomeruli than previously reported, including VL1 (Ir25a+) and DP1l (Ir8a+). Bottom: The *Ir76b-T2A-QF2* knock-in labels several Orco+ glomeruli, such as DC3 and VC4 (outlined). In the subset, gain has been increased in the GFP channel to visualize weakly labeled glomeruli more clearly. **D.** The *Ir25a-T2A-QF2* knock-in drives GFP expression broadly in the antennal lobes and SEZ (arrowhead). Ir25a+ neurons innervate many Orco+ glomeruli, such as those outlined. The transgenic *Ir25a-Gal4* line labels a subset of the knock-in expression pattern. N = 3 – 10 for co-labeling experiments, N = 5 – 15 for additional analyses of the knock-in lines alone. Scale bars = 25 µm, except zoom panel scale bar = 10 µm. See also Figure 3-Figure Supplements 1 and 2, Table 3, and Figure 3-Source data 1 and 2.

**Table 3.** Summary of Expression Patterns for All Knock-in Lines. Related to Figures 3 – 5. Summarized here are all of the OSN classes innervating the 58 antennal lobe glomeruli^†^; their corresponding sensilla and tuning receptors; the previously reported (original) co-receptors they express; and whether or not each of the co-receptor knock-in lines labels those glomeruli. Variable indicates that the glomerulus was labeled in <50% of brains examined in the given knock-in line. Sensilla or glomeruli that have been renamed or reclassified have their former nomenclature listed in parentheses. Question marks indicate expression that has been reported but not functionally validated. ^†^The VM6 subdivisions (VM6v, VM6m, VM6l) are separated in this table for clarity but counted together as one glomerulus in accordance with Schlegel et al., 2021. *VM6l was initially named VC6 in version 1 of our pre-print (Task et al., 2020) but was reclassified using additional data from EM reconstructions in the AL and immunohistochemical experiments in the periphery (see Figure 5). See also Figure 3-Source Data 1 and 2.

| Glomerulus<sup>†</sup> | Sensillum | Tuning Receptor(s) | Original Co-receptor(s) | *Orco*-*T2A*-*QF2* | *Ir8a*-*T2A*-*QF2* | *Ir76b*-*T2A*-*QF2* | *Ir25a*-*T2A*-*QF2* | References |
| --- | --- | --- | --- | --- | --- | --- | --- | --- |
| D | Ab9A | Or69aA, Or69aB | Orco | Yes | Variable | No | No | (Couto et al., 2005; Fishilevich and Vosshall, 2005) |
| DA1 | At1A | Or67d | Orco | Yes | No | Variable | Yes | (Couto et al., 2005; Fishilevich and Vosshall, 2005; Kurtovic et al., 2007) |
| DA2 | Ab4B | Or56a, Or33a | Orco | Yes | No | No | Yes | (Couto et al., 2005; Fishilevich and Vosshall, 2005) |
| DA3 | Ai2B (At2B) | Or23a | Orco | Yes | No | No | Yes | (Couto et al., 2005; Fishilevich and Vosshall, 2005; Lin and Potter, 2015) |
| DA4l | Ai3C (At3C) | Or43a | Orco | Yes | No | No | Yes | (Couto et al., 2005; Fishilevich and Vosshall, 2005; Lin and Potter, 2015) |
| DA4m | Ai3B (At3B) | Or2a | Orco | Yes | No | No | Yes | (Couto et al., 2005; Lin and Potter, 2015) |
| DC1 | Ai3A (At3A) | Or19a, Or19b | Orco | Yes | No | No | Yes | (Couto et al., 2005; Fishilevich and Vosshall, 2005; Lin and Potter, 2015) |
| DC2 | Ai1A (Ab6A) | Or13a | Orco | Yes | No | No | Yes | (Couto et al., 2005; Fishilevich and Vosshall, 2005; Lin and Potter, 2015) |
| DC3 | Ai2A (At2A) | Or83c | Orco | Yes | No | Yes | Variable | (Couto et al., 2005; Fishilevich and Vosshall, 2005; Lin and Potter, 2015) |
| DL1 | Ab1D | Or10a, Gr10a | Orco | Yes | Yes | Yes | Yes | (Couto et al., 2005; Fishilevich and Vosshall, 2005) |
| DL3 | At4B | Or65a, Or65b, Or65c | Orco | Yes | No | No | Yes | (Couto et al., 2005; Fishilevich and Vosshall, 2005) |
| DL4 | Ab10B | Or49a, Or85f | Orco | Yes | No | No | Yes | (Couto et al., 2005; Fishilevich and Vosshall, 2005) |
| DL5 | Ab4A | Or7a | Orco | Yes | No | No | Variable | (Couto et al., 2005) |
| DM1 | Ab1A | Or42b | Orco | Yes | No | No | Yes | (Couto et al., 2005; Fishilevich and Vosshall, 2005) |
| DM2 | Ab3A | Or22a, Or22b | Orco | Yes | No | No | Yes | (Couto et al., 2005; Fishilevich and Vosshall, 2005) |
| DM3 | Ab5B | Or47a, Or33b | Orco | Yes | No | No | No | (Couto et al., 2005; Fishilevich and Vosshall, 2005) |
| DM4 | Ab2A | Or59b | Orco | Yes | No | No | Yes | (Couto et al., 2005) |
| DM5 | Ab2B | Or85a, Or33b | Orco | Yes | No | No | No | (Couto et al., 2005; Fishilevich and Vosshall, 2005) |
| DM6 | Ab10A | Or67a | Orco | Yes | No | No | Yes | (Couto et al., 2005) |
| VA1d | At4C | Or88a | Orco | Yes | No | No | Yes | (Couto et al., 2005; Fishilevich and Vosshall, 2005) |
| VA1v | At4A | Or47b | Orco | Yes | No | No | Yes | (Couto et al., 2005; Fishilevich and Vosshall, 2005) |
| VA2 | Ab1B | Or92a | Orco | Yes | No | No | Yes | (Couto et al., 2005; Fishilevich and Vosshall, 2005) |
| VA3 | Ab9B | Or67b | Orco | Yes | Yes | Yes | Yes | (Couto et al., 2005; Fishilevich and Vosshall, 2005) |
| VA4 | Pb3B | Or85d | Orco | Yes | No | No | Yes | (Couto et al., 2005) |
| VA5 | Ai1B (Ab6B) | Or49b | Orco | Yes | Yes | Yes | Yes | (Couto et al., 2005; Lin and Potter, 2015) |
| VA6 | Ab5A | Or82a | Orco | Yes | Yes | Yes | Yes | (Couto et al., 2005; Fishilevich and Vosshall, 2005) |
| VA7l | Pb2B | Or46a | Orco | Yes | No | No | Yes | (Couto et al., 2005; Fishilevich and Vosshall, 2005) |
| VA7m | UNK | UNK | Orco | Yes | No | Variable | Yes | (Couto et al., 2005) |
| VC1 | Pb2A | Or33c,<br>Or85e | Orco | Yes | No | No | Yes | (Couto et al., 2005;<br>Fishilevich and Vosshall,<br>2005) |
| VC2 | Pb1B | Or71a | Orco | Yes | No | No | Yes | (Couto et al., 2005;<br>Fishilevich and Vosshall,<br>2005) |
| VC4 | Ab7B | Or67c | Orco | Yes | No | Yes | Yes | (Couto et al., 2005) |
| VM2 | Ab8A | Or43b | Orco | Yes | No | No | No | (Couto et al., 2005) |
| VM3 | Ab8B | Or9a | Orco | Yes | No | No | No | (Couto et al., 2005) |
| VM5d | Ab3B | Or85b?,<br>Or98b? | Orco | Yes | Variable | No | Yes | (Couto et al., 2005) |
| VM5v | Ab7A | Or98a | Orco | Yes | Yes | No | Yes | (Couto et al., 2005;<br>Fishilevich and Vosshall,<br>2005) |
| VM7d | Pb1A | Or42a | Orco | Yes | No | No | Yes | (Couto et al., 2005; Endo et<br>al., 2007; Fishilevich and<br>Vosshall, 2005) |
| VM7v (1) | Pb3A | Or59c | Orco | Yes | No | No | Yes | (Couto et al., 2005; Endo et<br>al., 2007) |
| VC3 | Ac3B | Or35a | Orco, Ir76b | Yes | Yes | Yes | Yes | (Couto et al., 2005;<br>Fishilevich and Vosshall,<br>2005; Silbering et al., 2011) |
| V | Ab1C | Gr21a,<br>Gr63a | N/A | Yes | No | No | Yes | (Couto et al., 2005; Jones et<br>al., 2007; Kwon et al., 2007) |
| DC4 | Sacculus,<br>Chamber III | Ir64a | Ir8a | Variable | Yes | No | Yes | (Ai et al., 2013; Ai et al.,<br>2010; Silbering et al., 2011) |
| DL2d | Ac3A | Ir75b | Ir8a | Yes | Yes | No | Yes | (Prieto-Godina et al., 2017;<br>Silbering et al., 2011) |
| DL2v | Ac3A | Ir75c | Ir8a | Yes | Yes | No | Yes | (Prieto-Godina et al., 2017;<br>Silbering et al., 2011) |
| DP1l | Ac2 | Ir75a | Ir8a | Yes | Yes | Yes | Yes | (Silbering et al., 2011) |
| DP1m | Sacculus,<br>Chamber III | Ir64a | Ir8a | No | Yes | Yes | Yes | (Ai et al., 2013; Ai et al.,<br>2010; Silbering et al., 2011) |
| VL2a | Ac4 | Ir84a | Ir8a | Yes | Yes | Yes | Yes | (Silbering et al., 2011) |
| VL2p | Ac1 | Ir31a | Ir8a | No | Yes | Yes | Yes | (Silbering et al., 2011) |
| VC5 | Ac2 | Ir41a | Ir8a, Ir25a, Ir76b | No | Yes | Yes | Yes | (Hussain et al., 2016; Min et al., 2013; Silbering et al., 2011) |
| VM1 | Ac1 | Ir92a | Ir8a, Ir25a, Ir76b | No | Yes | Yes | Yes | (Min et al., 2013; Silbering et al., 2011) |
| VM4 | Ac4 | Ir76a | Ir8a, Ir25a, Ir76b | Yes | Yes | Yes | Yes | (Benton et al., 2009; Min et al., 2013; Silbering et al., 2011) |
| VL1 | Ac1, Ac2, Ac4 | Ir75d | Ir25a | Yes | Yes | Yes | Yes | (Silbering et al., 2011) |
| VM6v (VM6) | Ac1 | Rh50, Amt | Ir25a | No | Yes (weak) | No | Yes | (Chai et al., 2019; Li et al., 2016; Schlegel et al., 2021; Vulpe et al., 2021), This paper |
| VM6m (new) | Sacculus, Chamber III | Rh50, Amt | N/A (this paper) | No | Yes (weak) | No | Yes | (Chai et al., 2019; Li et al., 2016; Schlegel et al., 2021; Vulpe et al., 2021), This paper |
| VM6l* (new) | Sacculus, Chamber III | Rh50, Amt | N/A (this paper) | No | Yes (strong) | No | Yes | (Chai et al., 2019; Li et al., 2016; Schlegel et al., 2021; Vulpe et al., 2021), This paper |
| VP1d | Sacculus, Chamber II | Ir40a, Ir93a | Ir25a | No | No | No | Yes | (Enjin et al., 2016; Frank et al., 2017; Knecht et al., 2017; Knecht et al., 2016; Marin et al., 2020; Silbering et al., 2011) |
| VP1l | Sacculus, Chamber I | Ir21a, Ir93a | Ir25a | No | No | No | Yes | (Frank et al., 2017; Knecht et al., 2017; Knecht et al., 2016; Marin et al., 2020; Silbering et al., 2011) |
| VP1m | Sacculus, Chamber I | Ir68a, Ir93a | Ir25a | No | No | No | Yes | (Frank et al., 2017; Knecht et al., 2017; Knecht et al., 2016; Marin et al., 2020; Silbering et al., 2011) |
| VP2 | Arista | Gr28b.d, Ir93a | Ir25a | No | No | No | Yes | (Enjin et al., 2016; Frank et al., 2017; Marin et al., 2020; Miwa et al., 2018; Ni et al., 2013) |
| VP3 | Arista | Ir21a, Ir93a | Ir25a | No | No | No | Yes | (Budelli et al., 2019; Enjin et al., 2016; Frank et al., 2017; Silbering et al., 2011) |
| VP4 | Sacculus, Chambers I + II | Ir40a, Ir93a | Ir25a | No | No | No | Yes | (Enjin et al., 2016; Frank et al., 2017; Knecht et al., 2017; Knecht et al., 2016; Marin et al., 2020; Silbering et al., 2011) |
| VP5 | Sacculus, Chamber II | Ir68a, Ir93a | Ir25a | No | No | No | Yes | (Frank et al., 2017; Knecht et al., 2017; Marin et al., 2020) |

*Orco-T2A-QF2* labels seven ‘non-canonical’ glomeruli consistently, and one sporadically. These include VM4 and VL2a, which correspond to Ir76b+ and Ir8a+ OSN populations, respectively (Figure 3A, outlines). We also found that the *Orco* knock-in sparsely but consistently labels the V glomerulus, which is innervated by Gr21a+/Gr63a+ neurons (Figure 3A, box and zoom panel). *Orco-T2A-QF2* also labels one Ir25a+ glomerulus consistently (VL1), three additional Ir8a+ glomeruli consistently (DL2d, DL2v, DP1l) and one variably (DC4). Surprisingly, when we crossed the transgenic *Orco-Gal4* line (Larsson et al., 2004) to a stronger reporter (Shearin et al., 2014), we found that several of these additional glomeruli were weakly labeled by the transgenic line (Figure 3-Figure Supplement 2A). This suggests that there are OSN populations in which *Orco* is expressed either at low levels or in few cells, which might be why this expression was previously missed. We found this to be the case with the *IrCo* knock-ins, as well (described below).

There has been some inconsistency in the literature as to which glomeruli are innervated by Ir8a-expressing OSNs. For example, Silbering et al., 2011 note that their *Ir8a-Gal4* line labels approximately ten glomeruli, six of which are identified (DL2, DP1l, VL2a, VL2p, DP1m, DC4). An *Ir8a-Gal4* line generated by Ai et al., 2013 also labels about ten glomeruli, only two of which are identified (DC4 and DP1m) and which correspond to two glomeruli in Silbering et al., 2011. Finally, Min et al., 2013 identify three additional glomeruli innervated by an *Ir8a-Gal4* line (VM1, VM4, and VC5) but not reported in the other two papers. DL2 was later subdivided into two glomeruli (Prieto-Godino et al., 2017), bringing the total number of identified Ir8a+ glomeruli to ten. However, we found that *Ir8a-T2A-QF2* consistently labels twice as many glomeruli as previously reported. These additional glomeruli include an Ir25a+ glomerulus (VL1, Figure 3B), numerous Orco+ glomeruli (such as VA3 and VA5), and an Orco+/Ir76b+ glomerulus (VC3) (see Figure 3-Source Data 1 for a full list of new glomeruli and Figure 3-Source Data 2 for outlined examples). Some of these additional glomeruli are weakly labeled by an *Ir8a-Gal4* line (Figure 3-Figure Supplement 2B), but this innervation is only apparent when examined with a strong reporter.

Of the four chemosensory co-receptor genes, the previously reported expression of *Ir76b* is the narrowest, with only four identified glomeruli (VM1, VM4, VC3, VC5) (Silbering et al., 2011). The *Ir76b-T2A-QF2* knock-in labels more than three times this number, including several Orco+ glomeruli (such as DC3 and VC4), most Ir8a+ glomeruli (including DP1l), and one additional Ir25a+ glomerulus (VL1) (Figure 3C). As with *Orco* and *Ir8a*, some but not all of these glomeruli can be identified by crossing the transgenic *Ir76b-Gal4* line to a strong reporter (Figure 3-Figure Supplement 2C). However, the *Ir76b-Gal4* line labels additional glomeruli not seen in the knock-in (Figure 3-Figure Supplement 2C, Orco+ cluster). In total, the *Ir76b-T2A-QF2* knock-in labels 15 glomeruli consistently and two variably (Figure 3-Source Data 1 and 2).

*Ir25a-T2A-QF2* innervation of the AL was the most expanded compared to what has previously been reported. In addition to the novel expression we identified in the palps (Figure 2H), we found that the *Ir25a* knock-in innervates many Orco+ glomeruli receiving inputs from the antennae (Figure 3D). The extensive, dense innervation of the AL by Ir25a+ processes made identification of individual glomeruli difficult and necessitated further experiments to fully characterize this expression pattern (described in greater detail below). While it was previously reported that the transgenic *Ir25a-Gal4* line labels only a subset of Ir25a+ neurons (compared to anti-Ir25a antibody staining), it was assumed that neurons not captured by the transgenic line would reside in coeloconic sensilla, the arista or sacculus (the original locations for all IR+ OSNs) (Abuin et al., 2011). When we crossed *Ir25a-Gal4* to a strong reporter, we found labeling of a few Orco+ glomeruli (Figure 3-Figure Supplement 2D), but this was a small fraction of those labeled by the knock-in. To further examine *Ir25a* expression and the potential co-expression of multiple co-receptors in greater detail, we employed a combination of approaches including single nucleus RNAseq (snRNAseq), immunohistochemistry, and optogenetics.

### Confirmation of Co-Receptor Co-Expression

The innervation of the same glomeruli by multiple co-receptor knock-in lines challenges the previous view of segregated chemosensory receptor expression in *Drosophila* and suggests two possible explanations: either the same olfactory neurons express multiple co-receptors (co-expression), or different populations of olfactory neurons expressing different receptors converge upon the same glomeruli (co-convergence). These scenarios are not necessarily mutually exclusive. To examine these possibilities in a comprehensive, unbiased way, we analyzed snRNAseq data from adult fly antennae (McLaughlin et al., 2021). Figure 4A shows the expression levels of the four co-receptor genes in 20 transcriptomic clusters (tSNE plots (Van der Maaten and Hinton, 2008), top row) which were mapped to 24 glomerular targets in the brain (AL maps, bottom row). The proportion of cells in each cluster expressing the given co-receptor gene is indicated by the opacity of the glomerular fill color, normalized to maximum expression for that gene (see Methods and Figure 4-Source Data 1 for details on expression normalization). The OSN classes to which these clusters map include Orco+ neurons (Figure 4A right column, teal), Ir25a+ neurons (Figure 4A right column, purple), Ir8a+ neurons (Figure 4A right column, pink), and GR+ neurons (Figure 4A right column, dark blue). They also include example OSNs from all sensillar types (basiconic, intermediate, trichoid, coeloconic) as well as from the arista and sacculus. The snRNAseq analyses confirmed expanded expression of all four co-receptor genes into OSN classes not traditionally assigned to them. For example, *Orco* and *Ir25a* are expressed in cluster 1, which maps to the V glomerulus (Gr21a+/Gr63a+). Similarly, *Ir8a* and *Ir76b* are expressed in cluster 19 (VL1 glomerulus, Ir25a+), and *Ir25a* is expressed in multiple Orco+ clusters (such as 15/VA2, 16/DL3, and 8/DC1).

**Figure 4.**
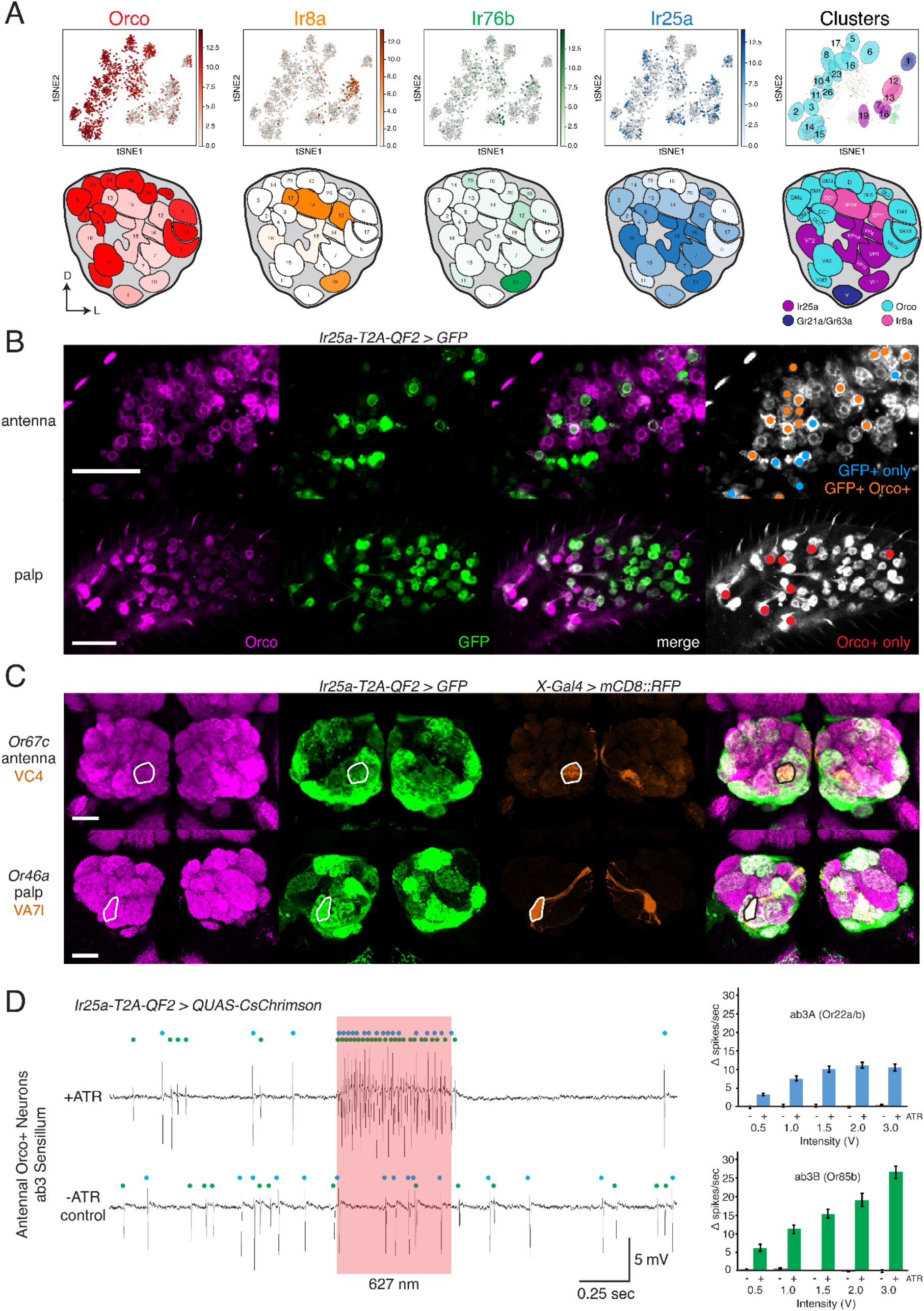
Confirmation of Co-Receptor Co-Expression. **A.** snRNAseq of adult fly antennae (McLaughlin et al., 2021) confirms expanded expression of olfactory co-receptors. Top: tSNE plots show expression of each co-receptor in 20 decoded OSN clusters. Bottom: clusters were mapped to 24 glomeruli. Opacity of fill in each glomerulus indicates the proportion of cells in that cluster expressing the given co-receptor, normalized to total expression for that co-receptor gene (see Figure 4-Source Data 1). Right column: clusters color-coded according to original chemoreceptor gene family. Compass: D = dorsal, L = lateral. **B.** Anti-Orco antibody staining in antennal cryosections (top) and whole-mount palps (bottom) confirms co-expression of Orco and Ir25a in the periphery (genotype: *Ir25a-T2A-QF2 > GFP*). Right panels show cells pseudo-colored grey with specific single- or double-labeled cells indicated by colored cell markers (GFP+ only in blue, GFP+ Orco+ in orange, Orco+ only in red). **C.** Co-labeling experiments with various transgenic *Gal4* lines driving mCD8::RFP (orange) and the *Ir25a-T2A-QF2* knock-in driving GFP (green). *Ir25a-T2A-QF2* labels glomeruli innervated by both antennal (top) and palpal (bottom) OSNs. **D.** Verification of *Ir25a* expression in antennal ab3 sensilla using optogenetics. SSR recordings from ab3 Orco+ neurons in *Ir25a-T2A-QF2* > *QUAS-CsChrimson* flies. Representative traces from ab3 using 1.5V of 627 nm LED light (red box) to activate *CsChrimson*. Bottom trace is control animal which was not fed the required all-trans retinal cofactor (-ATR). Spikes from the ab3A and ab3B neurons are indicated by blue and green dots, respectively. Right: Quantification of neuronal activity in response to light at various LED intensities (N = 7 – 12). These optogenetic experiments support *Ir25a* expression in both ab3A neurons (*Or22a/b*, top; corresponding to DM2 glomerulus) and ab3B neurons (*Or85b*, bottom; corresponding to VM5d glomerulus). Scale bars = 25 µm. See also Figure 4-Figure Supplement 1, Table 3, and Figure 4-Source Data 1 – 3.

The snRNAseq analyses confirm transcript co-expression in olfactory neurons in the periphery. To demonstrate protein co-expression in OSNs, we performed anti-Orco antibody staining on *Ir25a-T2A-QF2 > GFP* antennae and palps (Figure 4B). In the antennae, we found examples of Orco+ GFP+ double-labeled cells, as well as many cells that were either GFP+ or Orco+ (Figure 4B, top right panel). Interestingly, in the palps the vast majority of cells were double-labeled. We found a small population of palpal neurons which were only Orco+, and no neurons that were only GFP+ (Figure 4B, bottom right panel). These results are consistent with our anti-Ir25a staining experiments in the palps (Figure 2I), which showed that most of the ∼120 palpal OSNs express Ir25a protein.

The snRNAseq data from the antennae and peripheral immunohistochemical experiments in the palps helped to identify some of the novel OSN populations expressing *Ir25a*. We extended these analyses with co-labeling experiments in which we combined transgenic *OrX-*, *IrX-*, or *GrX-Gal4* lines labeling individual glomeruli with the *Ir25a* knock-in to verify the glomerular identity of Ir25a+ axonal targets in the AL. Two examples are shown in Figure 4C (one antennal and one palpal OSN population), and the full list of OSN classes checked can be found in Figure 4-Source Data 2.

For some OSN classes not included in the snRNAseq dataset for which co-labeling experiments yielded ambiguous results, we employed an optogenetic approach. We used the *Ir25a-T2A-QF2* knock-in to drive expression of *QUAS-CsChrimson*, a red-shifted channelrhodopsin (Klapoetke et al., 2014), and performed single sensillum recordings (SSR) from sensilla previously known to house only Orco+ neurons. If these neurons do express *Ir25a*, then stimulation with red light should induce neuronal firing. We recorded from ab3 sensilla, which have two olfactory neurons (A and B; indicated with blue and green dots, respectively, in Figure 4D). Ab3A neurons innervate DM2 and ab3B neurons innervate VM5d. Both neurons responded to pulses of 627 nm light at various intensities in a dose-dependent manner, confirming *Ir25a* expression in these neurons. No light-induced responses were found in control flies not fed all-trans retinal (-ATR), a necessary co-factor for channelrhodopsin function (see Methods). We used similar optogenetic experiments to examine *Ir25a* expression in OSN classes innervating DM4 (ab2A, Or59b+) and DM5 (ab2B, Or85a/Or33b+) (Figure 4-Figure Supplement 1A-B), as well as D (ab9A, Or69aA/aB+) and VA3 (ab9B, Or67b+) (Figure 4-Figure Supplement 1C-D). These experiments indicated that *Ir25a* is expressed in ab2A (DM4) and ab9B (VA3) neurons, but not ab2B (DM5) or ab9A (D) neurons (see also Figure 4-Source Data 2 and 3). Results of these experiments are summarized in Table 3.

### Identification of New OSN Classes

The co-receptor knock-ins allowed us to analyze the olfactory neuron innervation patterns for all antennal lobe glomeruli. Interestingly, the *Ir8a-T2A-QF2* and *Ir25a-T2A-QF2* knock-ins strongly labeled a previously uncharacterized posterior region of the AL. By performing a co-labeling experiment with *Ir41a-Gal4*, which labels the VC5 glomerulus, we narrowed down the anatomical location of this region and ruled out VC5 as the target of these axons (Figure 5A). While both knock-ins clearly labeled VC5, they also labeled a region lateral and slightly posterior to it (Figure 5A, outline). We performed additional co-labeling experiments with *Ir8a-T2A-QF2* and various *Gal4* lines labeling all known posterior glomeruli to confirm that this AL region did not match the innervation regions for other previously described OSN populations (Figure 5-Figure Supplement 1). We recognized that this novel innervation pattern appeared similar to a portion of the recently identified Rh50+ ammonia-sensing olfactory neurons (Vulpe et al., 2021a). Co-labeling experiments with *Rh50-Gal4* and *Ir8a-T2A-QF2* confirmed that they indeed partially overlapped (Figure 5B). We determined that these Rh50+ olfactory neurons mapped to a portion of the VM6 glomerulus, with the strongly Ir8a+ region innervating the ‘horn’ of this glomerulus. The difference in innervation patterns between Ir8a+ and Rh50+ neurons in this AL region suggested at least two different subdivisions or OSN populations within this VM6 glomerulus. In fact, in between the main body of VM6 and the Ir8a+ horn there appeared to be a third region (Figure 5B, horn outlined in white, other two regions outlined in blue). We designated these subdivisions VM6l, VM6m, and VM6v (for lateral, medial, and ventral). We coordinated the naming of this glomerulus with recent connectomics analyses of the entire fly antennal lobe (Schlegel et al., 2021). In this connectomics study, dendrites of olfactory projection neurons were found to innervate the entire region described here as VM6l, VM6m, and VM6v. No projection neurons were identified to innervate only a subdomain. As such, the new VM6 nomenclature reflects this unique subdivision of a glomerulus by olfactory sensory neurons but not second order projection neurons.

**Figure 5.**
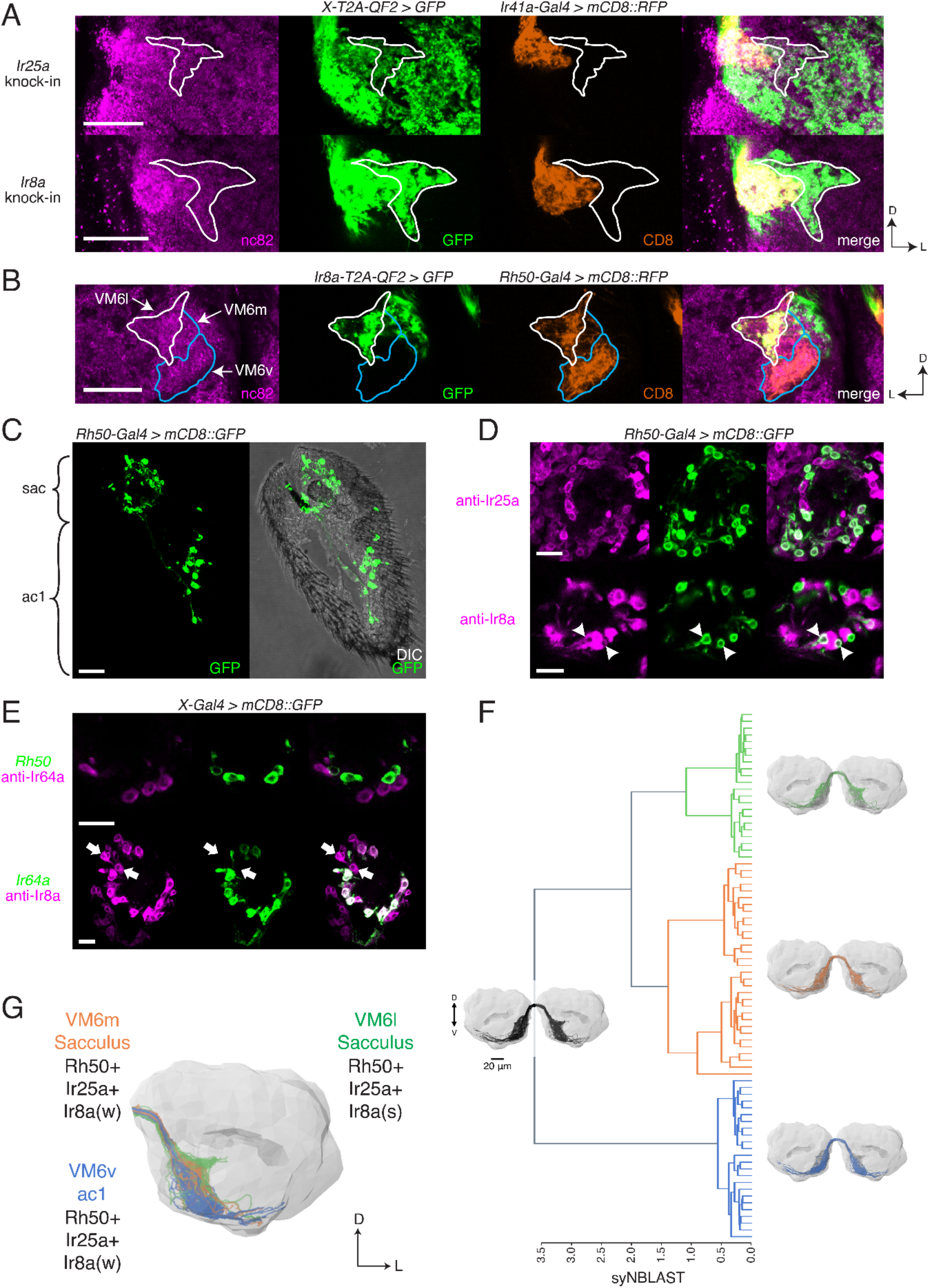
Identification of New OSN Classes. **A.** Co-labeling experiments with *Ir41a-Gal4* show that both *Ir25a-T2A-QF2* and *Ir8a-T2A-QF2* label the VC5 glomerulus (orange), and also a previously unidentified AL region (outline). **B.** The new innervation pattern corresponds to the ‘horn’ (white outline) of the VM6 glomerulus labeled by Rh50+ neurons (orange). One portion of VM6 is strongly Ir8a+ (VM6l), while two other portions show little to no Ir8a expression (VM6m and VM6v, blue outlines). **C.** *Rh50-Gal4 > GFP* labels neurons in the sacculus (sac) and antennal coeloconic ac1 sensilla. **D.** In the sacculus, all Rh50+ neurons appear to be Ir25a+ (top), and a subset are Ir8a+ (bottom, arrowheads). **E.** Top: Rh50+ neurons in the sacculus do not overlap with Ir64a+ neurons. Bottom: there are two distinct populations of Ir8a+ neurons in the sacculus – those that are Ir64a+ and those that are Ir64a-(arrows). The latter likely correspond to Rh50+ neurons. **F.** EM reconstructions of VM6 OSNs in a full brain volume (Dorkenwald et al., 2020) reveal three distinct subpopulations. **G.** Model of OSN innervation of the VM6 region. VM6 can be subdivided into three OSN populations based on anatomical location in the periphery and chemoreceptor expression: VM6v (blue) OSNs originate in ac1, strongly (s) express Rh50 and Ir25a, and weakly (w) or infrequently express Ir8a; VM6m (orange) neurons originate in the sacculus and have a similar chemoreceptor expression profile to VM6v; VM6l (green) OSNs originate in the sacculus but strongly express Ir8a in addition to Rh50 and Ir25a. Compass: D = dorsal, L = lateral. Scale bars: 20 µm in (A-C) and (F), 10 µm in (D-E). See also Figure 5-Figure Supplement 1 and Tables 3 and 4.

**Table 4.**
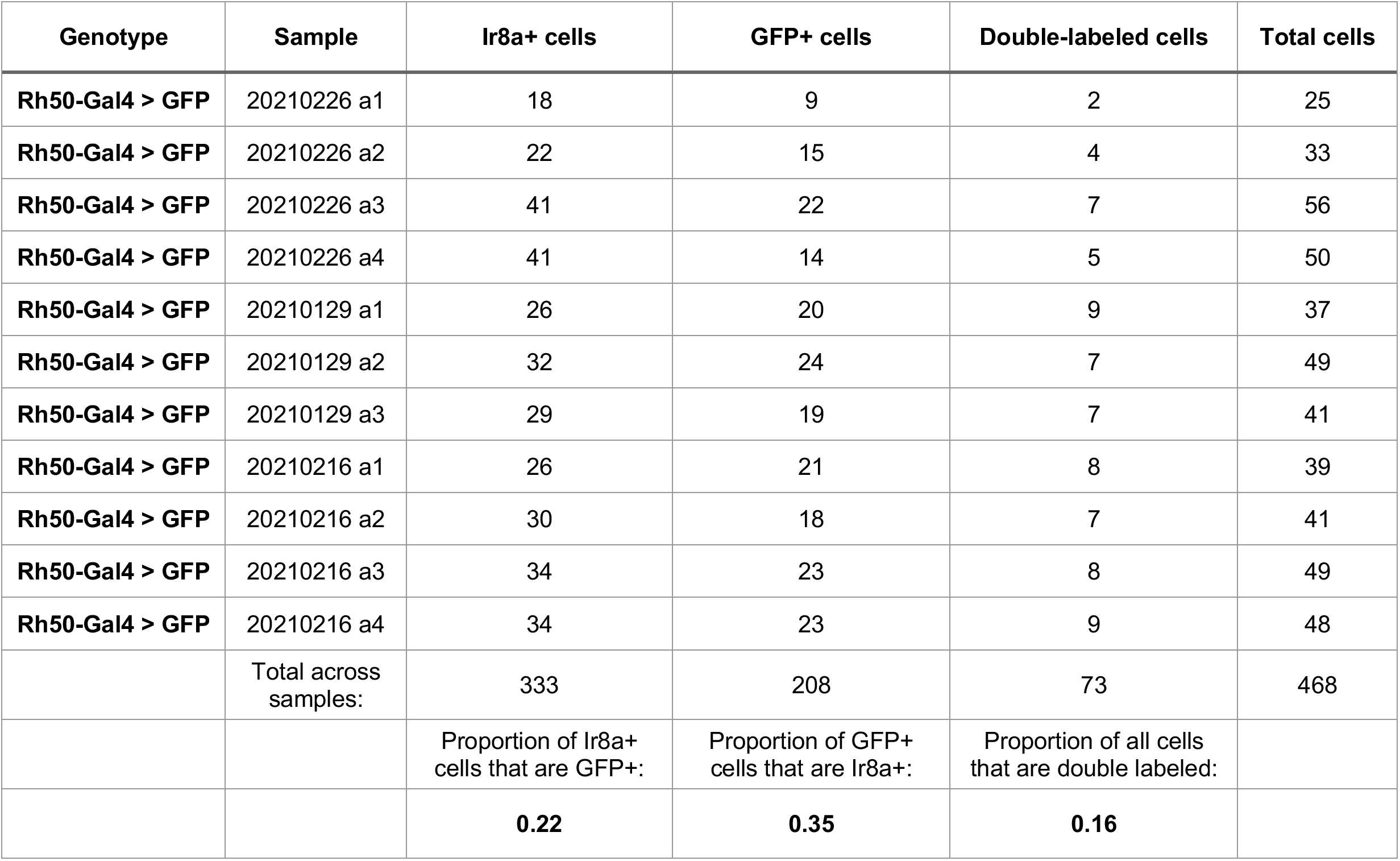
Co-expression of Rh50 and Ir8a in the Sacculus. Related to Figure 5. Antennal cryosections of *Rh50-Gal4>GFP* flies were stained with an anti-Ir8a antibody and the overlap of Ir8a+ and GFP+ cells was quantified in the sacculus. 22% of Ir8a+ cells expressed Rh50, 35% of Rh50+ cells expressed Ir8a, and 16% of all cells were double labeled. N = 11.

| Genotype | Sample | Ir8a+ cells | GFP+ cells | Double-labeled cells | Total cells |
| --- | --- | --- | --- | --- | --- |
| **Rh50-Gal4 > GFP** | 20210226 a1 | 18 | 9 | 2 | 25 |
| **Rh50-Gal4 > GFP** | 20210226 a2 | 22 | 15 | 4 | 33 |
| **Rh50-Gal4 > GFP** | 20210226 a3 | 41 | 22 | 7 | 56 |
| **Rh50-Gal4 > GFP** | 20210226 a4 | 41 | 14 | 5 | 50 |
| **Rh50-Gal4 > GFP** | 20210129 a1 | 26 | 20 | 9 | 37 |
| **Rh50-Gal4 > GFP** | 20210129 a2 | 32 | 24 | 7 | 49 |
| **Rh50-Gal4 > GFP** | 20210129 a3 | 29 | 19 | 7 | 41 |
| **Rh50-Gal4 > GFP** | 20210216 a1 | 26 | 21 | 8 | 39 |
| **Rh50-Gal4 > GFP** | 20210216 a2 | 30 | 18 | 7 | 41 |
| **Rh50-Gal4 > GFP** | 20210216 a3 | 34 | 23 | 8 | 49 |
| **Rh50-Gal4 > GFP** | 20210216 a4 | 34 | 23 | 9 | 48 |
|  | Total across samples: | 333 | 208 | 73 | 468 |
|  |  | Proportion of Ir8a+ cells that are GFP+: | Proportion of GFP+ cells that are Ir8a+: | Proportion of all cells that are double labeled: |  |
|  |  | **0.22** | **0.35** | **0.16** |  |

We sought to determine the identity of the olfactory neurons that might be innervating these three VM6 subdivisions. Rh50+ neurons can be found in two regions of the antenna: ac1 coeloconic sensilla and the sacculus (Figure 5C) (Vulpe et al., 2021a). The shape of the VM6v subdomain most closely matches the glomerulus described as VM6 by previous groups (e.g. Couto et al., 2005; Endo et al., 2007), which had been suggested to be innervated by coeloconic sensilla (Chai et al., 2019; Li et al., 2016). In addition, antibody staining had previously shown that Rh50+ ac1 neurons broadly co-express Ir25a but generally not Ir8a (Vulpe et al., 2021a). This suggested that the other VM6 subdomains might be innervated by the Rh50+ sacculus olfactory neurons. Antibody staining in *Rh50-Gal4>GFP* antennae confirmed co-expression with both Ir25a protein (broad overlap) and Ir8a protein (narrow overlap) in the third chamber of the sacculus (Figure 5D; quantified in Table 4). Most sacculus neurons appear to be Ir25a+, and in contrast to the *Ir8a* knock-in, the three VM6 subdivisions are all strongly innervated by the *Ir25a* knock-in (Figure 5A). Two previously described OSN populations in the third chamber of the sacculus had been characterized to express *Ir8a* along with *Ir64a* and innervate the DP1m and DC4 glomeruli (Ai et al., 2013; Ai et al., 2010). To demonstrate that the Rh50+ Ir8a+ sacculus neurons represented a distinct olfactory neuron population, we performed immunohistochemistry experiments in *Rh50-Gal4>GFP* antennae with an anti-Ir64a antibody (Figure 5E, top), and in *Ir64a-Gal4>GFP* antennae with an anti-Ir8a antibody (Figure 5E, bottom). These experiments confirmed a new, distinct population of Ir8a+ Ir64a-cells in the sacculus.

The VM6l olfactory projections are difficult to identify in the hemibrain connectome (Scheffer et al., 2020) due to the medial truncation of the AL in that dataset (see Schlegel et al., 2021 for additional details). Here, we used FlyWire (Dorkenwald et al., 2020), a recent segmentation of a full adult fly brain (FAFB) (Zheng et al., 2018), to reconstruct the VM6 OSN projections in both left and right AL. Synapse-based hierarchical clustering (syNBLAST) (Buhmann et al., 2021) of the VM6 OSNs demonstrated the anatomical segregation into three distinct subpopulations: VM6l, VM6m and VM6v (Figure 5F). This subdivision was subsequently confirmed in a re-analysis of the VM6 glomerulus in the hemibrain dataset (Schlegel et al., 2021). Olfactory neurons innervating VM6l were strongly Ir8a+, while olfactory neurons innervating VM6m and VM6v were weakly and sparsely Ir8a+ (see Figure 3-Source Data 2, page 3). This pattern may be due to *Ir8a* expression in only one or a few cells. Interestingly, this sparse Ir8a+ expression in olfactory neurons targeting VM6m and VM6v may be sexually dimorphic: we found that male brains had stronger and more frequent Ir8a+ innervation in these two glomeruli compared to female brains (see also Figure 3-Source Data 1).

Based on the EM reconstructions, genetic AL analyses, and peripheral staining experiments, we propose a model of the anatomical locations and molecular identities of the olfactory neurons innervating the VM6 subdivisions (Figure 5F). All VM6 subdivisions broadly express *Rh50* and *Ir25a*; the VM6v OSNs are housed in ac1 sensilla and express *Ir8a* either weakly or only in a small subset of neurons; both the VM6m and VM6l OSNs are found in the sacculus and can be distinguished by their levels or extent of *Ir8a* expression, with VM6l neurons being strongly Ir8a+. Because all three VM6 subdivisions share the same downstream projection neurons, this AL region has been classified as a single glomerulus (Schlegel et al., 2021). We maintain this convention here, for a total of 58 AL glomeruli. It is possible that this number may need to be re-evaluated in the future, and the three VM6 subdivisions reconsidered as *bona fide* separate glomeruli (bringing the OSN glomerular total to 60). Such a separation might be warranted if it is found that these OSN populations express different tuning receptors, and those receptors respond to different odorants.

Table 3 summarizes the chemosensory receptor expression patterns for all four co-receptor knock-in lines across all OSNs, sensillar types, and glomeruli. For clarity, this summary considers the newly identified OSN populations described here separately. We find that *Orco-T2A-QF2* consistently labels 45 total glomeruli out of 58 (7 more than previously reported); *Ir8a-T2A-QF2* consistently labels 18 glomeruli (8 more than previously identified); *Ir76b-T2A-QF2* consistently labels 15 glomeruli (11 more than previously identified); and *Ir25a-T2A-QF2* consistently labels 51 glomeruli (39 more than previously identified).

### Co-receptor Contributions to Olfactory Neuron Physiology

How might the broad, combinatorial co-expression of various chemosensory families affect olfactory neuron function? To begin to address this question we examined olfactory responses in neuronal populations co-expressing just two of the four chemosensory receptor families (*Orco* and *Ir25a*). We chose to test eight OSN classes previously assigned to the Orco+ domain that we found to have strong or intermediate *Ir25a* expression – two in the antennae, and six in the maxillary palps. The two antennal OSN classes are found in the same ab3 sensillum (ab3A, Or22a/b+, DM2 glomerulus; and ab3B, Or85b+, VM5d glomerulus). The six palpal OSN classes represent the entire olfactory neuron population of the maxillary palps (pb1A, Or42a+, VM7d; pb1B, Or71a+, VC2; pb2A, Or33c/Or85e+, VC1; pb2B, Or46a+, VA7l; pb3A, Or59c+, VM7v; pb3B, Or85d+, VA4). In both the antennae and the palps, we compared the olfactory responses of OSNs to a panel of 13 odorants in three genotypes: *wildtype*, *Ir25a^2^* mutant, and *Orco^2^* mutant flies. This panel included odorants typically detected by *ORs*, such as esters and aromatics, and odorants typically detected by *IRs*, such as acids and amines (Silbering et al., 2011). In the previously accepted view of olfaction in *Drosophila*, Orco+ neurons express only Orco/OrX receptors, and all olfactory responses in the neurons can be attributed to these receptors. Thus, in an *Ir25a^2^* mutant background, there should be no difference in olfactory responses from *wildtype* if either a) *Ir25a* is not expressed in these neurons, or b) *Ir25a* is expressed, but is not playing a functional role in these neurons. In an *Orco^2^* mutant background, there would be no trafficking of Orco/OrX receptors to the dendritic membrane, and no formation of functional ion channels (Benton et al., 2006; Larsson et al., 2004). Thus, in the traditional view of insect olfaction, *Orco^2^* mutant neurons should have no odor-evoked activity. However, in the new co-receptor co-expression model of olfaction, if *Ir25a* is contributing to olfactory responses in Orco+ neurons, then mutating this co-receptor might affect the response profiles of these neurons. Similarly, *Orco^2^* mutant neurons that co-express *Ir25a* might retain some odor-evoked activity.

We first examined olfactory responses in palp basiconic sensilla. In the palps, three types of basiconic sensilla (pb1, pb2, and pb3) contain two neurons each (A and B) (Figure 6A), for a total of six OSN classes (Couto et al., 2005; de Bruyne et al., 1999; Fishilevich and Vosshall, 2005; Goldman et al., 2005; Ray et al., 2008; Ray et al., 2007). We found robust responses to several odorants in our panel in both the *wildtype* and *Ir25a^2^* mutant flies, including odorants like 1-octen-3-ol typically considered as an OR ligand (Figure 6B), and IR ligands like pyrrolidine. Neither odor-evoked nor spontaneous activity was detected in the *Orco^2^* mutant (Figure 6B, bottom row; see also Figure 6-Figure Supplement 1A). This was true of all sensilla tested in the palps. The SSR experiments in Figure 6A-D were performed at 4 – 8 DPE. We recently discovered that neurodegeneration of *Orco^2^* mutant olfactory neurons occurs in the palps by ∼6 DPE (Task and Potter, 2021), which could potentially confound our interpretation. We repeated the experiments in young (1 – 3 DPE) flies but similarly detected neither odor-evoked activity nor spontaneous activity in mutant palpal neurons (Figure 6-Figure Supplement 1B). There was also no spontaneous or odor-evoked activity in an *Ir25a^2^*; *Orco^2^* double mutant (Figure 6-Figure Supplement 1C). This suggests one of three possibilities: first, *Orco^2^* mutant neurons in the palps could already be dysfunctional at this early stage, despite not yet showing cell loss, and *Ir25a*-dependent activity is not sufficient to maintain either baseline or stimulus-induced activity; second, *Ir25a* function may be *Orco*-dependent in these cells, or act downstream of *Orco*, such that loss of *Orco* function affects *Ir25a* function; third, we did not stimulate neurons with an *Ir25a*-dependent odorant. The latter possibility would not, however, explain why there is no spontaneous activity in these cells. Future experiments will be needed to address these possibilities. Given the lack of neuronal activity in the *Orco^2^* mutant, we focused subsequent analyses in the palps on the two other genotypes: *wildtype* and *Ir25a^2^*.

**Figure 6.**
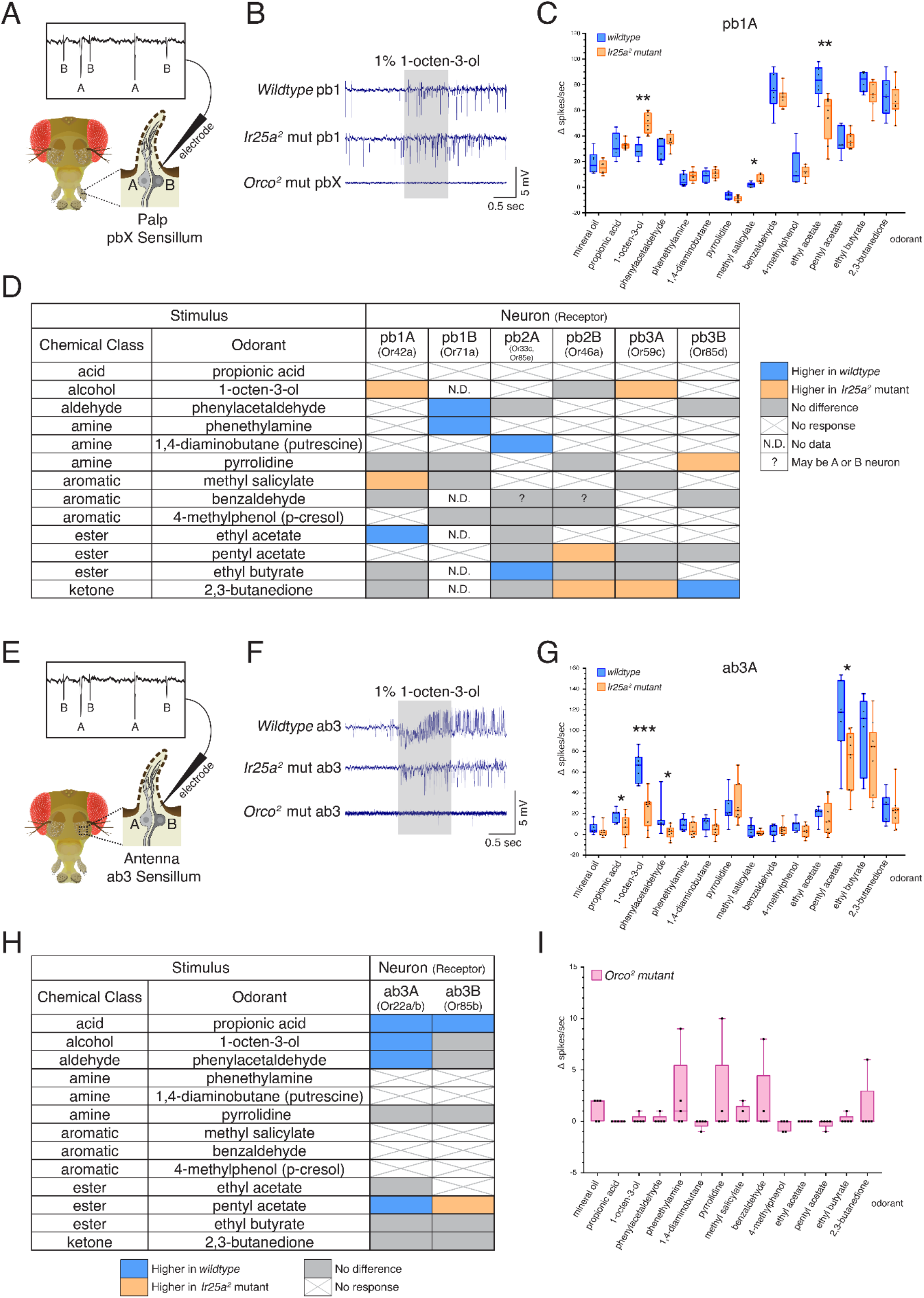
Co-Receptor Contributions to Olfactory Neuron Physiology. **A-I.** SSR experiments were performed in three genetic backgrounds: *wildtype*, *Ir25a^2^* mutant, and *Orco^2^* mutant flies. A panel of 13 odorants was tested. In all box plots, * *p* < 0.05, ** *p* < 0.01, and *** *p* < 0.001. **A.** Cartoon of a fly head, zooming in on a single sensillum in the palp. Each palpal sensillum (pbX) contains two neurons, A and B. An electrode is inserted into the sensillum, and neuronal activity is recorded in response to odorants. Activity of the A and B neurons can be distinguished based on their spike amplitudes (top). **B.** Representative traces from recordings in palp basiconic pb1 sensilla in the three genotypes in response to 1% 1-octen-3-ol. Sensilla were identified based on responses to reference odorants (de Bruyne et al., 1999). The *Orco^2^* mutant did not exhibit odor-evoked activity nor spontaneous activity making it difficult to determine the identity of the recorded sensillum. *Orco^2^* mutant sensilla are thus denoted pbX. **C.** Quantification of responses to the panel of odorants in *wildtype* (blue; N = 5 – 9 flies) and *Ir25a^2^* mutant (orange; N = 6 – 10 flies) pb1A neurons. Responses were higher in the *Ir25a^2^* mutant than in the *wildtype* for 1-octen-3-ol and methyl salicylate, and lower in the *Ir25a^2^* mutant for ethyl acetate. Mann-Whitney U tests indicated these differences were statistically significant: 1-octen-3-ol: *Mdn_Ir25amut_* = 50, *Mdn_wildtype_* = 28, *U*(*N_Ir25amut_* = 8, *N_wildtype_* = 5) = 0, *p* = 0.0016; methyl salicylate: *Mdn_Ir25amut_* = 5, *Mdn_wildtype_* = 2, *U*(*N_Ir25amut_* = 7, *N_wildtype_* = 5) = 3, *p* = 0.0177; ethyl acetate: *Mdn_Ir25amut_* = 63.5, *Mdn_wildtype_* = 83.5, *U*(*N_Ir25amut_* = 8, *N_wildtype_* = 6) = 4, *p* = 0.008. **D.** Summary of differences in responses across all six neuron classes in the palps between *wildtype* and *Ir25a^2^* mutant flies. Comparisons were made using Mann-Whitney *U* tests. Orange indicates higher response in *Ir25a^2^* mutant, blue indicates higher response in *wildtype*. Grey is no difference between genotypes, X indicates no response to the given stimulus, and N.D. is no data (strong A neuron response obscured B neuron spikes preventing quantification). In the *wildtype*, for one sensillum-odorant combination (pb2 and benzaldehyde), it could not be distinguished if responses arose from the A or B neuron or both (indicated by a question mark). **E.** Fly head cartoon, zooming in on a single sensillum in the antenna. We recorded from antennal ab3 sensilla, each of which contains two neurons, A and B. As in the palps, responses from these neurons can be distinguished based upon their spike amplitude (top). **F.** Representative traces from recordings in antennal basiconic ab3 sensilla in the three genotypes in response to 1% 1-octen-3-ol. In *Orco^2^* mutant ab3 sensilla spontaneous activity was observed, but there was no significant odor-evoked activity. *Wildtype* N = 7 sensilla from 5 flies; *Ir25a^2^* mutant N = 10 sensilla from 5 flies. **G.** Quantification of responses in *wildtype* (blue; N = 7) and *Ir25a^2^* mutant (orange; N = 9) ab3A neurons. Responses were significantly higher in *wildtype* compared to *Ir25a^2^* mutant ab3A neurons for four odorants (Mann-Whitney *U* results in parentheses; all *N_wildtype_* = 7 and *N_Ir25amut_* = 9): propionic acid (*Mdn_wildtype_* = 21, *Mdn_Ir25amut_* = 7, *U* = 12.5, *p* = 0.0441); 1-octen-3-ol (*Mdn_wildtype_* = 67, *Mdn_Ir25amut_* = 29, *U* = 1.5, *p* = 0.0004); phenylacetaldehyde (*Mdn_wildtype_* = 10, *Mdn_Ir25amut_* = 3, *U* = 9, *p* = 0.015); and pentyl acetate (*Mdn_wildtype_* = 118, *Mdn_Ir25amut_* = 77, *U* = 9, *p* = 0.0164). Difference between *wildtype* and *Ir25a^2^* mutant to phenylacetaldehyde is significant even with the large *wildtype* outlier removed (*p* = 0.0336). **H.** Summary of differences in responses in the two neuron classes in ab3 between *wildtype* and *Ir25a^2^* mutant flies. Comparisons were made using Mann-Whitney *U* tests. Orange indicates higher response in *Ir25a^2^* mutant, blue indicates higher response in *wildtype*, grey is no difference between genotypes, and X is no response to the given stimulus. One *Ir25a^2^* mutant fly was excluded from analyses as it had high responses to the mineral oil control (40 – 53 Δ spikes/sec), not seen in any other animal of any genotype. **I.** Weak responses in *Orco^2^* mutant flies to certain stimuli (≤ 10 Δ spikes/sec) were occasionally detected. While there were some statistically significant differences from mineral oil control (pentyl acetate *p* = 0.0109, propionic acid *p* = 0.0434, ethyl acetate *p* = 0.0434, 1,4-diaminobutane *p* = 0.0109, p-cresol *p* = 0.0021), these were not deemed biologically significant due to very small Δ spike values relative to zero. For more details, see Methods. N = 5 flies. See also Figure 6-Figure Supplements 1 and 2, and Figure 6-Source Data 1 and 2.

The response in the pb1A neuron to 1-octen-3-ol was significantly higher in the *Ir25a^2^* mutant compared to the *wildtype* (Mann-Whitney *U* test, *p* = 0.0016), as was the response to methyl salicylate (*p* = 0.0177), while the response to ethyl acetate was higher in *wildtype* (*p* = 0.008) (Figure 6C; see Figure 6-Source Data 1 for results of all statistical analyses). The differences in responses across all six OSN classes in the palps between *wildtype* and *Ir25a^2^* mutant flies are summarized in Figure 6D. In each neuron class, we found one to three odorants whose response profiles differed between the two genotypes. However, the specific stimuli eliciting different responses, and the directionality of those responses, varied. For example, 2,3-butanedione elicited higher responses in the *Ir25a^2^* mutant in both pb2B and pb3A neurons, but lower responses in the mutant (higher in the *wildtype*) in pb3B. Interestingly, when we examined a list of candidate IrX tuning receptors (Li et al., 2021) in the palps using *in situs*, we did not find expression (see Figure 6-Figure Supplement 2 and Figure 6-Source Data 2). This suggests that *Ir25a* may not be functioning as a traditional co-receptor in Orco+ olfactory neurons in the palps (an expanded role for *Ir25a* beyond co-reception has previously been suggested; see Budelli et al., 2019; Chen et al., 2015).

We next examined olfactory responses in antennal basiconic ab3 sensilla in *wildtype*, *Ir25a^2^* mutant, and *Orco^2^* mutant flies (Figure 6E – I). As in the palps, ab3 sensilla contain two neurons, A and B (Figure 6E). In contrast to the palps, *Orco^2^* mutant ab3 sensilla did occasionally show spontaneous activity (Figure 6F, bottom row; see Figure 6-Figure Supplement 1D for additional example traces). Although there are two Orco+ neurons in this sensillum, we consistently observed only a single spike amplitude in the *Orco^2^* mutant. Thus, we cannot determine at this time whether this activity arises from the A or B neuron. We occasionally observed small responses (≤ 10 Δ spikes/sec) in the *Orco^2^* mutant; however, across all flies tested, these responses were not significantly different from the mineral oil control (Figure 6I; statistical analyses can be found in Figure 6-Source Data 1). For these reasons, *Orco^2^* mutant flies were excluded from the analyses in Figure 6G and Figure 6H.

As in the palps, we found significant differences in the responses of both ab3A and ab3B neurons to some odorants between the two genotypes. Comparison of all ab3A responses between *wildtype* and *Ir25a^2^* mutant are shown in Figure 6G, and results from both the A and B neurons are summarized in Figure 6H (Mann-Whitney *U*, as in Figure 6A-C; see Figure 6-Source Data 1 for all analyses). In the ab3A neuron, the *wildtype* showed higher responses to propionic acid (*p* = 0.0441), 1-octen-3-ol (*p* = 0.0004), phenylacetaldehyde (*p* = 0.015), and pentyl acetate (*p* = 0.0164). Interestingly, two of these four odorants are typically associated with *IRs* (propionic acid and phenylacetaldehyde). In the ab3B neuron, only two odorants elicited significantly different responses between the *wildtype* and *Ir25a^2^* mutant: propionic acid (response higher in *wildtype*, as with ab3A; *p* = 0.0388), and pentyl acetate (response higher in mutant, in contrast to ab3A; *p* = 0.0385). While responses to propionic acid are small in both ab3 neurons, they are abolished in the *Ir25a^2^* mutant background (Kruskal-Wallis with uncorrected Dunn’s comparing odorant responses to mineral oil control; ab3A *p* = 0.3957; ab3B *p* = 0.5184), suggesting that propionic acid detection in ab3 may be *Ir25a*-dependent.

### The Co-Receptor Co-Expression Map of Olfaction in Drosophila

Co-receptor co-expression of insect chemosensory receptors suggests multiple receptors may influence the response properties of an olfactory neuron, as we have shown in ab3 and palpal sensilla. To aid future investigations of co-receptor co-expression signaling, we synthesized our results (Table 3) into a comprehensive new map of the AL. Figure 7 summarizes the expression patterns of all the co-receptor knock-in lines and presents a new model for chemosensory receptor expression in *Drosophila melanogaster*. In Figure 7A, the expression pattern of each knock-in line is presented separately (see also Figure 3-Source Data 1). The new AL map is updated with the recent re-classification of VP1 into three glomeruli (Marin et al., 2020) and indicates the new VM6 subdivisions. In Figure 7A, the original glomerular innervation pattern for each co-receptor is shown in green, with new innervation revealed by the *T2A-QF2* knock-in lines color coded by intensity: strongly labeled glomeruli are in orange, intermediate glomeruli in yellow, and weakly labeled glomeruli are in pink. Glomeruli labeled in <50% of brains examined are designated variable (grey), and glomeruli not labeled by the given knock-in are in white. The new VM6v, VM6m, and VM6l subdivisions are labeled with grey stripes.

**Figure 7.**
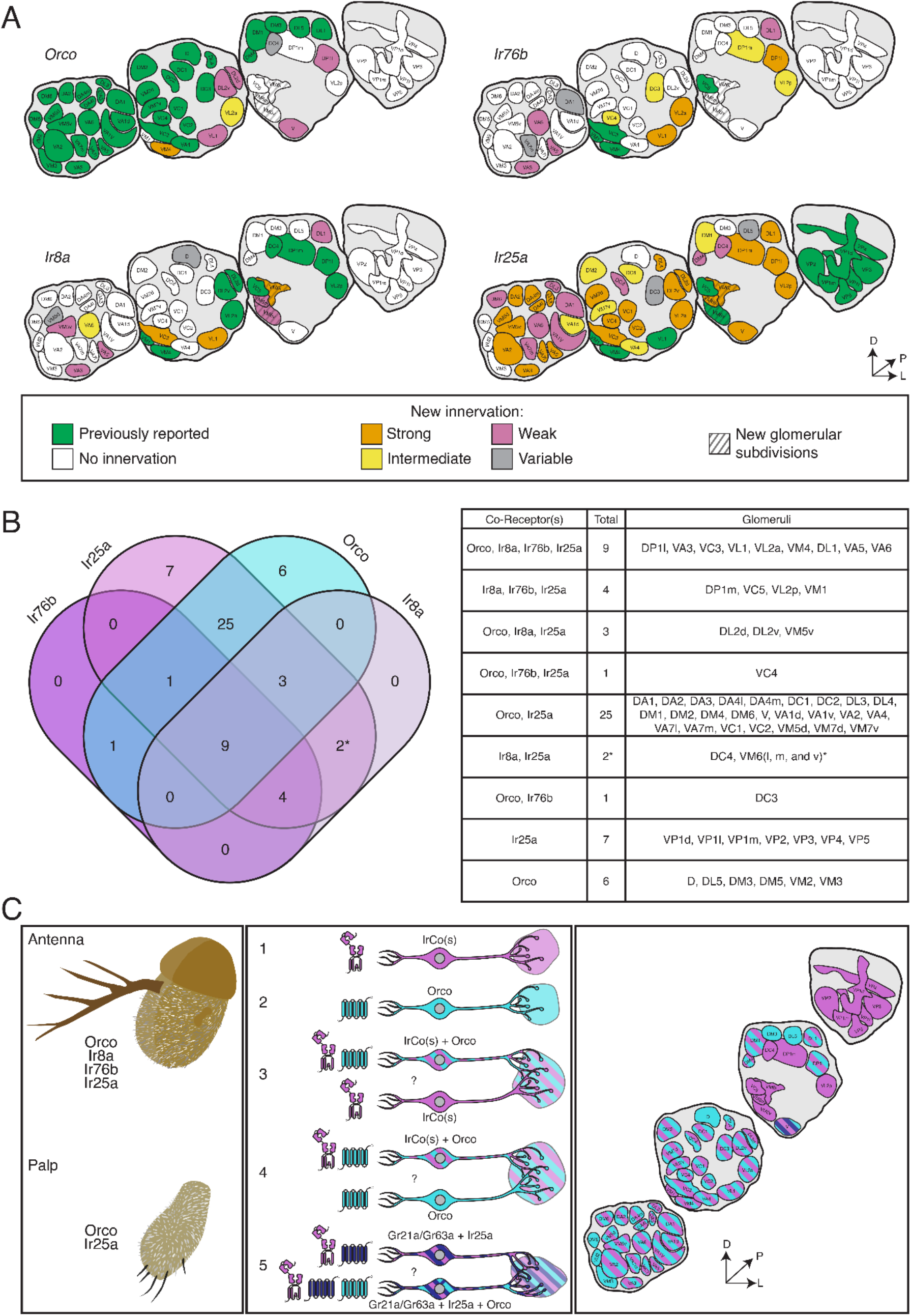
The Co-Receptor Co-Expression Map of Olfaction in *Drosophila*. **A.** Summary of AL expression for all co-receptor knock-in lines (from all brains examined in Figure 3 – 5; *Orco* N = 8, *Ir8a* N = 15, *Ir76b* N = 11, *Ir25a* N = 15). The previously reported innervation pattern for each co-receptor is shown in green; new innervation reported here is color-coded according to strength of glomerular labeling, from strong (orange), to intermediate (yellow), to weak (pink). Glomeruli labeled in <50% of brains examined for a given knock-in line are designated variable (grey); glomeruli not labeled are white. The novel VM6 glomerular subdivisions reported here are indicated by grey stripes. **B.** Overlap of chemosensory modalities in the AL. In the Venn diagram (left), *IR* co-receptors are color-coded in shades of purple, while *Orco* is in teal, as in Figure 1. Numbers indicate how many glomeruli are found in the given intersection of co-receptors out of 58 total glomeruli. Variably labeled glomeruli were excluded from these analyses. The table lists the names of the glomeruli in each section of the Venn diagram. The new glomerular subdivisions are indicated with an asterisk. **C.** New view of olfaction in *Drosophila*. Left: in the periphery, all four co-receptors are expressed in the antenna (top), while palpal neurons express *Orco* and *Ir25a* (bottom). Middle: Many different classes of OSNs express various combinations of chemosensory receptors and co-receptors. While some neurons express only *IrCos* (purple, #1) or *Orco* (teal, #2), many neurons co-express these chemoreceptors (indicated with striped fill, #3 and 4). Within the latter group, there may be OSN populations in which *IRs* are the dominant receptors, and *OR* expression is sparse (#3), and other populations where *ORs* are the primary receptors and *IR* expression is infrequent (#4). GR+ neurons (dark blue) also express *Ir25a* (#5, dark blue and purple striped fill), and some of these neurons additionally express *Orco* (#5, dark blue, purple, and teal striped fill). Question marks indicate potential instances of co-convergence of different sub-types of OSNs onto the same glomeruli. Right: a comprehensive map of the antennal lobe shows that most glomeruli are innervated by OSNs that co-express multiple chemoreceptors. Compass in (A) and (C): D = dorsal, L = lateral, P = posterior. See also Table 3 and Figure 3-Source Data 1 and 2.

In the previous model of olfaction in *Drosophila*, the Orco/OR domain primarily occupied the anterior AL, while the IR domains innervated more posterior glomeruli. While the former is still, for the most part, accurate (Figure 7A, *Orco*), the latter is not: both *Ir8a-T2A-QF2* and *Ir76b-T2A-QF2* label several more anterior glomeruli (such as VA3 or VA6), and *Ir25a-T2A-QF2* labels the majority of glomeruli throughout the anterior to posterior axis (Figure 7A, *Ir25a*). The expansion of the Ir25a+ domain is the most dramatic of the four co-receptors: previously, Ir25a+ glomeruli accounted for 21% of the AL (12/58 glomeruli) (Enjin et al., 2016; Frank et al., 2017; Marin et al., 2020; Silbering et al., 2011); the *Ir25a-T2A-QF2* knock-in consistently labels 88% of the AL (51/58 glomeruli, excluding variable). This represents a greater than fourfold expansion. Similarly, the number of Ir76b+ glomeruli increased more than threefold, from 7% of the AL (4/58 glomeruli) (Silbering et al., 2011), to 26% (15/58, excluding variable). The Ir8a+ domain has nearly doubled, from 17% of the AL originally (10/58 glomeruli) (Silbering et al., 2011) to 31% (18/58 glomeruli, excluding variable). The most modest increase in reported expression is in the Orco+ domain: from 66% of the AL (38/58 glomeruli) (Couto et al., 2005; Fishilevich and Vosshall, 2005), to 78% (45/58, excluding variable).

The expression overlap in the AL of the four co-receptor families is summarized in the Venn diagram shown in Figure 7B (excluding the variably labeled glomeruli from Figure 7A). The table at the right lists the names of the glomeruli which correspond to the sections of the Venn diagram. This analysis reveals 9 glomeruli labeled by all four knock-in lines; furthermore, it shows that the Ir8a+ and Ir76b+ domains do not have glomeruli unique to them. Most of the AL is innervated by Orco+ Ir25a+ neurons (25 glomeruli that are only Orco+ Ir25a+, plus an additional 13 that have Orco, Ir25a, and one or both other co-receptors). The Orco+ and Ir25a+ domains reveal glomeruli unique to them (6 glomeruli that are only Orco+, 7 glomeruli that are only Ir25a+). Expression analyses also reveal that Ir8a does not co-express with Orco alone or Ir76b alone.

A unified AL map organized by chemosensory gene families (ORs, IRs, and GRs) is shown in Figure 7C (right panel), and the left two panels extend this information into the periphery. Here we include the GR+ innervation of the V glomerulus. However, a knock-in line for either *Gr21a* or *Gr63a* does not currently exist; thus, it is possible these receptors (as well as other poorly characterized antennal GRs) might also be more broadly expressed than previous transgenic lines indicate (Fujii et al., 2015; Menuz et al., 2014). All four OR and IR co-receptors are expressed in the antenna, while olfactory neurons in the palps express *Orco* and *Ir25a* (Figure 7C, left panel). In the antennae, there are many different classes of OSNs expressing various combinations of chemosensory receptors and co-receptors: there are Orco+ only neurons (Figure 7C, middle panel, #2), such as those innervating the VM2 and VM3 glomeruli (teal); IrCo+ only neurons (purple), which include neurons expressing one, two, or all three *IR* co-receptors (such as VP2, VM6v, or DP1m respectively) (Figure 7C, middle panel, #1); and neurons expressing both *Orco* and *IrCo(s)* (teal and purple stripe) (Figure 7C, middle panel, #3 and 4).

The expression data suggest that different sub-populations of olfactory neurons might be targeting a shared glomerulus. Our data indicate that both Orco+ and Ir25a+ neurons innervate the GR+ V glomerulus (dark blue; see also Figure 7A). Based on the sparse innervation of the V glomerulus by the *Orco-T2A-QF2* knock-in (Figure 3A) and the lower expression levels in the snRNAseq data (Figure 4A), we hypothesize that *Orco* may be expressed in only a subset of Gr21a/Gr63a+ neurons. This contrasts with the *Ir25a-T2A-QF2* knock-in, which appears to label most Gr21a/Gr63a+ neurons. Thus, two sub-populations of neurons may be co-converging upon the same V glomerulus: neurons that express *Gr21a/Gr63a* and *Ir25a* (dark blue and purple stripes), and neurons that express *Gr21a/Gr63a, Ir25a* and *Orco* (dark blue, purple, and teal stripes) (Figure 7C, middle panel, #5). Such co-convergence has recently been shown in the olfactory system of *Aedes aegypti* mosquitoes (Younger et al., 2020). Similarly, the sparse *Orco-T2A-QF2* knock-in innervation of the DP1l glomerulus suggests that there are OSN populations expressing mostly *IRs*, but with a subset of neurons that additionally express *Orco* (Figure 7C, middle panel, #3). The converse may also be possible (Figure 7C, middle panel, #4): OSN populations which have some neurons expressing only *Orco*, and a subset expressing both *Orco* and *IrCo(s)* co-converging onto the same glomerulus. There is some evidence for this in the palps, based on our anti-Orco and anti-Ir25a antibody staining (Figure 2C and I; Figure 4B; Table 2). The snRNAseq data suggest this may also be the case in the antennae (see Figure 4-Source Data 1).

## Discussion

Here we present evidence in *Drosophila melanogaster* of widespread chemosensory co-receptor co-expression in the olfactory system, contrasting a previously accepted view of segregated, mutually exclusive olfactory domains. By generating targeted knock-ins of the four main chemosensory co-receptors (*Orco*, *Ir8a*, *Ir76b, Ir25a*), we demonstrate that all four co-receptors have broader olfactory neuron expression than previously appreciated. Co-expression of these co-receptors was found in the periphery and in the central projections to the brain. The *Ir25a* co-receptor was previously thought to be expressed only in a small subset of olfactory neurons (coeloconic, sacculus, and arista neurons), but we present evidence that it is expressed in subsets of nearly all OSN classes that innervate the majority of the fly ALs. We further demonstrate that co-expression of chemosensory co-receptors may have functional relevance for olfactory neurons, and that the *Ir25a* co-receptor may be involved in modulating the activity of some Orco+ OSNs, both in the antennae and maxillary palps. We present a new AL map which will aid future inquiries into the role that specific chemoreceptor co-expression plays in distinct OSN populations.

A model for OR/IR segregation was initially supported by developmental evidence. Two pro-neural genes specify the development of sensory structures on the antennae: *amos* and *atonal*. *Amos* mutants lack basiconic and trichoid sensilla, while *atonal* mutants do not develop coeloconic sensilla, the arista, or the sacculus (Goulding et al., 2000; Gupta and Rodrigues, 1997; Jhaveri et al., 2000a; Jhaveri et al., 2000b). It was observed that *amos* mutants lose *Orco* expression while retaining *Ir25a* expression (Benton et al., 2009). Our results generally do not conflict with this view. In the traditional segregated model of the fly olfactory system, it was presumed that *atonal* mutant antennae would show the reverse pattern: loss or *IrCo* expression but not *Orco* expression. However, our co-receptor knock-in expression results suggest that *atonal* mutants should have significant *IrCo* expression, particularly of *Ir25a*. This was indeed found to be the case in RNAseq analyses performed on *atonal* mutant antennae, which showed that both *Ir25a* and *Ir76b* expression, but not *Ir8a* expression, remained (Menuz et al., 2014). Based upon the strength of the corresponding glomerular innervations, it does appear that the previously reported Ir25a+ neurons have stronger or more consistent *Ir25a* expression, while the new Ir25a+ olfactory neurons in the antennae reported here (e.g. *OR*-expressing OSNs) are often weakly or stochastically labeled. This might also explain why *Ir25a* expression was initially overlooked in these Orco+ neural populations. The developmental pattern is different in the maxillary palps, where it is *atonal* and not *amos* which is required for development of the basiconic sensilla (Gupta and Rodrigues, 1997). Interestingly, the *Ir25a* knock-in expression corresponds well with the *atonal* developmental program across the olfactory appendages: the strongest expression of *Ir25a* is in coeloconic sensilla, but outside of these sensilla the strongest and most consistent *Ir25a* expression is in palp basiconic sensilla.

Chemosensory co-receptor co-expression may help clarify previously confounding observations regarding *Drosophila* odor coding. For example, olfactory neurons in the sacculus that express the *Ir64a* tuning receptor along with the *Ir8a* co-receptor project to two different glomeruli: DC4 and DP1m (Ai et al., 2013; Ai et al., 2010). These two glomeruli exhibit different olfactory response profiles, with DC4 being narrowly tuned to acids or protons, while DP1m is more broadly tuned. The molecular mechanism for these differences was previously unclear as it was thought these neurons expressed the same receptors. However, the co-receptor knock-in data presented here reveal that the Ir64a+ OSN sub-populations do in fact express different combinations of chemoreceptors: in addition to the *Ir8a* co-receptor, neurons innervating DC4 express *Ir25a* (and occasionally *Orco*) (see Figure 7). In contrast, neurons innervating DP1m express *Ir8a*, *Ir25a*, and *Ir76b*. Thus, perhaps it is *Ir76b* expression in DP1m-targeting Ir64a+ neurons which makes them olfactory generalists. This idea is supported by experiments in which *Ir64a* was misexpressed in neurons targeting the VM4 glomerulus, conferring a DP1m-like, rather than a DC4-like, response profile to VM4 (Ai et al., 2010). We show here that VM4-targeting neurons express *Ir8a*, in addition to *Ir25a* and *Ir76b* (as well as *Orco*): thus, molecularly, the VM4 neuron profile is more similar to DP1m (co-expressing all *IrCos*) than DC4 (co-expressing two *IrCos*), and the key distinguishing component appears to be *Ir76b*. It would be interesting to repeat such misexpression experiments in an *Ir76b^-^* mutant background to test this hypothesis.

The widespread expression of *Ir25a* in the fly olfactory system raises the possibility that it might have roles in addition to its function as an IrX co-receptor. For example, *Ir25a* has been found to play a developmental role in forming the unique structure of Cold Cells in the arista (Budelli et al., 2019). Evolutionary studies also suggest that *Ir25a* is the most ancient of all the insect chemosensory receptors (Croset et al., 2010), and the currently broad expression might reflect its previous ubiquitous role in chemosensory signaling. *Ir25a* may also be involved in downstream olfactory signal transduction or amplification, as suggested by the work presented here, and as has recently been shown for a pickpocket ion channel (Ng et al., 2019). Experiments addressing potentially expanded roles for *Ir25a* in olfactory neurons will be aided by the new chemosensory co-receptor map presented here.

While we demonstrate here that multiple chemosensory co-receptors can be co-expressed in the same olfactory neurons, it remains to be determined if this also applies to tuning (odor-binding) receptors. Previous studies suggest that OrX tuning receptors are generally limited to a single class of olfactory neurons, with the sole exception being *Or33b* (Couto et al., 2005; Fishilevich and Vosshall, 2005). However, many IrX tuning receptors remain to be characterized and could be co-expressed in multiple olfactory neurons. For example, recordings from ab1, ab3, and ab6 sensilla indicate responses to the typical IR odors 1,4-diaminobutane, ammonia, and butyric acid, respectively (de Bruyne et al., 2001), suggesting tuning IrXs may be involved. We show that *Ir25a* plays a functional role in Orco+ neurons in the antennae and palps; this suggests that these Orco+ neurons could also express as yet unidentified ligand binding IrXs. The recent release of the whole fly single-cell atlas, which includes RNAseq data from maxillary palps, allowed us to identify 6 IRs that might be expressed in palpal OSNs (*Ir40a*, *Ir51a*, *Ir60a*, *Ir62a*, *Ir76a*, *Ir93a*) (Li et al., 2021). However, *in situ* analyses for these 6 IRs in the maxillary palps did not detect a signal (see Figure 6-Figure Supplement 2 and Figure 6-Source Data 2). This suggests that *Ir25a* in the palps may be playing a role independent of its role as a co-receptor, as discussed above, or that a tuning IrX was missed by the RNAseq analyses. In antennal ab3 sensilla, we did find one odorant (propionic acid) which elicited a small response in *wildtype* neurons, and no response in *Ir25a^2^* mutant neurons. It is possible that other antennal Orco+ OSNs might utilize IR chemoreceptors for signaling. For example, the ac3B neuron, which expresses Or35a/Orco and all IR co-receptors, has recently been suggested to utilize an unidentified IrX to mediate responses to phenethylamine (Vulpe et al., 2021b). The chemoreceptor expression patterns revealed in this work will help the search for olfactory neurons that may utilize multiple chemosensory families for odor detection.

Based on the co-receptor innervation patterns in the antennal lobes, we identified a glomerulus, VM6, that is uniquely partitioned by different olfactory sensory neurons (Figure 5) (also see Schlegel et al., 2021). The co-receptor expression patterns allowed us to pinpoint the likely origin of the innervating OSNs. Since the VM6 glomerulus was labeled by both the *Ir25a-T2A-QF2* and *Ir8a-T2A-QF2* knock-in lines, the cell bodies of these neurons had to reside in the antenna; furthermore, since we did not find *Ir8a-T2A-QF2* labeling of the arista, these neurons were likely to be either in coeloconic sensilla or in the sacculus. Indeed, we determined the VM6 glomerulus to be innervated by the newly discovered Rh50+ Amt+ olfactory neurons which originate in the sacculus and ac1 sensilla (Vulpe et al., 2021a). Based on our results, Rh50+ Amt+ sacculus neurons are further subdivided into those that strongly express Ir8a, which innervate the VM6l region, and those that weakly or infrequently express Ir8a, which innervate VM6m. The functional consequences of this unusual subdivision by olfactory neurons for a glomerulus, and how this relates to the fly’s olfactory perception of ammonia or other odorants, remains to be determined. These results also highlight the value of exploring chemosensory receptor expression patterns even in the era of connectomics, as the VM6 glomerulus and its subdivisions were not easily identifiable in prior connectomics reconstructions of the entire antennal lobe (Bates et al., 2020; Scheffer et al., 2020; Schlegel et al., 2021).

Co-expression of chemosensory co-receptors might function to increase the signaling capabilities of an olfactory neuron. For example, the signaling of an Orco+ olfactory neuron may be guided primarily by the tuning OrX, and the sensitivity range extended to include odors detectable by an IrX. Co-expression might also allow synergism, such that weak activation of a co-expressed receptor could increase neuronal activity to levels sufficient to drive behavior. This might be useful in tuning behavioral response to complex odors, such that certain combinations of odors lead to stronger olfactory neuron responses. Alternatively, a co-expressed receptor inhibited by odorants might be able to attenuate a neuron’s response to odor mixtures. Co-expressed chemosensory receptors might also modulate the perceived valence of an olfactory stimulus through the same olfactory circuit or channel. For example, a recent study showed that CO_2_, which is normally aversive to flies, can be attractive in certain contexts (van Breugel et al., 2018). While the aversive response requires the known CO_2_ receptors, *Gr21a* and *Gr63a*, the attractive response was shown to be *Ir25a*-dependent. However, previous calcium imaging experiments in the AL have shown that CO_2_ primarily activates a single glomerulus, the V glomerulus, which receives inputs from the Gr21a+/Gr63a+ neurons (Suh et al., 2004). Our *Ir25a-T2A-QF2* knock-in line reveals that *Ir25a* is expressed in this OSN class. This raises the possibility that the same neurons, through a different chemoreceptor, could drive both attractive and aversive behavior in different contexts (MacWilliam et al., 2018). Co-expression of chemosensory receptors could be a mechanism to increase the functional flexibility of a numerically limited olfactory system.

*Drosophila* often serve as a model for many other insect olfactory systems, and information gleaned from *Drosophila* is frequently extrapolated to other insects (for example, DeGennaro et al., 2013; Fandino et al., 2019; Riabinina et al., 2016; Trible et al., 2017; Yan et al., 2017). The work presented here raises the possibility that other insects may also exhibit co-expression of chemosensory co-receptors. New work in *Aedes aegypti* mosquitoes suggests that this may indeed be the case: mosquito olfactory neurons can co-express Orco/IrCo/Gr receptors; in the CO_2_-sensing olfactory neuron, this co-expression functionally expands the range of activating odors (Younger et al., 2020). This suggests that co-expression of chemosensory co-receptors may be an important feature of insect olfactory neurons.

## Acknowledgements

We thank E. Marr and K. Robinson for splinkerette genetic mapping, Y.-T. Chang for cloning of *pHACK^Ir8a^* components, O. Riabinina for initial *QUAS-CsChrimson* characterization, S. Maguire for preliminary SSR experiments, P. Mohapatra for RNAseq insights, D. Baktash for help with figures, J. Konopka for advice on statistical analyses, S. Shankar for discussion of AL mapping, and R. Mann for providing lab resources. We would like to thank E.C. Marin and M. Costa for discussions regarding posterior AL glomeruli. We are also grateful to the Seung and Murthy labs for access to the flywire.ai reconstruction community. Many thanks to the following labs for sharing antibody and fly stock reagents: Leslie B. Vosshall (Rockefeller), Richard Benton (UNIL), Paul Garrity (Brandeis), Marco Gallio (Northwestern), Greg Suh (NYU/KAIST), and Thomas R. Clandinin (Stanford). We are grateful for *in situ* advice from Richard Benton/Steeve Cruchet and Margo Herre. We thank the Center for Sensory Biology Imaging Facility (NIH P30DC005211) for use of the LSM700 confocal microscope, and the Johns Hopkins School of Public Health Malaria Research Institute for use of the Olympus SZX7 microscope equipped with QImaging QIClick Cooled digital CCD camera. We thank J. Raji, S. Maguire, and J. Konopka for comments on the manuscript and members of the Potter and Menuz labs for discussion. We thank the Vosshall lab for sharing their *Aedes* findings before publication. This work was supported by a Shelanski award to C.-C. L; a Wellcome Trust Collaborative Award (203261/Z/16/Z) and an NIH BRAIN Initiative grant (1RF1MH120679-01) to G.S.X.E. Jefferis; grants from the National Institutes of Health to H.L. (R00 AG062746); grants from the National Institutes of Health to K.M. (NIGMS R35GM133209; NIDCD 1R21DC017868); grants from the Department of Defense to C.J.P. (W81XWH-17-PRMRP) and from the National Institutes of Health to C.J.P. (NIAID R01Al137078; NIDCD R01DC013070).

Portions of this work appear in Chapter 3 of D.T.’s doctoral dissertation.

## Author Contributions

**Conceptualization**: D.T. (knock-in analysis, SSR, AL maps and glomerular analyses), C-C.L. (HACK development, knock-in analysis, SSR, optogenetics), A.V. (SSR), M.B. (snRNAseq), P.S. (EM reconstruction and analyses), H.L. (snRNAseq), K.M. (SSR, IRs, palp analysis), C.J.P. (all aspects); **Methodology**: D.T. (cloning and generation of knock-ins (*Ir25a*, *Ir8a*, *Ir76b*), immunohistochemistry, SSR, statistics, AL maps), C-C.L. (HACK, genetics, cloning and generation of knock-ins (*Orco*, *Ir8a*, *Ir76b*), cloning and generation of *QUAS-CsChrimson*, SSR, optogenetics), A.V. (SSR), A.A. (SSR), S.B. (immunohistochemistry), M.B. (snRNAseq), P.S. (EM reconstruction and analyses), H.L. (snRNAseq), K.M. (SSR, IR expression), C.J.P. (all aspects); **Formal Analysis**: D.T. (knock-ins, SSR, cell counts), C-C.L. (knock-ins, SSR), A.V. (SSR), A.A. (SSR), S.B. (cell counts), M.B. (snRNAseq), P.S. (EM analyses), H.L. (snRNAseq); **Investigation**: D.T. (knock-in generation and validation (*Ir25a*, *Ir8a*, *Ir76b*), immunohistochemistry, AL analyses), C-C.L. (HACK development, knock-in generation and validation (*Orco*, *Ir8a*, *Ir76b*), *QUAS-CsChrimson* generation and validation, optogenetics, SSR – palp, ab2), A.V. (SSR – coeloconic), A.A. (SSR – ab3), S.B. (immunohistochemistry), M.B. (snRNAseq), P.S. (EM reconstruction and analyses), H.L. (snRNAseq); **Writing, Original Draft**: D.T. and C.J.P.; **Writing, Review & Editing**: All Authors; **Visualization**: D.T. (olfactory system/fly cartoons, AL maps, microscopy images, SSR, Venn diagram, tables), C-C.L. (genetics cartoons, crossing schemes, SSR, optogenetics), M.B. (snRNAseq), P.S. (EM traces and dendrograms), H.L. (snRNAseq), C.J.P. (all aspects); **Supervision**: G.S.X.E.J., K.M. and C.J.P.; **Funding Acquisition**: C.J.P.

## Materials and Methods

### Key Resources Table

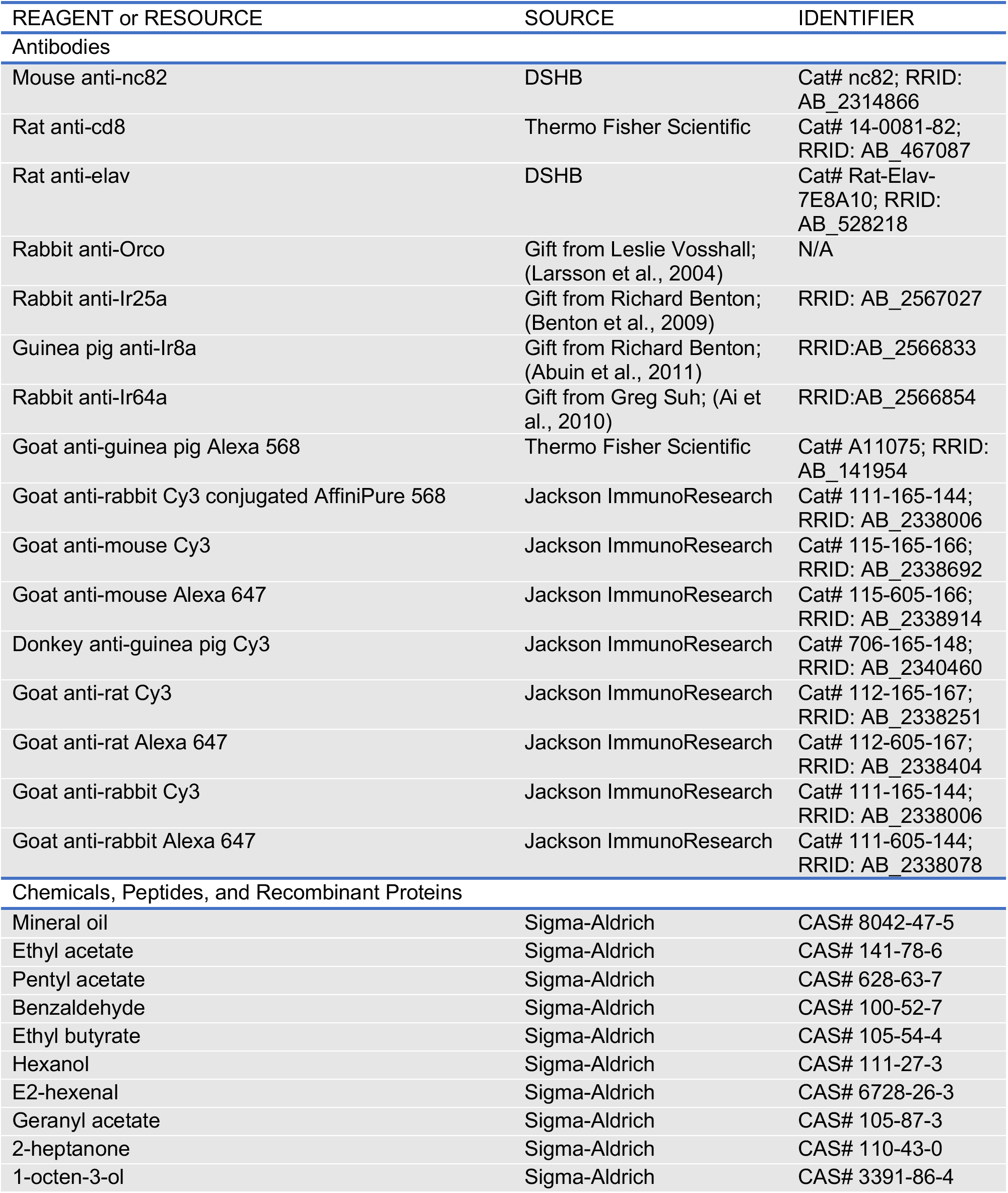
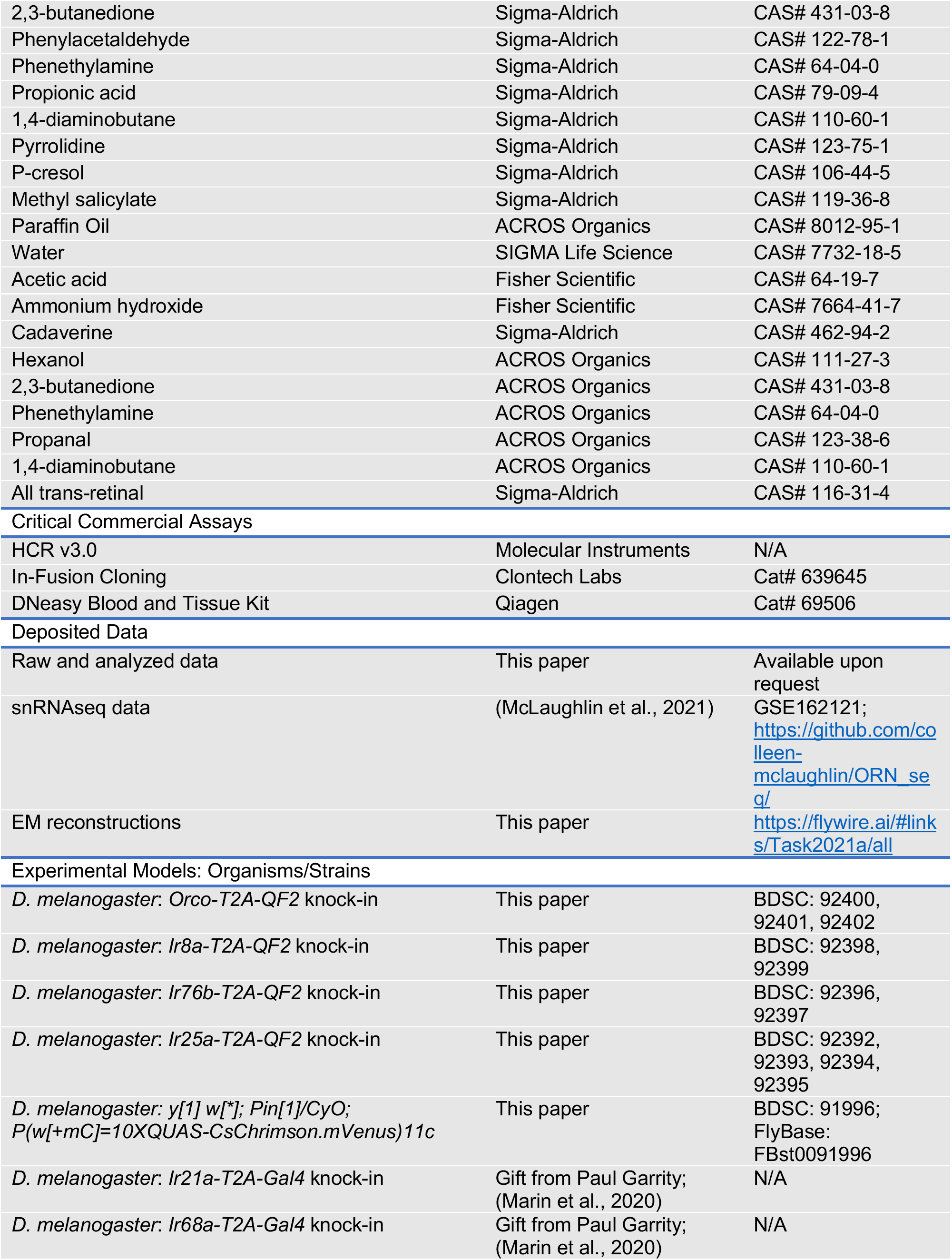
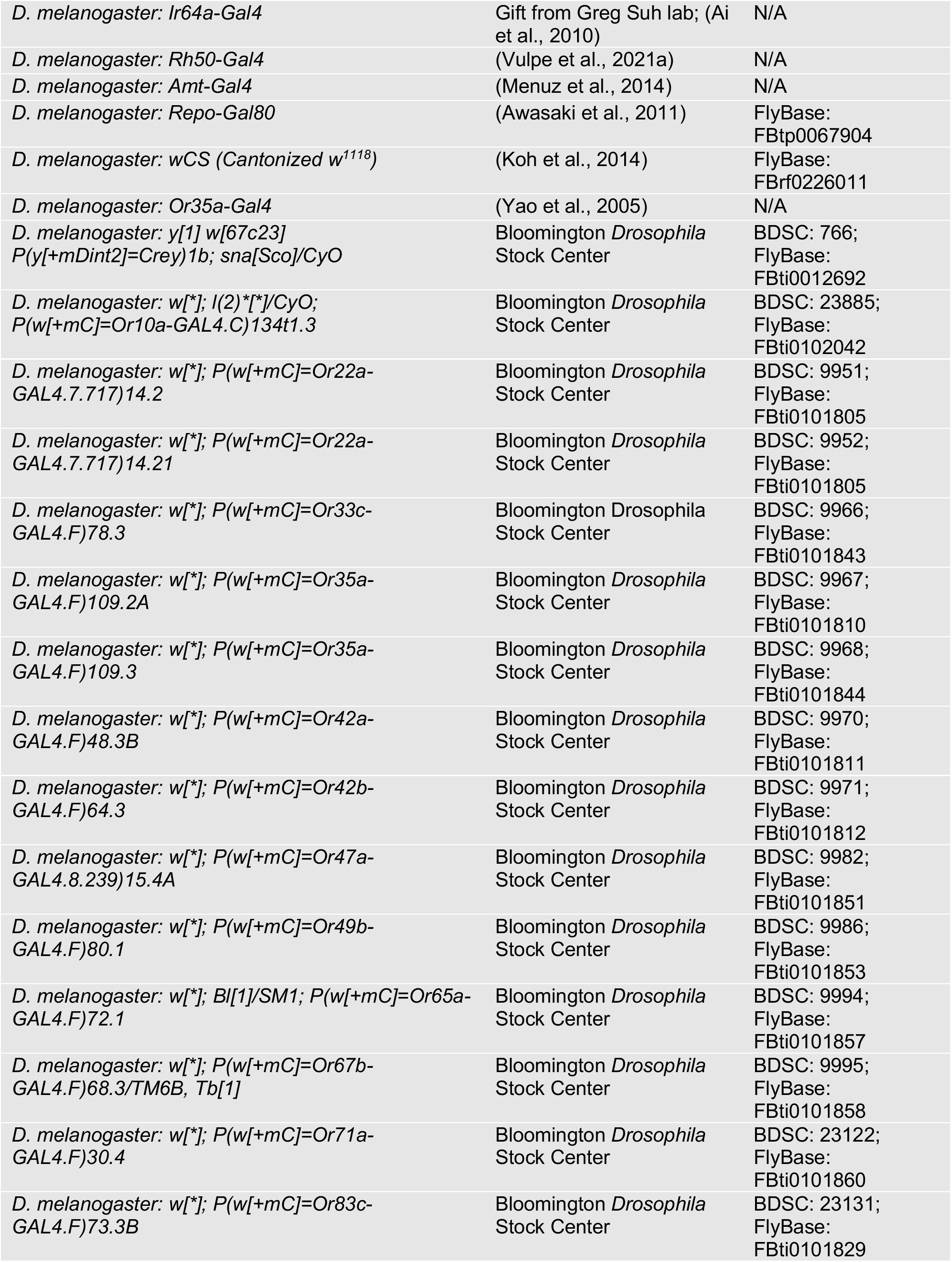
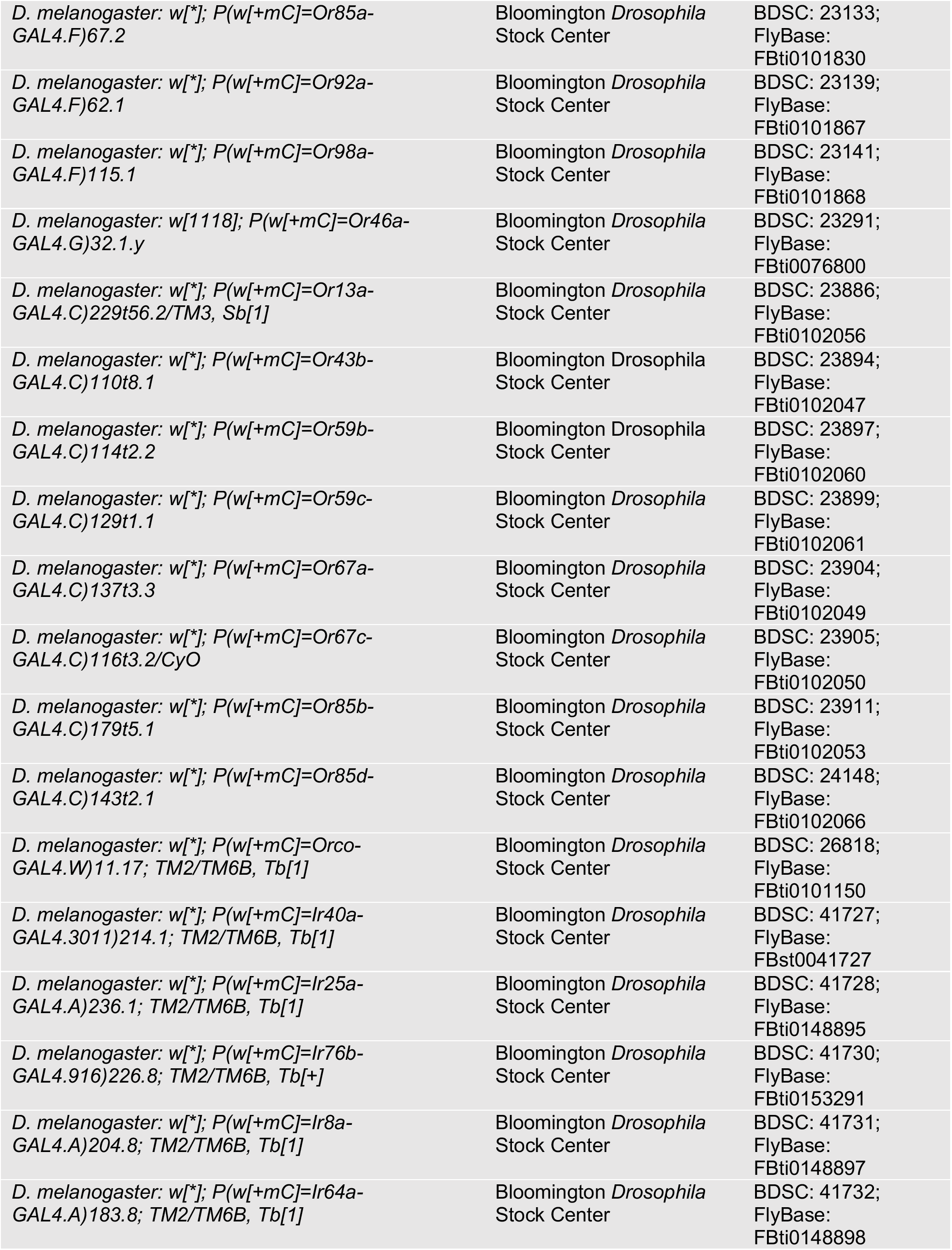
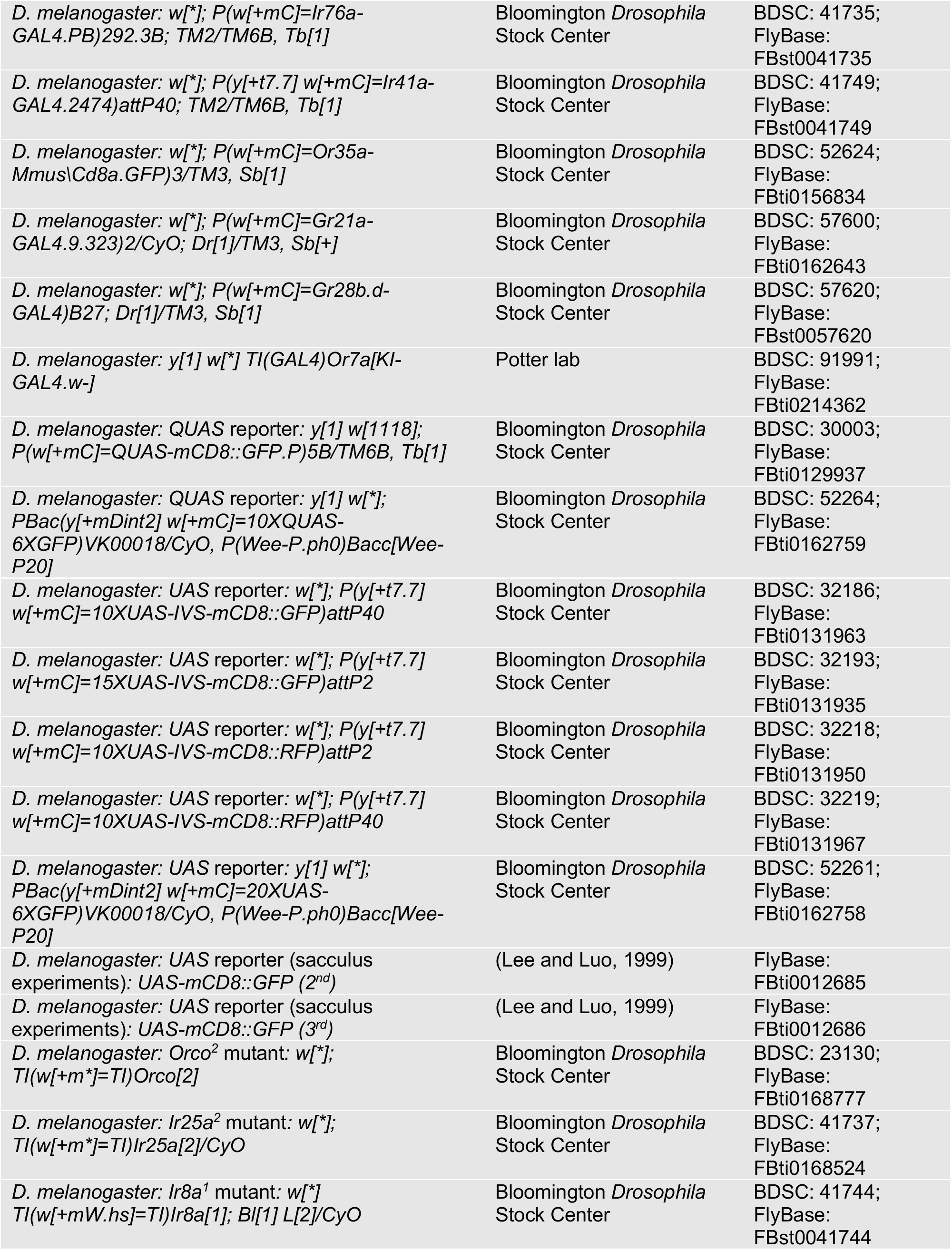
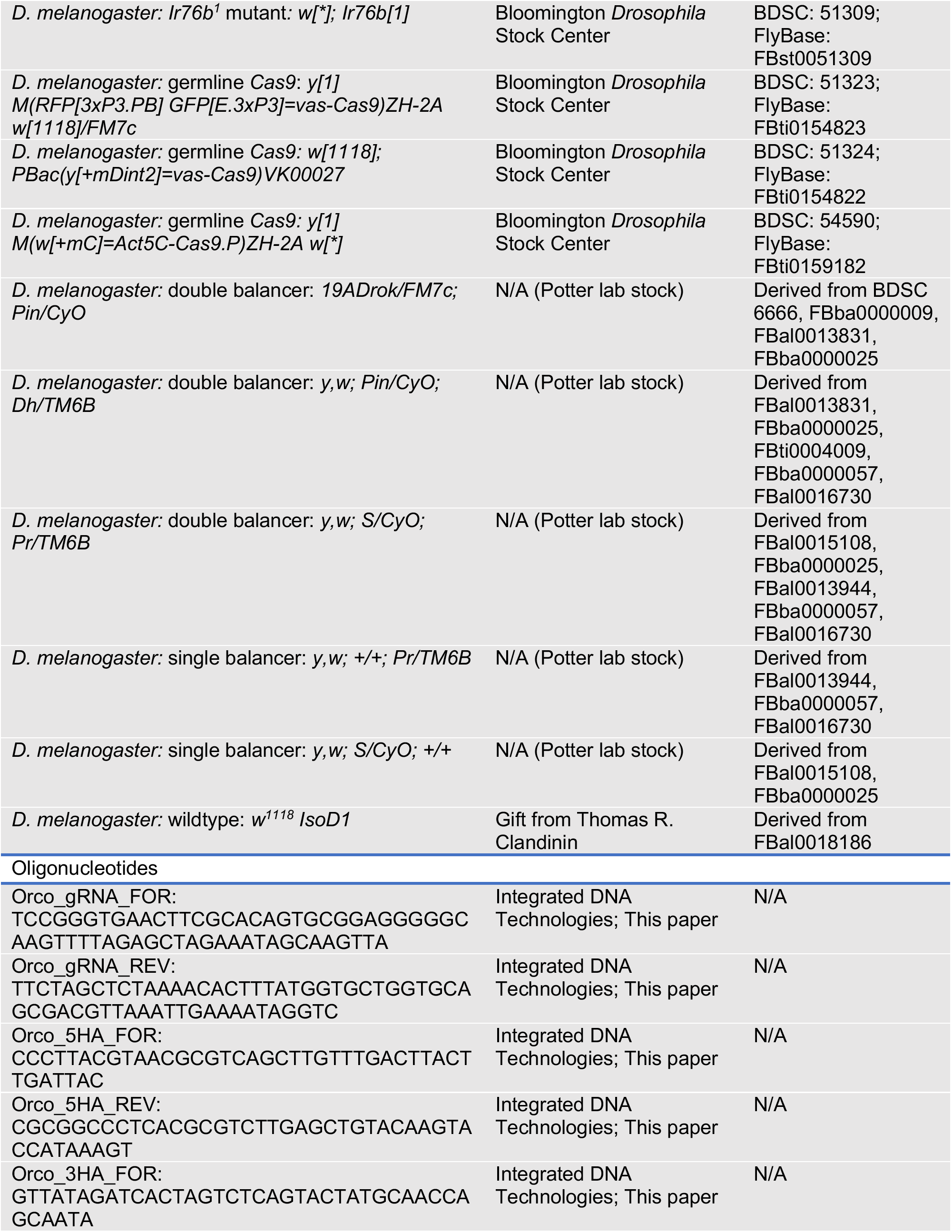
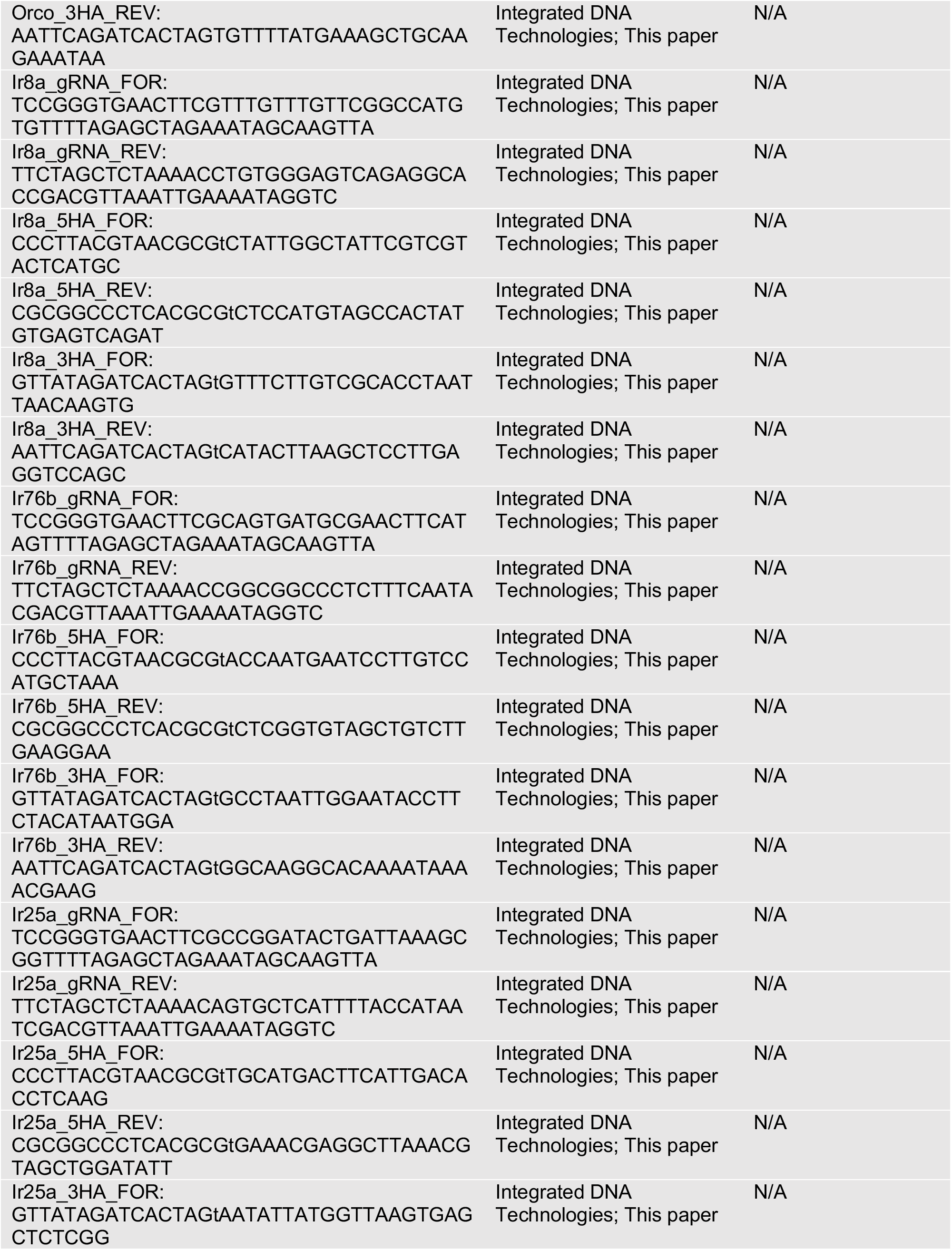
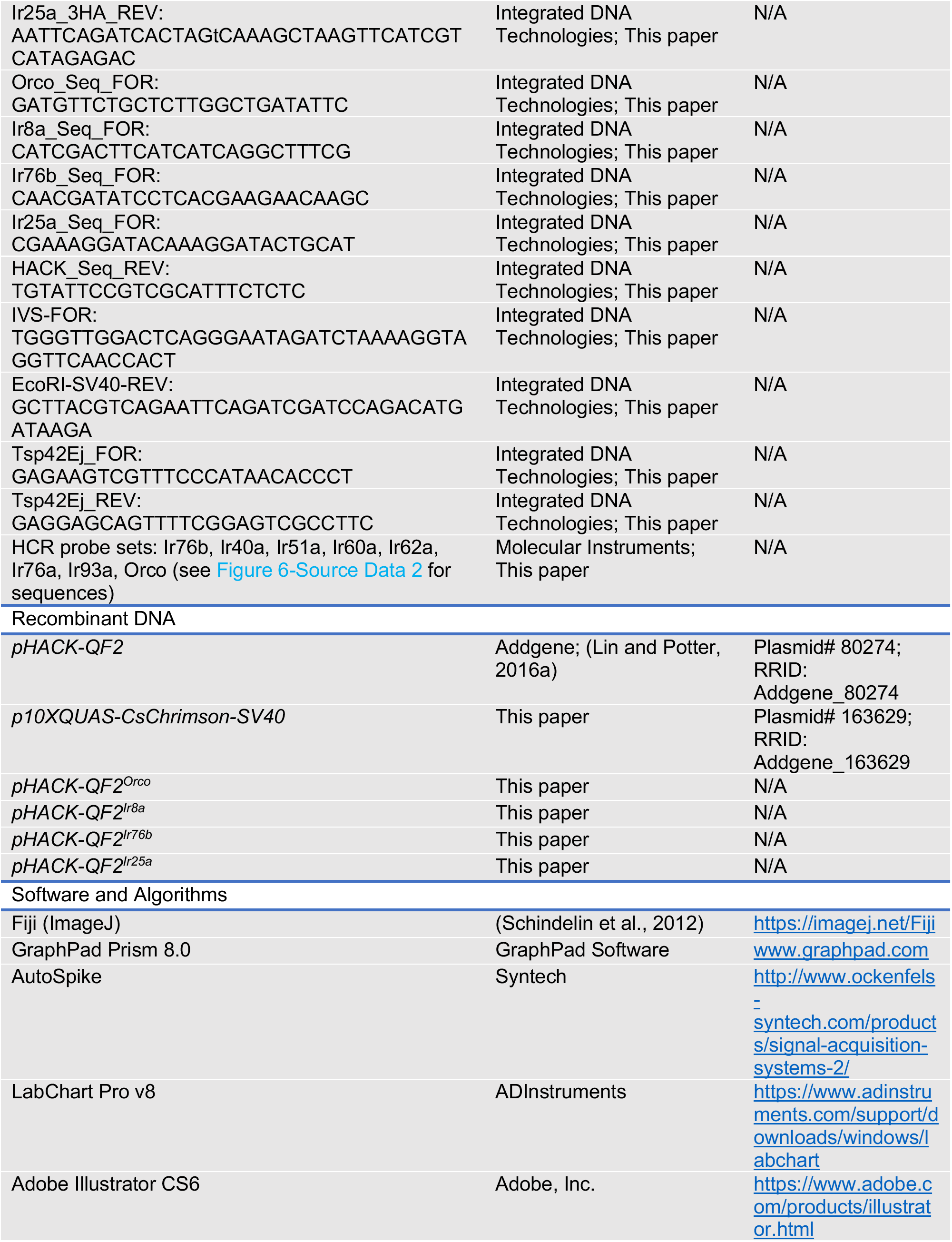
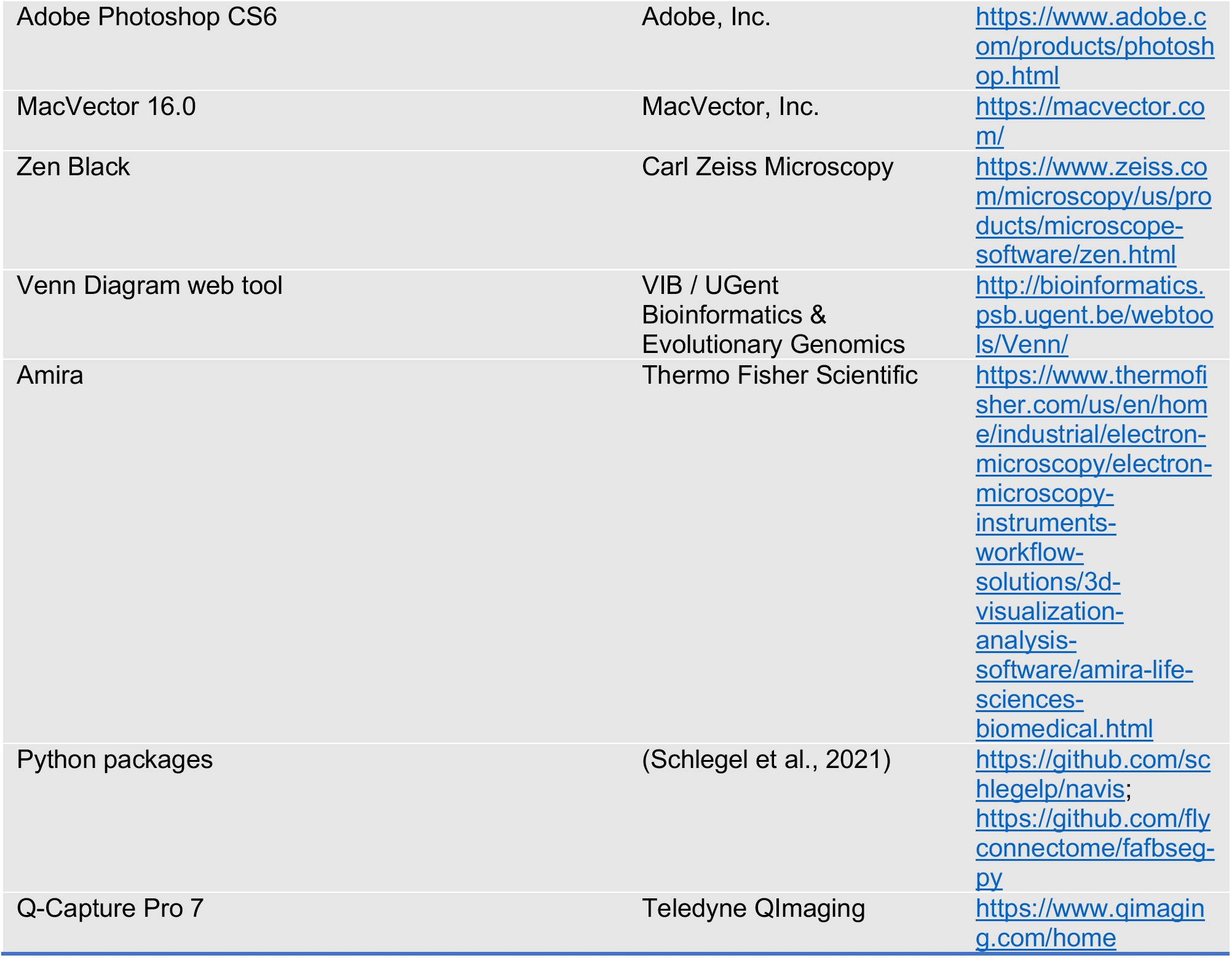

### Resource Availability

#### Lead Contact

Further information and requests for resources and reagents should be directed to and will be fulfilled by the Lead Contact, Christopher J. Potter.

#### Materials Availability

Fly lines generated in this study have been deposited to Bloomington *Drosophila* Stock Center.

#### Data and Code Availability

snRNAseq dataset analyzed in this paper is published in (McLaughlin et al., 2021). Sequencing reads and pre-processed sequencing data are available in the NCBI Gene Expression Omnibus (GSE162121). Python code for generating figures is publicly available on GitHub (https://github.com/colleen-mclaughlin/ORN_seq/). VM6 reconstructions using FlyWire can be viewed at https://flywire.ai/#links/Task2021a/all. All other data are available upon request.

### Experimental Model and Subject Details

#### Fly Husbandry

Fly stocks were maintained at 20 – 25°C on standard cornmeal-agar food. Male and female flies used for experiments were 3 – 11 days old, unless otherwise noted.

#### Fly Stocks

Fly lines used in this paper can be found in the Key Resources Table.

Generation of *QUAS-CsChrimson*: the sequence of CsChrimson. Venus was PCR amplified from the genomic DNA of *UAS-CsChrimson.mVenus* flies (Klapoetke et al., 2014) and cloned into the *10XQUAS* vector (Addgene #163629). Fly line was established through random *P*-element insertion. Cloning was confirmed with Sanger sequencing (Genewiz) before being sent for injection (Rainbow Transgenic Flies, Inc). Primers used for PCR amplification and In-Fusion cloning:

IVS-FOR:

TGGGTTGGACTCAGGGAATAGATCTAAAAGGTAGGTTCAACCACT

EcoRI-SV40-REV:

GCTTACGTCAGAATTCAGATCGATCCAGACATGATAAGA

While performing co-labeling experiments, we discovered that several *OrX-Gal4* lines label multiple glomeruli, and thus do not accurately represent single OSN classes. These lines were excluded from analyses and should be used with caution: *Or33c-Gal4* (BDSC# 9966), *Or42a-Gal4* (BDSC# 9970), *Or43b-Gal4* (BDSC# 23894), *Or59b-Gal4* (BDSC# 23897), *Or65a-Gal4* (BDSC# 9994), *Or85a-Gal4* (BDSC# 23133), *Or85b-Gal4* (BDSC# 23911). We also found that the following *Or35a* lines label the newly identified VM6l glomerulus in addition to VC3: *Or35a-Gal4* (BDSC# 9967), *Or35a-Gal4* (BDSC# 9968), *Or35a-mCD8.GFP* (BDSC# 52624), as well as an *Or35a-Gal4* line from the Carlson lab (Yao et al., 2005).

#### Generation of HACK Knock-in Lines

The HACK knock-in approach requires two components: a donor construct and Cas9 (Lin and Potter, 2016a). The donor includes *gRNAs* specific to the target gene, as well as the template for HDR-mediated insertion of *T2A-QF2* into the genome (Figure 2A, middle row). This template includes ∼1kb homology arms directly up and downstream of the gene’s stop codon flanking a cassette containing *T2A-QF2* and a *3XP3-mCherry* fluorescent eye marker (see Figure 2-Figure Supplement 1D-E and Figure 2-Figure Supplement 3A-B). Outside of these homology arms, the construct has two *RNA polymerase III U6* promoters driving independent expression of two *gRNAs* specific to the region around the target gene’s stop codon (Port et al., 2014). Two *gRNAs* were used to increase the probability of successfully inducing double stranded breaks in the target (Port et al., 2014). The knock-in construct replaces the target gene’s stop codon (Figure 2A, bottom row), and introduces a transcriptional stop at the end of *QF2*.

The donor construct can be supplied in one of two ways (Figure 2-Figure Supplement 1). The first is to inject the HACK construct directly into embryos expressing Cas9 in their germline (direct injection method) (Figure 2-Figure Supplement 1A-B). The second approach is to establish transgenic donor lines through random *P*-element insertion or *ΦC31* integration (Bischof et al., 2007; Gloor et al., 1991; Groth et al., 2004) of the construct into the genome, followed by genetic crosses with germline Cas9 flies for generation of the knock-in (cross method)(Figure 2-Figure Supplement 1C). Only one (direct injection method) or two (cross method) generations of crosses are required for the creation of a knock-in line (Figure 2-Figure Supplement 1B-C). The HACK 3XP3-mCherry selection marker is bright but shows positional effects (Figure 2-Figure Supplement 3A). Potential knock-in flies can be screened at the adult stage (Figure 2-Figure Supplement 3A), or at the larval or pupal stages (Figure 2-Figure Supplement 1D). We generated *T2A-QF2* knock-in lines for all four co-receptor genes using the direct injection method. Additionally, we tested the feasibility of the cross approach with two genes: *Orco* and *Ir25a*. Knock-ins were confirmed by PCR genotyping and sequencing (Figure 2-Figure Supplement 3B-D), and by crosses to a *QUAS-GFP* reporter to check for expression in the brain (*QUAS-mCD8::GFP* was used only to establish the *Orco-T2A-QF2* knock-in line, after which the reporter was removed via genetic crosses. For all antennal lobe analyses, we used the *10XQUAS-6XGFP* reporter line). We found no difference in expression pattern in the brain between these two approaches (Figure 2-Figure Supplement 1G). After establishing a knock-in line, the 3XP3-mCherry marker can be removed via *Cre* recombination (Siegal and Hartl, 1996). This can be useful as 3XP3-mCherry is expressed broadly throughout the fly nervous system and can interfere with red fluorescent reporters (Figure 2-Figure Supplement 1E). We produced two unmarked knock-in lines (for *Orco* and *Ir25a*) and confirmed no difference in brain GFP expression between marked and unmarked lines (Figure 2-Figure Supplement 1F).

Both approaches produced knock-ins at high rates (Table 1). Efficiency was calculated as the number of potentially HACKed knock-in flies (mCherry+), divided by the total number of flies from the given cross (G_1_ or F_2_ progeny; see Figure 2-Figure Supplement 1A-C). We further calculated the percentage of founders producing knock-in lines, as this gives an indication of effort (how many initial crosses need to be set up to produce a knock-in). The aggregate efficiency rates for a given target locus ranged from 8% for Ir8a to 33% for Orco (Table 1); however, for individual crosses, efficiency rates were as high as 100% (see Table 1-Source Data 1), meaning that all progeny were potential mCherry+ knock-ins. For the two genes for which we created knock-in lines via both direct injection and genetic cross (*Orco* and *Ir25a*), we found efficiency rates comparable between approaches (*Orco*: 33% for direct injection, 28% for cross; *Ir25a*: 23% for direct injection, 24% for cross). For the direct injection approach, we tested 51 independent knock-in lines across the four target genes and found 100% to be correctly targeted events (Table 1). However, for the genetic cross approach, of the 32 independent knock-in lines tested for the two target genes, 6 (∼19%) had the HACK mCherry eye marker but did not have *QF2*-driven GFP expression in the brain.

Information on plasmid construction can be found in Method Details below. All *Drosophila* embryo injections were performed by Rainbow Transgenic Flies, Inc. (Camarillo, CA). For HACKing via genetic cross, *Orco-T2A-QF2* and *Ir25a-T2A-QF2* constructs were injected into *w^1118^* flies for *P*-element insertion, and donor lines were established on the second or third chromosomes by crossing to double balancers (see Key Resources Table). Donor lines were then crossed to *Vas-Cas9* (BDSC# 51323). Knock-in lines were established from *cis*-chromosomal HACK (donor line on same chromosome as target gene) (Lin and Potter, 2016a). For HACKing via direct injection, knock-in constructs were injected into the following lines: *Vas-Cas9* (BDSC# 51324) for *Ir8a*; *Act5C-Cas9* (BDSC# 54590) for *Orco*, *Ir76b*, and *Ir25a*. The following lines were used to verify knock-in expression: *QUAS-mCD8::GFP* (BDSC# 30003), *10XQUAS-6XGFP* (BDSC# 52264). Knock-in lines were confirmed by PCR genotyping (Phusion, NEB) and Sanger sequencing (Genewiz). Unmarked *Orco-T2A-QF2* and *Ir25a-T2A-QF2* knock-in lines were generated by crossing mCherry+ knock-in flies to the Crey+1B fly line (see Key Resources Table) (Siegal and Hartl, 1996).

To investigate the effect of *T2A-QF2* knock-in on gene function, we performed single sensillum recordings (SSR) on homozygous flies for each co-receptor knock-in (Figure 2-Figure Supplement 2), comparing their responses to *wildtype* flies to panels of Orco- and Ir-dependent odorants (Abuin et al., 2011; de Bruyne et al., 2001; Lin and Potter, 2015). In ab2 basiconic sensilla, *Orco-T2A-QF2* knock-in flies had slightly lower baseline activity as compared to *wildtype* (Figure 2-Figure Supplement 2A); however, there were no significant differences in odor-evoked activity between these two genotypes across all stimuli tested (Figure 2-Figure Supplement 2B). In ac2 coeloconic sensilla, responses of *Ir8a-T2A-QF2* knock-in flies to hexanol and cadaverine were slightly lower than *wildtype* (Figure 2-Figure Supplement 2D); however, these are not typically considered Ir8a-dependent odorants. Responses of the *Ir8a-T2A-QF2* knock-in to Ir8a-dependent odorants (Abuin et al., 2011) were similar to *wildtype* controls (example trace in Figure 2-Figure Supplement 2C, quantification in Figure 2-Figure Supplement 2D). Responses of *Ir76b-T2A-QF2* knock-in ac2 neurons to phenethylamine and acetic acid differed slightly from *wildtype* controls (Figure 2-Figure Supplement 2E-F). The reasons for this are unclear. The largest difference in responses between a knock-in and *wildtype* were for *Ir25a-T2A-QF2* (Figure 2-Figure Supplement 2G-H); the knock-in has significantly reduced or abolished responses to Ir25a-dependent odorants, recapitulating an *Ir25a* mutant phenotype (Abuin et al., 2011; Silbering et al., 2011). See also Figure 2-Source Data 1.

### Method Details

#### Plasmid Construction

The construction of *QF2^X-HACK^* knock-in plasmids requires three steps of cloning, as previously described (Lin and Potter, 2016a). All knock-in constructs were created using the *pHACK-QF2* plasmid (Addgene #80274) as the backbone. The backbone was digested with the following enzymes for cloning: *MluI* for the 5’ homology arms; *SpeI* for the 3’ homology arms; and *BbsI* for the gRNAs. All cloning was performed using In-Fusion Cloning (Clontech #639645). The homology arms were PCR amplified from genomic DNA extracted from *wildtype* flies using the DNeasy Blood and Tissue Kit (Qiagen #69506), while the gRNAs were PCR amplified using the *pHACK-QF2* backbone as a template, with the primers themselves containing the gRNA target sequences. All homology arms were approximately 1 kb (Orco: 5HA = 1012 bp, 3HA = 1027 bp; Ir8a: 5HA = 1027 bp, 3HA = 1079 bp; Ir76b: 5HA = 997 bp, 3HA = 956 bp; Ir25a: 5HA = 1119 bp, 3HA = 990 bp). gRNAs were selected by analyzing the region around the stop codon of each gene using an online tool (https://flycrispr.org/) (Gratz et al., 2014). When possible, gRNAs were chosen to minimize potential off-target cleavage sites (zero predicted for *Orco*, *Ir8a*, and *Ir76b*; one predicted for *Ir25a*, discussed below). They were selected such that: one gRNA targeted upstream of the stop codon, within the last exon of the gene; the second gRNA targeted downstream of the stop codon, within the 3’UTR; and the two gRNAs were <100 bp apart. In order to prevent the gRNAs from targeting the homology arms, three synonymous nucleotide substitutions were made in each homology arm. The final knock-in lines did not always have all three substitutions (see Figure 2-Figure Supplement 3D), possibly due to PCR or HDR error. Note that due to the way the primers are designed, each targeted gene loses a small portion of its native 3’UTR (Orco = 72 bp, Ir8a = 31 bp, Ir76b = 27 bp, Ir25a = 24 bp). Cloning was confirmed with Sanger sequencing (Genewiz) before being sent for injection (Rainbow Transgenic Flies, Inc). Below are the gRNAs used for each gene, with the PAM sequence in parentheses.

Orco:

CTGCACCAGCACCATAAAGT (AGG)

GCACAGTGCGGAGGGGGCAA (GGG)

Ir8a:

GTTTGTTTGTTCGGCCATGT (TGG)

GGTGCCTCTGACTCCCACAG (TGG)

Ir76b:

GCAGTGATGCGAACTTCATA (TGG)

GTATTGAAAGAGGGCCGCCG (AGG)

Ir25a:

GCCGGATACTGATTAAAGCG (CGG)

ATTATGGTAAAATGAGCACT (CGG)

PRIMERS

*Italics* = In-Fusion cloning 15 b.p. overhang; **bold** = gRNA; lowercase = adding back restriction site; <u>underline</u> = synonymous substitution to prevent Cas9 targeting of donor construct.

PCR Primers for Cloning:

Orco_gRNA_FOR:

*TCCGGGTGAACTTC***GCACAGTGCGGAGGGGGCAA**GTTTTAGAGCTAGAAATAGCAAGTTA

Orco_gRNA_REV:

*TTCTAGCTCTAAAAC***ACTTTATGGTGCTGGTGCAG**CGACGTTAAATTGAAAATAGGTC

Orco_5HA_FOR:

*CCCTTACGTAACGCG*tCAGCTTGTTTGACTTACTTGATTAC

Orco_5HA_REV:

*CGCGGCCCTCACGCG*tCTTGAGCTG<u>T</u>AC<u>A</u>AG<u>T</u>ACCATAAAGT

Orco_3HA_FOR:

*GTTATAGATCACTAG*tCTC<u>A</u>G<u>T</u>ACT<u>A</u>TGCAACCAGCAATA

Orco_3HA_REV:

*AATTCAGATCACTAG*tGTTTTATGAAAGCTGCAAGAAATAA

Ir8a_gRNA_FOR:

*TCCGGGTGAACTTCG***TTTGTTTGTTCGGCCATGT**GTTTTAGAGCTAGAAATAGCAAGTTA

Ir8a_gRNA_REV:

*TTCTAGCTCTAAAAC***CTGTGGGAGTCAGAGGCAC**CGACGTTAAATTGAAAATAGGTC

Ir8a_5HA_FOR:

*CCCTTACGTAACGCG*tCTATTGGCTATTCGTCGTACTCATGC

Ir8a_5HA_REV:

*CGCGGCCCTCACGCG*tCTCCATGTAGCCACT<u>A</u>TG<u>T</u>GAGTCAGA<u>T</u>

Ir8a_3HA_FOR:

*GTTATAGATCACTAG*tGTTTCTTGTCGCACCTAATTAACAAGTG

Ir8a_3HA_REV:

*AATTCAGATCACTAG*tCATACTTAAGCTCCTTGAGGTCCAGC

Ir76b_gRNA_FOR:

*TCCGGGTGAACTTCG***CAGTGATGCGAACTTCATA**GTTTTAGAGCTAGAAATAGCAAGTTA

Ir76b_gRNA_REV:

*TTCTAGCTCTAAAAC***CGGCGGCCCTCTTTCAATAC**GACGTTAAATTGAAAATAGGTC

Ir76b_5HA_FOR:

*CCCTTACGTAACGCG*tACCAATGAATCCTTGTCCATGCTAAA

Ir76b_5HA_REV:

*CGCGGCCCTCACGCG*tCTCGGTGTAGCTGT<u>C</u>TTGAA<u>G</u>GA<u>A</u>

Ir76b_3HA_FOR:

*GTTATAGATCACTAG*tGCCTAATTGGAATACCTTCTACATAATGGA

Ir76b_3HA_REV:

*AATTCAGATCACTAG*tGGCAAGGCACAAAATAAAACGAAG

Ir25a_gRNA_FOR:

*TCCGGGTGAACTTC***GCCGGATACTGATTAAAGCG**GTTTTAGAGCTAGAAATAGCAAGTTA

Ir25a_gRNA_REV:

*TTCTAGCTCTAAAAC***AGTGCTCATTTTACCATAAT**CGACGTTAAATTGAAAATAGGTC

Ir25a_5HA_FOR:

*CCCTTACGTAACGCG*tTGCATGACTTCATTGACACCTCAAG

Ir25a_5HA_REV:

*CGCGGCCCTCACGCG*tGAAACGAGGCTTAAACGT<u>A</u>GC<u>T</u>GGATA<u>T</u>T

Ir25a_3HA_FOR:

*GTTATAGATCACTAG*tAATATTATGGT<u>T</u>AA<u>G</u>TGAGC<u>T</u>CTCGG

Ir25a_3HA_REV:

*AATTCAGATCACTAG*tCAAAGCTAAGTTCATCGTCATAGAGAC

Genotyping and Sequencing Primers (PCR fragment size):

Orco_Seq_FOR (∼2kb):

GATGTTCTGCTCTTGGCTGATATTC

Ir8a_Seq_FOR (∼1.9kb):

CATCGACTTCATCATCAGGCTTTCG

Ir76b_Seq_FOR (∼1.9kb):

CAACGATATCCTCACGAAGAACAAGC

Ir25a_Seq_FOR (∼1.9kb):

CGAAAGGATACAAAGGATACTGCAT

HACK_Seq_REV (same for all):

TGTATTCCGTCGCATTTCTCTC

#### Checking for Off-target Effects

One of the gRNAs for *QF2^Ir25a-HACK^* had one predicted potential off-target cut site in the genome, in the *tetraspanin 42ej* (*Tsp42Ej*) gene. We sequenced this locus in the *Ir25a-T2A-QF2* knock-in line and compared the sequence to our *wildtype* lab stock. We found no evidence of indels in the knock-in line. Primers used:

Tsp42Ej_FOR:

GAGAAGTCGTTTCCCATAACACCCT

Tsp42Ej_REV:

GAGGAGCAGTTTTCGGAGTCGCCTTC

#### HACK Marker Screening

Adult flies were anaesthetized on a CO_2_ pad and screened in one of two ways: either with a Nightsea Stereo Microscope Fluorescence Adapter with the green SFA-GR LED light source (Nightsea LLC, Lexington, MA) and viewed with a Zeiss Stemi SV6 stereo microscope; or illuminated with an X-Cite 120Q excitation light source and viewed with a Zeiss SteREO Discovery V8 microscope equipped with a ds-Red filter.

#### Whole Animal Imaging

Whole adults were anaesthetized on ice before imaging. Whole larvae, pupae, or freshly dissected adult heads were affixed to slides with clear nail polish before imaging. All animals were imaged on an Olympus SZX7 microscope equipped with GFP and RFP filters. Animals were illuminated with an X-Cite Series 120Q light source. Images were acquired using a QImaging QIClick Cooled digital CCD camera and Q-Capture Pro 7 software. Multiple images were taken at different focal planes and then merged in Photoshop (CS6). Gain was adjusted in Fiji. Images appear in the following figures/panels: Figure 2B, D, F, H; Figure 2-Figure Supplement 1D; Figure 2-Figure Supplement 3A; Figure 2-Figure Supplement 4.

#### Immunohistochemistry

All flies were used at 3- to 11-days old. Apart from the cryosection protocols and portions of the antennal whole-mount protocol, all immunostaining steps were done on a nutator. All steps involving or following the addition of fluorescently conjugated secondary antibodies were done in the dark.

Brain and ventral nerve cord (VNC) staining was performed as in (Xie et al., 2018). Tissue was dissected in PBS and then fixed for 20 minutes at room temperature (RT) in 4% paraformaldehyde in PBT (1XPBS with 0.3% Triton X-100). After fixation, tissue was quickly rinsed three times with PBT, then put through three longer washes in PBT at RT (at least 15 minutes each). Tissue was blocked in PBT + 5% normal goat serum (NGS) at RT for at least 30 minutes, then transferred to block + primary antibody solution, and incubated at 4°C in primary antibodies for 1-2 days. Tissue was then washed three times with PBT at RT (at least 15 minutes per wash) and incubated in secondary antibody solution in block at 4°C for 1 day. Tissue was washed three final times with PBT at RT for 15 minutes each, and then mounted in SlowFade gold (Thermo Fisher Scientific #S36936). For experiments in which the 10XQUAS-6XGFP or 20XUAS-6XGFP reporters were used, the endogenous, unstained GFP signal was visualized, and no secondary green antibodies were used. Primary antibodies used: mouse anti-nc82 (DSHB, 1:25), rat anti-mCD8 (Thermo Fisher #14-0081-82, 1:100 or 1:200). Secondary antibodies used: Cy3 goat anti-rat (Jackson ImmunoResearch #112-165-167, 1:200), Cy3 goat anti-mouse (Jackson ImmunoResearch #115-165-166, 1:200), Alexa 647 goat anti-rat (Jackson ImmunoResearch #112-605-167, 1:200), Alexa 647 goat anti-mouse (Jackson ImmunoResearch #115-605-166, 1:200).

Whole-mount staining of maxillary palps was performed according to the brain staining protocol above, with the exception of a shorter fixation step (15 minutes). Primary antibodies used: rabbit anti-Ir25a (gift from Richard Benton, University of Lausanne, 1:100), rabbit anti-Orco (gift from Leslie Vosshall, Rockefeller University, 1:100), rat anti-elav (DSHB, 1:100). Secondary antibodies used: Cy3 goat anti-rabbit (Jackson ImmunoResearch #111-165-144, 1:200), Alexa 647 goat anti-rabbit (Jackson ImmunoResearch #111-605-144, 1:200), Alexa 647 goat anti-rat (Jackson ImmunoResearch #112-605-167, 1:200). For whole-mount staining of *Orco^2^* mutant palps, 3-day-old flies were used to check for Orco expression before neurons degenerate (Task and Potter, 2021).

The protocol for whole-mount staining of antennae was adapted from (Karim et al., 2014; Saina and Benton, 2013; Younger et al., 2020). Fly heads were dissected into CCD buffer (50 units chitinase, 1000 units chymotrypsin (25 mg of 40 units/mg), 10 mL HEPES larval buffer (119 mM NaCl, 48 mM KCl, 2 mM CaCl_2_, 2 mM MgCl_2_, 25 mM HEPES), 100 µl DMSO) on ice, then warmed on a 37°C heat block for 10 min. Heads were incubated in CCD buffer at 37°C while rotating for 1 hour 20 minutes. Antennae were subsequently dissected off heads into fixative solution (4% PFA in PBT). All subsequent steps were done without rotation to prevent antennae from sticking to the walls or lids of the tubes. Antennae were fixed at room temperature for 40 min, then washed with PBT three times at room temperature, at least 15 minutes each time, and blocked in PBT plus 5% NGS for at least 1 hour at room temperature. Antennae were incubated in primary antibodies in blocking solution at 4°C for four days, washed three times for 15 minutes each at room temperature, and incubated in secondary antibody solution at 4°C for three days. Antennae were then washed three times for 15 minutes each time at room temperature and mounted in SlowFade Gold. Primary antibody: rabbit anti-Orco (gift from Leslie Vosshall, 1:100); secondary antibody: Cy3 goat anti-rabbit (Jackson ImmunoResearch #111-165-144, 1:200). The endogenous GFP signal was visualized.

The cryosection protocol was adapted from (Spletter et al., 2007). Fly heads were dissected and lined up in cryomolds (Tissue-Tek #4565), covered with OCT compound (Tissue-Tek #4583), and frozen at −80°C. Samples were sectioned at ∼12 µm on a Microm HM 500 cryostat (Microm International GmbH, Walldorf, Germany) and collected on SuperFrost Plus slides (Fisher #12-550-15). Slides were stored at −80°C until further processing. Slides were fixed at RT for 15 minutes in 4% paraformaldehyde in PBT (1XPBS with 0.3% Triton X-100), washed three times in PBT at RT (15 minutes each), blocked at RT for at least 30 minutes in PBT + 2.5% NGS + 2.5% normal donkey serum (NDS), then incubated overnight at 4°C in primary antibodies in fresh block solution in a special humidified chamber. On the next day, slides were washed three times (15 minutes each) with PBT at RT, and then incubated in secondary antibodies in block at 4°C overnight in the same humidified chamber covered in foil. Finally, slides were washed three times (15 minutes each) with PBT at RT. DAPI (1:10,000) was included in the first wash as a nuclear counterstain. After washes, slides were mounted in SlowFade gold (Thermo Fisher Scientific #S36936). Primary antibody used: guinea pig anti-Ir8a (gift from Richard Benton, University of Lausanne, 1:1,000). Secondary antibody used: Cy3 donkey anti-guinea pig (Jackson ImmunoResearch #706-165-148, 1:200).

For sacculus staining, 7-10 day old flies were placed in an alignment collar. Their heads were encased in OCT (Tissue-Plus Fisher) in a silicone mold, frozen on dry ice, and snapped off. The head blocks were stored in centrifuge tubes at −80°C. A Leica cryostat was used to collect 20 µm sections of antennae. Immunohistochemical staining was carried out by fixing tissue in 4% paraformaldehyde for 10 minutes, followed by three 5 minute washes in 1XPBS. Tissue was washed in 1XPBS containing 0.2% Triton-X (PBST) for 30 minutes to permeabilize the cuticle. Lastly, tissue was washed in PBST containing 1% Bovine Serum Albumin (BSA) to block non-specific antibody binding. Primary antibody solution was made in PBST+1%BSA, 200 µl was pipetted onto each slide under bridged coverslips, and slides were placed at 4°C overnight to incubate. The following day, primary antibody was removed, and slides were washed three times for 10 minutes each in PBST. Secondary antibody solution was made in PBST+1%BSA, 200uL was pipetted onto each slide under bridged coverslips, and left at room temperature in a dark box to incubate for 2 hours. After the 2 hour incubation, slides were washed three times for 5 minutes each in PBST. After the last wash, the slides were allowed to dry in the dark staining box for ∼30 minutes before being mounted in Vectashield, coverslipped, and stored at 4°C. Primary antibodies: rabbit anti-Ir25a (gift from Richard Benton, University of Lausanne, 1:100), guinea pig anti-Ir8a (gift from Richard Benton, University of Lausanne, 1:100), rabbit anti-Ir64a (gift from Greg Suh, NYU/KAIST, 1:100). Secondary antibodies: Jackson Immuno Cy3 conjugated AffiniPure 568 goat anti-rabbit (111-165-144), 1:500; AlexaFluor 568 goat anti-guinea pig (A11075), 1:500.

#### In situ HCR

Cryosectioning for antennal *in situs* was performed as described above. The HCR protocol was adapted from Molecular Instruments HCR v3.0 protocol for fresh frozen or fixed frozen tissue sections (Choi et al., 2018). Slides were fixed in ice-cold 4% PFA in PBT for 15 minutes at 4°C, dehydrated in an ethanol series (50% EtOH, 70% EtOH, 100% EtOH, 100% EtOH, 5 minutes each step), and air dried for 5 minutes at RT. Slides were then incubated in proteinase K solution in a humidified chamber for 10 minutes at 37°C, rinsed twice with PBS and dried, then pre-hybridized for 10 minutes at 37°C in a humidified chamber. Slides were incubated in probe solution (0.4 pmol Ir76b probe) overnight in the 37°C humidified chamber. On day two, slides were washed with a probe wash buffer/SSCT series (75% buffer/25% SSCT, 50% buffer/50% SSCT, 25% buffer/75% SSCT, 100% SSCT, 15 minutes each) at 37°C, then washed for 5 min at RT with SSCT and dried. Slides were pre-amplified for 30 min at RT in the humidified chamber while hairpins were snap cooled (6 pmol concentration). Slides were incubated in fresh amplification buffer with hairpins overnight in a dark humidified chamber at RT. On day three, slides were rinsed in SSCT at RT (2 x 30 minutes, 1 x 10 minutes with 1:10,000 DAPI, 1 x 5 minutes) and mounted in SlowFade Diamond (Thermo Fisher S36972). For the overnight steps, slides were covered with HybriSlips (Electron Microscopy Sciences 70329-62) to prevent solution evaporation.

The whole-mount palp *in situ* protocol was adapted from a combination of (Prieto-Godino et al., 2020; Saina and Benton, 2013; Younger et al., 2020) and the *Drosophila* whole-mount embryo protocol from Molecular Instruments (Choi et al., 2016). All steps after dissection were performed while rotating unless otherwise noted. Fly mouthparts (palps and proboscises) were dissected into CCD buffer (same as for whole-mount IHC on antennae above), incubated for 20 minutes in CCD at 37°C (5 minutes on heat block, 15 minutes rotating), then pre-fixed in 4% PFA in PBT for 20 minutes at RT. Tissue was washed with 0.1% PBS-Tween on ice (4 x 5 minutes), incubated for 1 hour at RT in 80% methanol/20% DMSO, and washed for 10 minutes in PBS-Tween at RT. Tissue was then incubated in Proteinase K solution (1:1,000) in PBS-Tween at RT for 30 minutes, then washed in PBS-Tween at RT (2 x 10 minutes) and post-fixed in 4% PFA in PBS-Tween at RT for 20 minutes. After post-fixation, tissue was washed in PBS-Tween at RT (3 x 15 minutes), then pre-hybridized in pre-heated probe hybridization buffer at 37°C for 30 minutes. Tissue was incubated in probe solution (2 – 4 pmol in hybridization buffer) at 37°C for two nights. On day three, tissue was washed in pre-heated probe wash buffer at 37°C (5 x 10 minutes), then washed in SSCT (1XSSC plus 1% Tween) at RT (2 x 5 minutes). Tissue was pre-amplified with RT-equilibrated amplification buffer at RT for 10 minutes, then incubated in hairpin mixture (6 pmol snap-cooled hairpins in amplification buffer) in the dark at RT overnight. On day four, tissue was washed at RT with SSCT (2 x 5 minutes, 2 x 30 minutes, 1 x 5 minutes), then mounted in SlowFade Diamond (Thermo Fisher S36972). Sequences for all *in situ* probes can be found in Figure 6-Source Data 2. Example images for palp *in situs* can be found in Figure 6-Figure Supplement 2.

#### Confocal Imaging and Analysis

Brains, ventral nerve cords, maxillary palps, and antennal cryosections were imaged on a Zeiss LSM 700 confocal microscope equipped with Fluar 10x/0.50 air M27, LCI Plan-Neofluar 25x/0.8 water Korr DIC M27, and C-Apochromat 63x/1.2 water Korr M27 objectives. Images were acquired at 512 x 512-pixel resolution with 0.58, 2.37 or 6.54 µm z-steps for each objective, respectively. For illustration purposes, confocal images were processed in Fiji/ImageJ to collapse Z-stacks into a single image using maximum intensity projection. Where noted, single slices or partial z projections were used as opposed to full stacks. For co-labeling experiments, Fiji was used to convert red LUTs to orange for a colorblind-friendly palette. Similarly, in Figure 2-Figure Supplement 1E, Fiji was used to convert magenta LUT to blue for clarity. For Figure 4B, Fiji was used to convert the two-channel maximum intensity projection to a grey LUT, and the cell-counting plug-in was used in separate channels to identify single- and double-labeled cells. Fiji was also used to adjust the gain in separate channels in all figures/images; no other image processing was performed on the confocal data. For Figure 7A, glomeruli were assigned to the categories strong, intermediate, and weak by visual inspection of the strength of their innervation compared to the previously reported glomeruli for each respective knock-in line. Strong glomeruli generally have similar brightness/intensity of GFP signal as most of the originally reported glomeruli for the given knock-in line.

For sacculus staining (Figure 5C-E), slides were imaged on a Nikon A1R confocal microscope in the UConn Advanced Light Microscopy Facility with a 40X oil immersion objective at 1024 x 1024-pixel resolution. Stacks of images (0.5 µm z-step size) were gathered and analyzed with ImageJ/Fiji software. Image processing was performed as described above.

Magnification used:

10X: Figure 1D (Brain), Figure 2-Figure Supplement 1E

25X: Figure 2-Figure Supplement 1F-G, Figure 3-Figure Supplement 1A-B

40X: Figure 5C-E

63X: Figure 2C, E, G, I; Figure 3A-D, Figure 4B-C, Figure 5A-B, Figure 2-Figure Supplement 3E-H, Figure 3-Figure Supplement 2A-E, Figure 5-Figure Supplement 1, Figure 6-Figure Supplement 2

#### Basiconic Single Sensillum Recordings

Flies were immobilized and visualized as previously described (Lin and Potter, 2015). Basiconic sensilla were identified either using fluorescent-guided SSR (for ab3 sensilla) or using reference odorants (for ab2 and pb1-3 sensilla) (de Bruyne et al., 1999; Lin and Potter, 2015). The glass recording electrode was filled with Beadle-Ephrussi Ringer’s solution (7.5g of NaCl + 0.35g of KCl + 0.279g of CaCl_2_-2H_2_O in 1L H_2_O). Extracellular activity was recorded by inserting the glass electrode into the shaft or base of the sensillum of 3- to 10-day-old flies (unless otherwise specified in the young *Orco^2^* mutant experiments). A tungsten reference electrode was inserted into the fly eye. Signals were amplified 100X (USB-IDAC System; Syntech, Hilversum, the Netherlands), input into a computer via a 16-bit analog-digital converter and analyzed off-line with AUTOSPIKE software (USB-IDAC System; Syntech). The low cutoff filter setting was 50 Hz, and the high cutoff was 5 kHz. Stimuli consisted of 1000 ms air pulses passed over odorant sources. The Δ spikes/second was calculated by counting the spikes in a 1000 ms window from ∼500 ms after odorant stimuli were triggered, subtracting the spikes in a 1000 ms window prior to each stimulation. For ab3 recordings from *wildtype*, *Orco^2^* mutant, and *Ir25a^2^* mutant flies, spikes were counted in a 500 ms window from the start of the response and multiplied by 2. Then, the spikes in the 1000 ms window prior to stimulation was subtracted from this to calculate the Δ spikes/second. Stimuli used: mineral oil (Sigma CAS #8042-47-5), ethyl acetate (Sigma CAS #141-78-6), pentyl acetate (Sigma CAS #628-63-7), benzaldehyde (Sigma CAS #100-52-7), ethyl butyrate (Sigma CAS #105-54-4), hexanol (Sigma CAS #111-27-3), e2-hexenal (Sigma CAS #6728-26-3), geranyl acetate (Sigma CAS # 105-87-3), 2-heptanone (Sigma CAS # 110-43-0), 1-octen-3-ol (Sigma CAS #3391-86-4), 2,3-butanedione (Sigma CAS #431-03-8), phenylacetaldehyde (Sigma CAS # 122-78-1), phenethylamine (Sigma CAS # 64-04-0), propionic acid (Sigma CAS #79-09-4), 1,4-diaminobutane (Sigma CAS #110-60-1), pyrrolidine (Sigma CAS #123-75-1), p-cresol (Sigma CAS #106-44-5), and methyl salicylate (Sigma CAS #119-36-8). Odorants were dissolved in mineral oil at a concentration of 1%, and 20 µl of solution was pipetted onto filter paper in a glass Pasteur pipette. Stimuli were delivered by placing the tip of the Pasteur pipette through a hole in a plastic pipette (Denville Scientific Inc, 10ml pipette) that carried a purified continuous air stream (8.3 ml/s) directed at the antenna or maxillary palp. A solenoid valve (Syntech) diverted delivery of a 1000 ms pulse of charcoal-filtered air (5 ml/s) to the Pasteur pipette containing the odorant dissolved on filter paper. Fresh odorant pipettes were used for no more than 5 odorant presentations. *Ir25a^2^* and *Orco^2^* mutant fly lines were outcrossed into the *w^1118^ wildtype* genetic background. Full genotypes for ab3 fgSSR were: *Pin/CyO;Or22a-Gal4,15XUAS-IVS-mcd8GFP/TM6B* (*wildtype*), *Ir25a^2^*;*Or22a-Gal4,15XUAS-IVS-mcd8GFP/TM6B* (*Ir25a^2^ mutant*), *Or22a-Gal4/10XUAS-IVS-mcd8GFP (attp40);Orco^2^* (*Orco^2^ mutant*). These stocks were made from the following Bloomington Stocks (outcrossed to the Potter lab *w^1118^* genetic background): BDSC# 9951, 9952, 23130, 32186, 32193, 41737.

#### Coeloconic Single Sensillum Recordings

Coeloconic SSR was performed similarly as for basiconic sensilla. 3-5-day old female flies were wedged in the tip of a 200 µl pipette, with the antennae and half the head exposed. A tapered glass electrode was used to stabilize the antenna against a coverslip. A BX51WI microscope (Olympus) was used to visualize the prep, which was kept under a 2,000mL/min humidified and purified air stream. A borosilicate glass electrode was filled with sensillum recording solution (Kaissling and Thorson, 1980) and inserted into the eye as a reference electrode. An aluminosilicate glass electrode was filled with the same recording solution and inserted into individual sensilla. Different classes of coeloconic sensilla were identified by their known location on the antenna and confirmed with their responses to a small panel of diagnostic odors. No more than 4 sensilla per fly were recorded. Each sensillum was tested with multiple odorants, with a rest time of at least 10s between applications. The odorants used were acetic acid (Fisher, 1%, CAS# 64-19-7), ammonium hydroxide (Fisher, 0.1%, CAS# 7664-41-7), cadaverine (Sigma-Aldrich, 1%, CAS# 462-94-2), hexanol (ACROS Organics, 0.001%, CAS# 111-27-3), 2,3-butanedione (ACROS Organics, 1%, CAS# 431-03-8), phenethylamine (ACROS Organics, 1%, CAS# 64-04-0), propanal (ACROS Organics, 1%, CAS# 123-38-6), and 1,4-diaminobutane (ACROS Organics, 1%, CAS# 110-60-1). Odorants were diluted in water or paraffin oil. Odorant cartridges were made by placing a 13 mm antibiotic assay disc (Whatman) into a Pasteur pipette, pipetting 50 µl odorant onto the disc, and closing the end with a 1 mL plastic pipette tip. Each odorant cartridge was used a maximum of four times. The tip of the cartridge was inserted into a hole in the main airflow tube, and odorants were applied at 500mL/min for 500ms. Delivery was controlled via LabChart Pro v8 software (ADInstruments), which directed the opening and closing of a Lee valve (02-21-08i) linked to a ValveBank 4 controller (AutoMate Scientific). Extracellular action potentials were collected with an EXT-02F amplifier (NPI) with a custom 10X head stage. Data were acquired and AC filtered (300-1,700 Hz) at 10kHz with a PowerLab 4/35 digitizer and LabChart Pro v8 software. Spikes were summed in coeloconic recordings due to their similar sizes, and they were counted over a 500 ms window, starting at 100 ms after stimulus onset.

#### Optogenetics

*Ir25a-T2A-QF2* was crossed to *QUAS-CsChrimson* #11C and double balanced to establish a stable stock (*Ir25a-T2A-QF2*/*CyO; QUAS-CsChrimson* #11C/*TM6B)*. Newly eclosed flies (age < 1 day old) were transferred to fly vials containing 0.4 mM all trans-retinal in fly food (Sigma-Aldrich #R2500, dissolved in pure DMSO with stock concentration of 0.4 M). Vials with flies were kept in the dark for at least 4 days before experiments. 627nm LED light source (1-up LED Lighting Kit, PART #: ALK-1UP-EH-KIT) powered by an Arduino Uno (www.arduino.cc/en/Main/ArduinoBoardUno) was used to activate CsChrimson. By setting the voltage to 2V and the distance of the light source to 20 cm between the LED and the fly antenna, the light intensity was equivalent to 1.13 W/m^2^. The antenna was stimulated for 500 ms followed by 5 s of recovery period for the total recording length of 20 s (3 stimulations). The identity of ab2 and ab3 sensilla were first verified with 1% ethyl acetate (Sigma #270989) and 1% pentyl acetate (Sigma #109584) before optogenetic experiments. Identification of ab9 sensilla was assisted by fluorescence-guided Single Sensillum Recording (fgSSR) (Lin and Potter, 2015) using *Or67b-Gal4* (BDSC #9995) recombined with *15XUAS-IVS-mCD8::GFP* (BDSC #32193). The Δ spikes/second was calculated as for other basiconic SSR recordings.

#### Single-nucleus RNA-sequencing Analyses

Dataset analyzed in this paper was published in (McLaughlin et al., 2021). The expression levels for the *Ir co-receptors* across all OSNs were lower than for *Orco*, even for their corresponding ‘canonical’ glomeruli. To account for these differences and facilitate comparisons, we performed within-gene normalization in Figure 4-Source Data 1 and used the normalized values to generate the AL maps in Figure 4A. The normalization was performed as follows: first, we determined the fraction of cells within each cluster expressing the given co-receptor (read Counts Per Million, CPM threshold ≥3). The cluster with the highest fraction value was taken as the maximum. Then, the fraction for each cluster was divided by this maximum value. The normalized value shows the relative strength of expression within each cluster for the given co-receptor gene.

#### EM Neuron Reconstruction

VM6 OSNs (Figures 5F-G) were reconstructed in the FAFB EM volume (Zheng et al., 2018) using FlyWire (https://flywire.ai/) (Dorkenwald et al., 2020). Initial candidates were selected based on either being upstream of the VM6 (previously called VC5 in Bates et al., 2020) projection neurons or based on co-fasciculation with already identified VM6 OSNs. Analyses were performed in Python using the open-source packages *navis* (https://github.com/schlegelp/navis) and *fafbseq* (https://github.com/flyconnectome/fafbseg-py). OSNs were clustered using FAFB synapse predictions (Buhmann et al., 2021) for a synapse-based NBLAST (“syNBLAST”, implemented in *navis*). The reconstructions can be viewed at https://flywire.ai/#links/Task2021a/all.

### Quantification and Statistical Analysis

#### Cell Counting

To quantify knock-in co-expression with the corresponding antibodies (Figure 2), the 3D reconstruction software Amira (FEI, Oregon, USA) was used to manually mark individual cell bodies throughout the z-stack in each channel (antibody in far red channel, knock-in in green channel), and the cell markers between channels were compared.

For sacculus cell counts (Figure 5C-E), cells were counted within ImageJ/Fiji using the cell counter tool. Counts were done manually by going through each stack within an image and using different colored markers for each cell type. Cell count data was gathered into an Excel spreadsheet, and analyzed for percent colocalization with n=9 antennal sections.

#### Statistics

Statistical analyses on SSR recordings were done in GraphPad Prism (version 8), except for optogenetic experiments, which were analyzed in Microsoft Excel. Box plots were made using GraphPad Prism; bar graphs were made in Excel. For all analyses, significance level α = 0.05. The following analyses were performed on all SSR data (excluding optogenetics): within genotype, Kruskal-Wallis test with uncorrected Dunn’s to determine which odorant responses were significantly different from mineral oil or paraffin oil control; between genotype, Mann-Whitney *U* to compare responses of two genotypes to the same odorant (e.g. *wildtype* vs *Ir25a^2^* mutant, or *wildtype* versus *Orco-T2A-QF2* knock-in). Summary tables in Figure 6 are filled in based on the following criteria: no response means neither *wildtype* nor *Ir25a^2^* mutant odor-evoked activity for given odorant was significantly different from its respective mineral oil control, nor was the difference between the genotypes statistically significant; no difference means that either *wildtype* or mutant or both had a significantly different odor-evoked response to the stimulus compared to mineral oil control, but the difference between the two genotypes was not statistically significant; higher response (in either *wildtype* or mutant) means that there was a statistically significant difference between genotypes for the given odorant. This could mean that: a) one genotype did not have a response, while the other did; b) both genotypes had a response, and one was higher; c) responses are different from each other, but not from their respective mineral oil controls; or d) neural activity was inhibited by the odorant in one genotype compared to mineral oil control, and either not inhibited in the other genotype or inhibited to a lesser degree. Non-parametric tests were chosen due to small sample sizes and/or data that were not normally distributed.

In Figure 6I, stimulus responses which were statistically significantly different from mineral oil control were those whose Δ spike values were zero, due to the fact that the mineral oil control Δ spike value was non-zero (median = 1.2, range = 0 to 2). Because of this, we did not deem these differences as biologically relevant. Nevertheless, *p* values are reported in the figure legend of Figure 6, and detailed information can be found in Figure 6-Source Data 1.

## Figure Supplements

**Figure 2-Figure Supplement 1.**
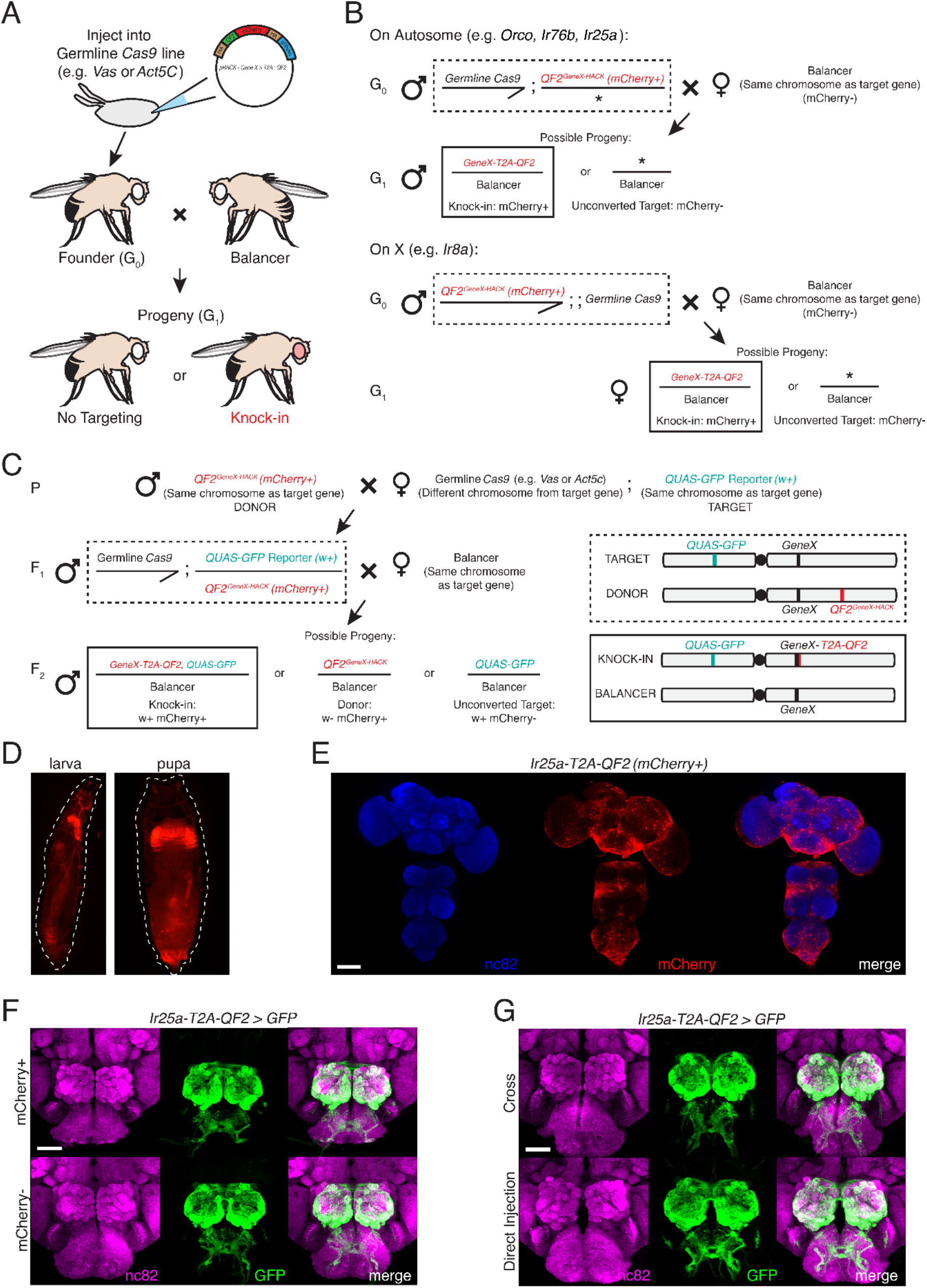
HACK Crossing Schematics, Marker Expression and Approach Comparison. **A.** Schematic of the direct injection method to generate a HACK knock-in. The HACK KI construct is injected into a germline Cas9 expressing embryo (both *Act5C-Cas9* and *Vas-Cas9* yielded successful knock-ins; see Methods for more details). This approach requires one genetic cross to generate a knock-in: injected founder flies (G_0_) are individually crossed to balancer flies, and progeny (G_1_) are screened for the mCherry knock-in marker. Each mCherry+ G_1_ represents an independent knock-in event; thus, single mCherry+ flies are used to establish knock-in lines. **B.** Details of the genetic crosses used to establish knock-in lines for each target gene using the direct injection approach. For genes on autosomes (*Orco* and *Ir76b*, 3^rd^ chromosome; *Ir25a*, 2^nd^ chromosome), a *Cas9* line on the X chromosome was used, and mCherry+ male progeny selected to remove the *Cas9* after successful HACKing. For *Ir8a* on the X chromosome, a *Cas9* line on the 3^rd^ chromosome was used which was marked with GFP to facilitate its removal in subsequent generations. In addition to mCherry marker screening, PCR genotyping, and sequencing, knock-in lines were confirmed by crossing to a *QUAS* reporter line and checking brain expression, either at the G_2_ stage, or by crossing G_0_ flies to balanced reporter lines to immediately check G_1_ brain expression. **C.** Details of using the crossing method to generate HACK knock-ins. This involves the creation of donor lines, in which the HACK targeting construct is inserted into the genome, such as through *P*-element insertion or *ΦC31* integration. Donor lines are crossed to a germline *Cas9* line (Parental). The progeny, F_1_, have all the components necessary for a knock-in to occur (gRNAs, Cas9, HDR cassette). An additional cross is required to select germline HACKing events at the F_2_ generation. Because the donor lines also express mCherry, as do the knock-ins, the target chromosome must be marked with a different marker (such as w+) for knock-in selection (see chromosome schematics at the right, illustrating the locations of the genetic elements in the F_1_ and F_2_ generations). F_2_ knock-ins will have both markers (in this case, mCherry+ w+), while donor flies will only have one (mCherry+ w-). The cross approach was tested with two of the four genetic targets (*Orco* and *Ir25a*) and successfully produced knock-in lines for both. However, the cross approach can produce false positives (see Table 1 and Table 1-Source Data 1) and is more time consuming. The direct injection approach is recommended for generation of targeted knock-ins. **D.** Knock-in screening can be performed at the larval (left) and pupal (right) stages. **E.** The *3XP3-mCherry* marker is expressed throughout the adult fly brain and ventral nerve cord. Nc82 channel has been pseudo-colored blue for clarity. **F.** The marker cassette is flanked by *loxP* sites and can be removed through *Cre* recombination (see Methods for details) (Siegal and Hartl, 1996). The *3XP3-mCherry* marker does not affect knock-in expression. *Ir25a-T2A-QF2*-driven GFP expression in the brain was similar before (top row) and after (bottom row) *Cre*-mediated removal of the mCherry marker. **G.** No difference was found in the expression patterns of *Ir25a-T2A-QF2* knock-in lines generated by the two HACK methods, genetic cross (top) and direct injection (bottom). For (F) and (G): N = 5 for direct injection; N = 4 for cross, mCherry+; N = 3 for cross, mCherry-. Expression patterns were comparable across all three groups. See also Table 1 and Table 1-Source Data 1. Scale bars: 100 µm in (E), 50 µm in (F-G).

**Figure 2-Figure Supplement 2.**
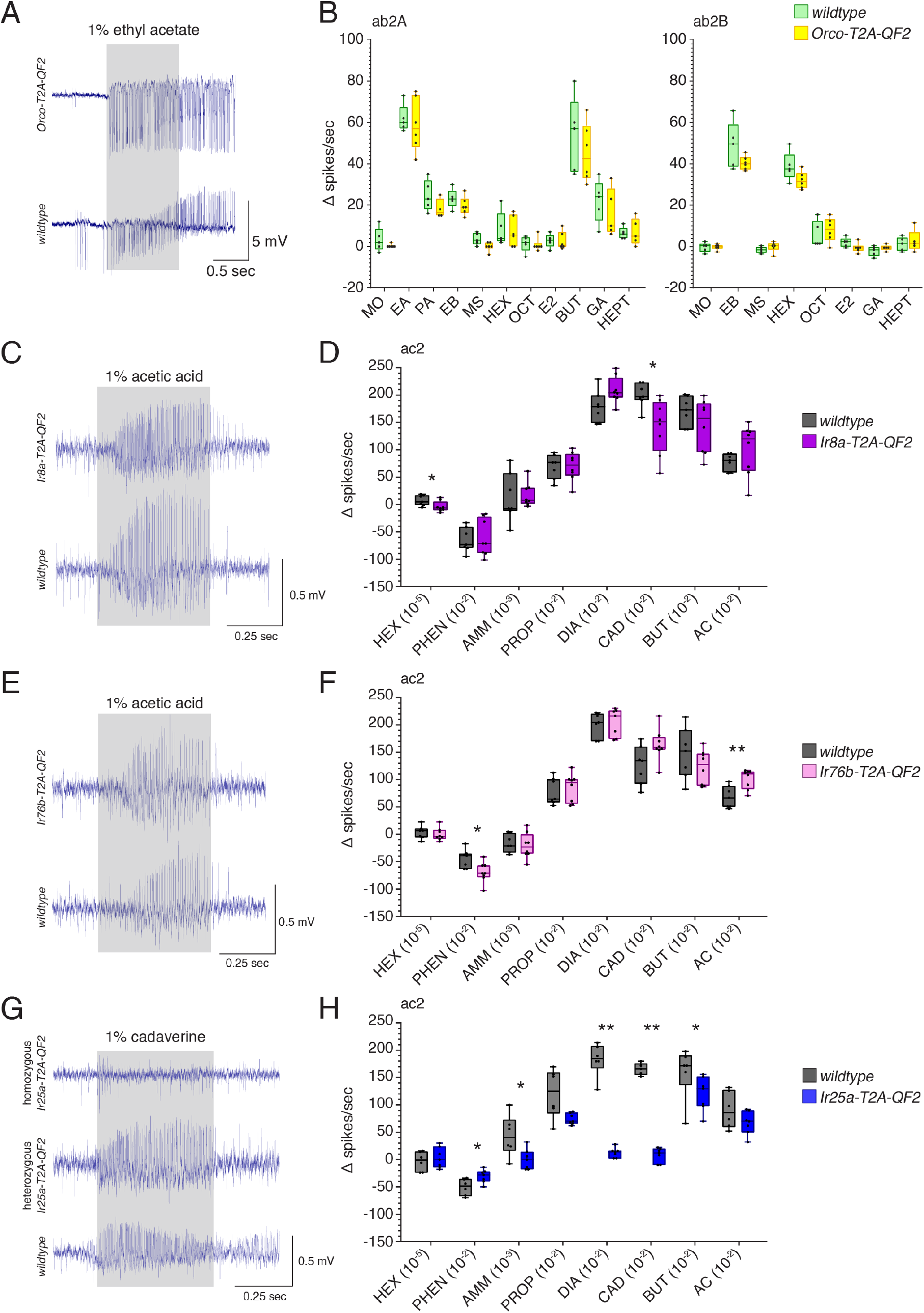
T2A-QF2 HACK knock-in Effects on Target Gene Function. **A.** *Orco* knock-in SSR experiments were performed in ab2 sensilla. Responses to ten odorants plus mineral oil control were compared in *wildtype* and homozygous *Orco-T2A-QF2* sensilla. Shown are example responses to 1% ethyl acetate. **B.** ab2 sensilla contain two neurons: A (left) and B (right). No differences were found in the responses of these neurons between the two genotypes across all odorants tested (Mann-Whitney *U* test, *p* > 0.05). A small difference in baseline (pre-stimulus) activity was observed, which can be seen in (A). Fewer stimuli are shown for ab2B because odorants strongly activating the A neuron obscured the B neuron responses. N = 5 for *wildtype*, N = 6 for *Orco-T2A-QF2*. Abbreviations: MO mineral oil, EA ethyl acetate, PA pentyl acetate, EB ethyl butyrate, MS methyl salicylate, HEX hexanol, OCT 1-octen-3-ol, E2 e2-hexenal, BUT 2,3-butanedione, GA geranyl acetate, HEPT 2-heptanone. **C**.-**H**. *IrCo* knock-in SSR experiments were performed in ac2 sensilla. Responses to eight odorants were compared in *wildtype* and homozygous knock-in flies. **C.** Example responses to 1% acetic acid in *Ir8a-T2A-QF2* and *wildtype* sensilla. **D.** There was a small but significant difference in responses between the two genotypes to hexanol (Mann-Whitney *U* test, *p* = 0.0286) and cadaverine (Mann-Whitney *U* test, *p* = 0.0106), which are not typically considered to be Ir8a-dependent odorants (Silbering et al., 2011). Responses to all other stimuli were not significantly different between genotypes. N = 7 for *wildtype*, N = 8 for *Ir8a-T2A-QF2*. **E.** Example responses to 1% acetic acid in *Ir76b-T2A-QF2* and *wildtype* flies. **F.** There were small but significant differences in responses to phenethylamine (Mann-Whitney *U* test, *p* = 0.012) and acetic acid (Mann-Whitney *U* test, *p* = 0.0087) between *Ir76b-T2A-QF2* and *wildtype* flies. Responses to the other stimuli were not significantly different between genotypes. N = 7 for *wildtype*, N = 8 for *Ir76b-T2A-QF2*. **G.** Homozygous *Ir25a-T2A-QF2* flies lost responses to most amines, recapitulating an *Ir25a* mutant phenotype (Abuin et al., 2011; Silbering et al., 2011). Heterozygous *Ir25a-T2A-QF2* flies with one *wildtype* copy of *Ir25a* had normal responses. Shown are example responses to 1% cadaverine in homozygous *Ir25a-T2A-QF2*, heterozygous *Ir25a-T2A-QF2*, and *wildtype* ac2 sensilla. **H.** Homozygous *Ir25a-T2A-QF2* flies had strongly reduced or abolished responses to phenethylamine (Mann-Whitney *U* test, *p* = 0.039), ammonia (Mann-Whitney *U* test, *p* = 0.0338), 1,4-diaminobutane (Mann-Whitney *U* test, *p* = 0.0012), cadaverine (Mann-Whitney *U* test, *p* = 0.0025), and 2,3-butanedione (Mann-Whitney *U* test, *p* = 0.0472). Responses to the other stimuli were not significantly different between genotypes. N = 6 for *wildtype*, N = 7 for homozygous *Ir25a-T2A-QF2*. Abbreviations in (D-H): PO paraffin oil, HEX hexanol, PHEN phenethylamine, AMM ammonia, PROP propanal, DIA 1,4-diaminobutane, CAD cadaverine, BUT 2,3-butanedione, AC acetic acid. See Figure 2-Source Data 1 for all *U* and *p* values.

**Figure 2-Figure Supplement 3.**
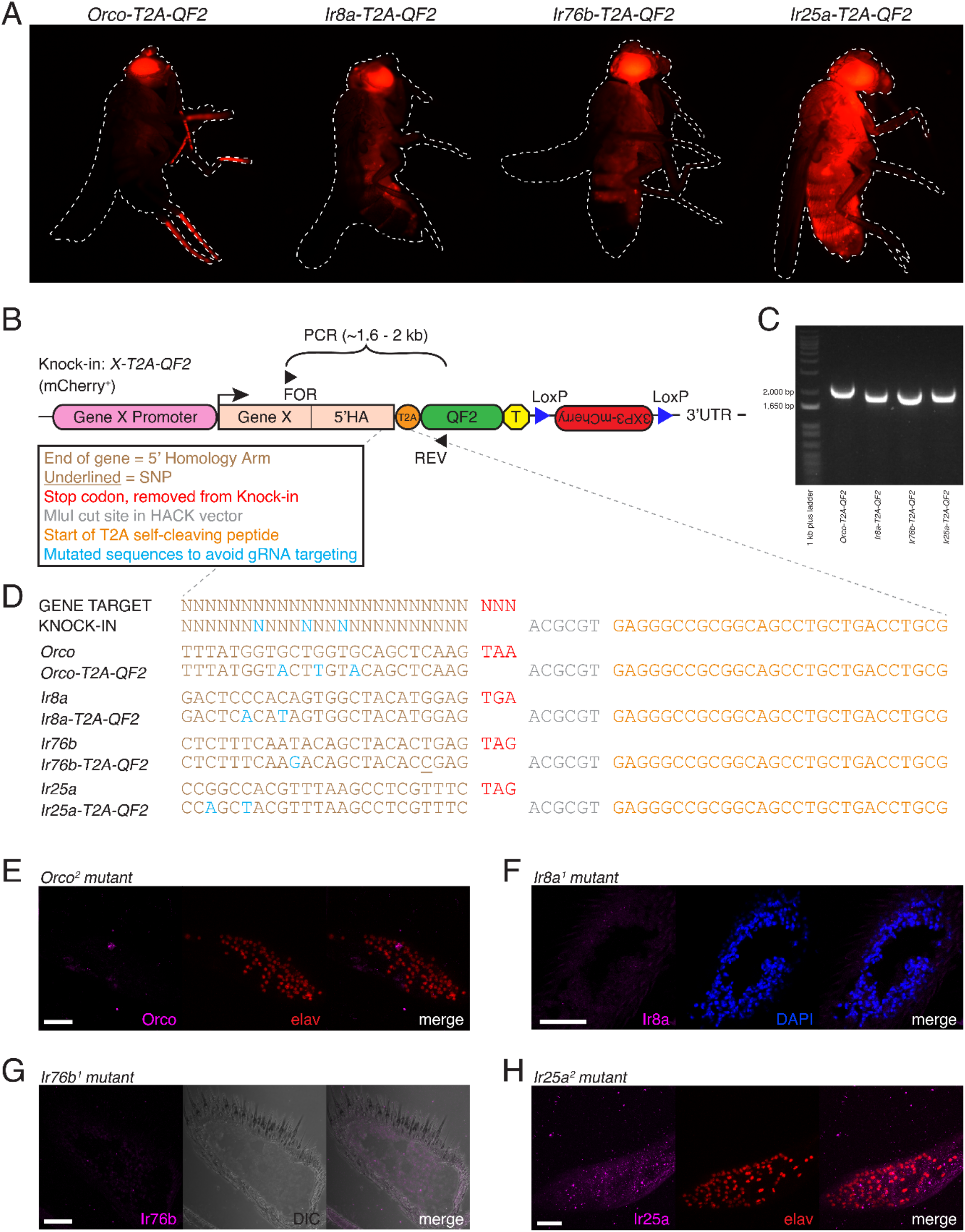
Additional Validation of Co-Receptor Knock-in Lines. **A.** The HACK *3XP3-mCherry* marker is expressed in the eyes and broadly throughout the fly nervous system and is subject to positional effects in expression. **B.** Each knock-in line is generated by CRISPR-Cas9 mediated homology-directed insertion of a *T2A-QF2* cassette marked by mCherry in front of the translational stop codon of the target gene. The 5’ homology arm (5’HA) is ∼1kb of the gene just before the stop codon. Successful knock-in lines were confirmed using forward primers upstream of the 5’HA (FOR), and a reverse primer within the *QF2* sequence (REV). Amplification of a ∼1.6 to 2 kb sequence occurs only if a knock-in is present. See Figure 2A for target and donor construct schematics. **C.** PCR genotyping using the primers shown in (B) confirms successful knock-in of the four targeted genes. **D.** Knock-ins were confirmed with sequencing of the targeted genomic site. For each co-receptor, the first line shows the target sequence (based on the FlyBase reference genome) (Thurmond et al., 2019), and the second line shows the verified sequence of the knock-in. The stop codon (red) has been replaced with the HACK *T2A-QF2* cassette. The portion shown here includes the *MluI* restriction site used for cloning the 5’HA (grey), as well as the start of the *T2A* sequence (orange). In each knock-in construct, synonymous substitutions (blue) have been made to prevent the gRNA/Cas9 from cutting the donor construct. Each knock-in construct was designed with three substitutions. The underlined base pair in the *Ir76b* knock-in is a SNP discovered in the lab *wildtype* stock and considered when designing the knock-in construct. **E.-H.** Antibody and *in situ* probe validation. **E.** No anti-Orco expression in *Orco^2^* mutant palps. elav is used as a pan-neuronal counterstain. **F.** No anti-Ir8a expression in *Ir8a^1^* mutant antennae. DAPI is used as a cellular counterstain. **G.** No *Ir76b* probe signal in *Ir76b^1^* mutant antennae. DIC used to visualize tissue. **H.** No anti-Ir25a expression in *Ir25a^2^* mutant palps (elav counterstain). Scale bars = 25 µm. See also Table 2.

**Figure 2-Figure Supplement 4.**
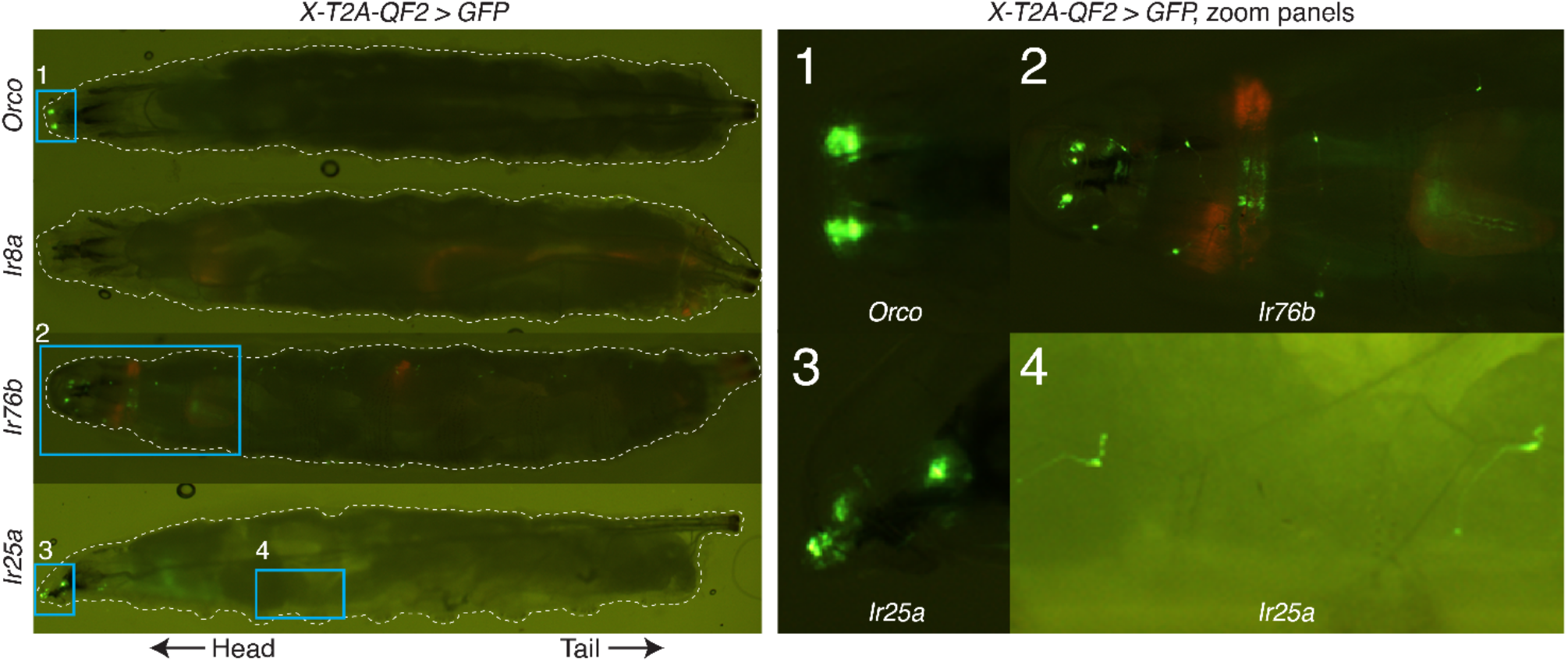
Knock-in Expression in the Larva. In the larval stage, *Orco-T2A-QF2* (first row) appears to drive expression only in the dorsal organs (zoom panel 1), as previously described (Larsson et al., 2004); *Ir8a-T2A-QF2* (second row) does not appear to drive GFP expression in the larva (although very weak expression may not be visible at this magnification); and both *Ir76b-T2A-QF2* (third row) and *Ir25a-T2A-QF2* (fourth row) drive GFP expression in the larval head as well as throughout the body wall (zoom panel 2 for *Ir76b*; zoom panels 3 and 4 for *Ir25a*), reflecting the role of these genes in additional, non-olfactory modalities. Weak red fluorescence from the *3XP3-mCherry* knock-in marker can be detected in the *Ir8a-T2A-QF2* and *Ir76b-T2A-QF2* larvae. This is due to channel bleed-through and is not seen in the other knock-in lines (*Orco-T2A-QF2* and *Ir25a-T2A-QF2*) that have had the marker removed (see Methods for details).

**Figure 3-Figure Supplement 1.**
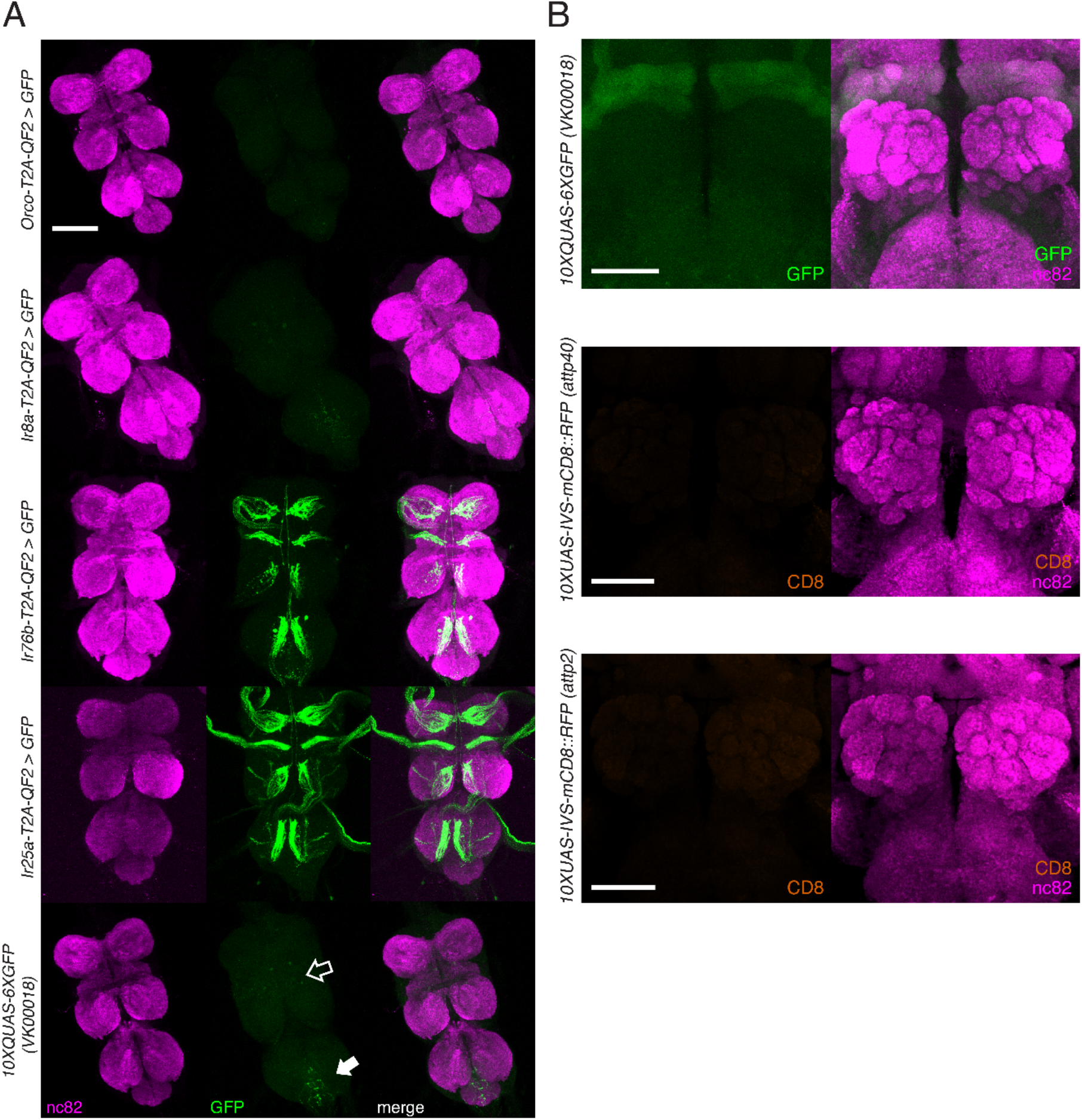
Knock-in Expression in the Adult Ventral Nerve Cord and Reporter Expression in the Brain. **A.** In adult flies, two of the four knock-ins (*Ir76b-T2A-QF2*, third row, and *Ir25a-T2A-QF2*, fourth row) drive GFP expression in neurons innervating the ventral nerve cord (VNC). This likely reflects the role of these co-receptors in gustation. There is weak GFP expression in the reporter control (bottom row) in the abdominal neuromere (filled arrow), as well as the accessory mesothoracic neuropil (empty arrow), which can be seen in the two other knock-ins (*Orco-T2A-QF2*, first row, and *Ir8a-T2A-QF2*, second row). This non-specific expression is likely from the reporter itself rather than driven by the two knock-ins. **B.** Control brains showing the *QUAS-GFP* reporter alone (top) and the *UAS-mCD8::RFP* reporters alone (middle and bottom). These reporters were used for all brain images in Figures 3 – 5. The *QUAS* reporter weakly labels the mushroom bodies. The ventral nerve cord from the fly in (A, fourth row) is also used in Figure 2-Figure Supplement 1E. Scale bars: 100 µm in (A), 50 µm in (B).

**Figure 3-Figure Supplement 2.**
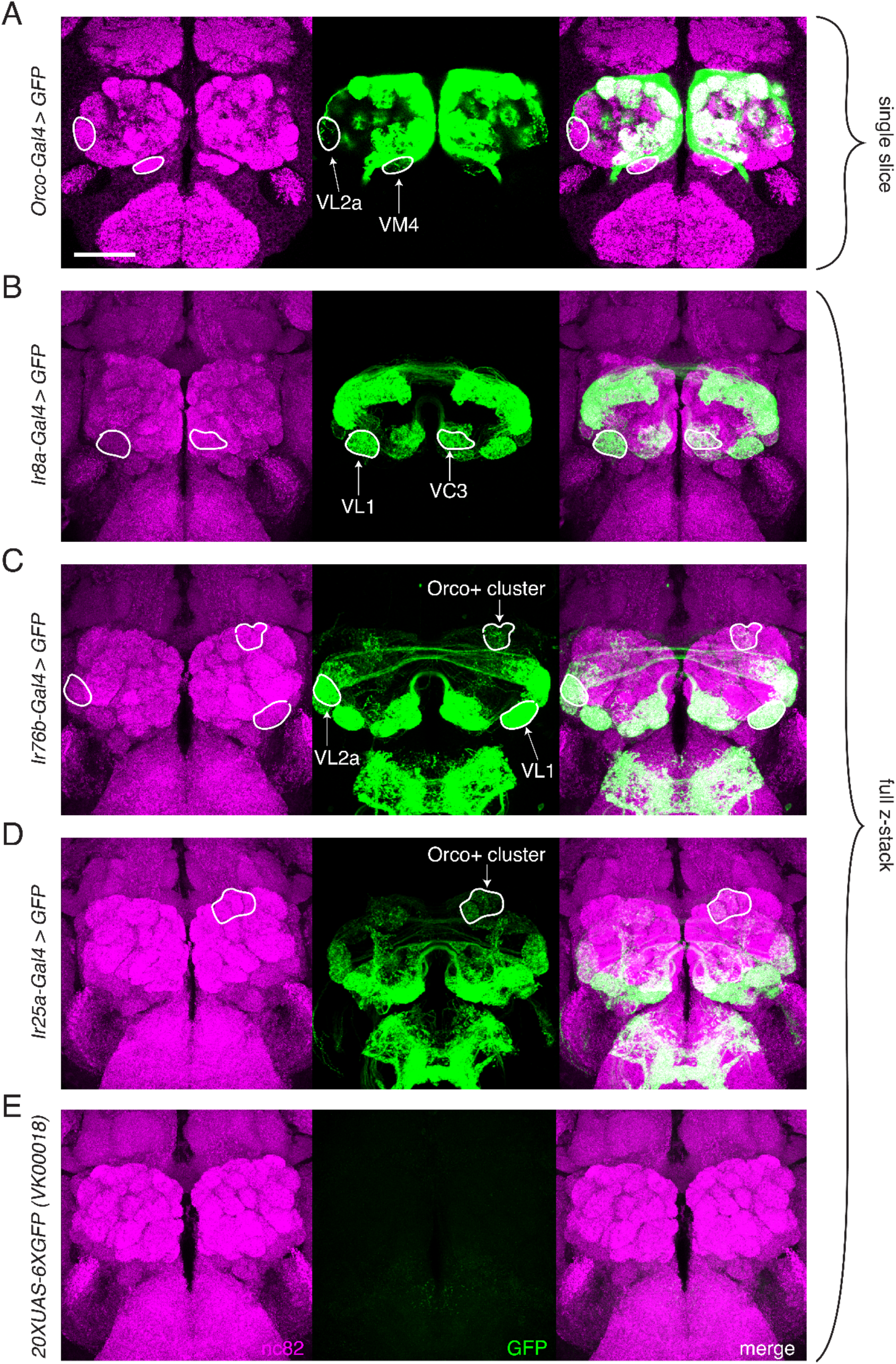
Transgenic Co-Receptor *Gal4* Lines Do Not Fully Recapitulate Knock-in Expression. **A.-E.** Transgenic co-receptor *Gal4* lines crossed to a strong *UAS* reporter reveal expression in some, but not all, of the additional glomeruli labeled by the respective knock-ins. Some discrepancies between the *Gal4* and knock-in lines are enumerated here, but do not represent a comprehensive list. **A.** *Orco-Gal4* drives weak GFP expression in VL2a and VM4 glomeruli (outlined), but lacks expression in DL2, V, and VL1 glomeruli, which are labeled by the knock-in. **B.** *Ir8a-Gal4>GFP* labels VC3 and VL1 (outlined), but lacks expression in the VM4 and VA3 glomeruli, which are labeled in the knock-in. **C.** *Ir76b-Gal4* labels the VL1 and VL2a glomeruli (outlined), but lacks expression in the VA5 and DC3 glomeruli which are consistently labeled in the knock-in. Additionally, it drives GFP expression in Orco+ glomeruli not seen in the knock-in (outlined). **D.** *Ir25a-Gal4* labels some Orco+ glomeruli (outline includes DA2, DA3, DA4m, and DA4l), but is lacking expression in many glomeruli labeled by the knock-in. **E.** The *UAS-GFP* control line has weak leaky expression in the SEZ but no expression in the AL. (A) shows single slice to visualize weakly labeled glomeruli; (B-E) are maximum intensity projections of z-stacks. Scale bar = 50 µm. See also Figure 3-Source Data 1.

**Figure 4-Figure Supplement 1.**
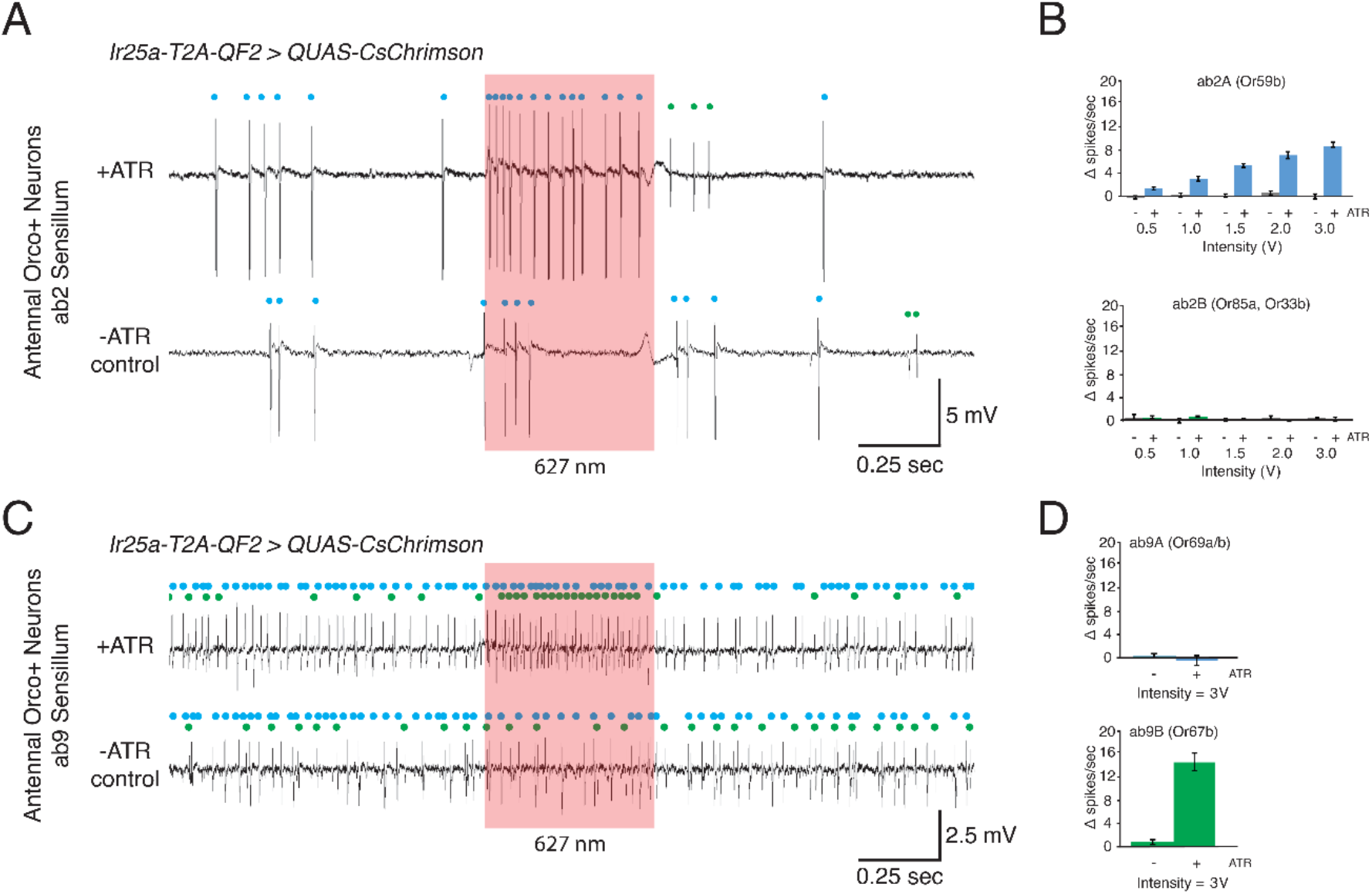
Optogenetic Experiments to Examine *Ir25a* Expression in Orco+ Neurons. **A-D.** Confirmation of *Ir25a* expression in ab2 and ab9 sensilla using optogenetic stimulation. *CsChrimson* expression is driven by *Ir25a-T2A-QF2*, and recordings are performed in neurons previously determined to express only *Orco*. **A.** Representative SSR traces from ab2 using 1.5V of a 627 nm LED light (red box) to activate *CsChrimson*. Bottom trace is control animal which was not fed the necessary cofactor all-trans retinal (-ATR). ab2 has two neurons: blue dots indicate A neuron spikes, while green dots indicate B neuron spikes (genotype: *Ir25a-T2A-QF2*; *QUAS-CsChrimson*). **B.** Quantification of neuronal activity in response to light stimulation at various intensities (N = 7 – 12). Optogenetic experiments confirm *Ir25a* expression in ab2A (expressing *Or59b*, top; DM4 glomerulus) but not ab2B (expressing *Or85a* and *Or33b*, bottom; DM5 glomerulus). **C.** Combination of optogenetics and fluorescent guided SSR (Lin and Potter, 2015) to examine *Ir25a* expression in ab9 sensilla (genotype: *Ir25a-T2A-QF2*; *QUAS-CsChrimson/Or67b-Gal4, 15XUAS-IVS-mcd8::GFP)*. Representative traces from ab9 in response to 3V of 627 nm LED light (red box). As in (A), -ATR indicates the inactive *CsChrimson* control; blue dots indicate A neuron spikes, green dots indicate B neuron. **D.** Quantification of activity in response to light stimulation verified that *Ir25a* is expressed in ab9B (*Or67b*, bottom; VA3 glomerulus) but not in ab9A neurons (*Or69aA/aB*, top; D glomerulus). N = 5.

**Figure 5-Figure Supplement 1.**
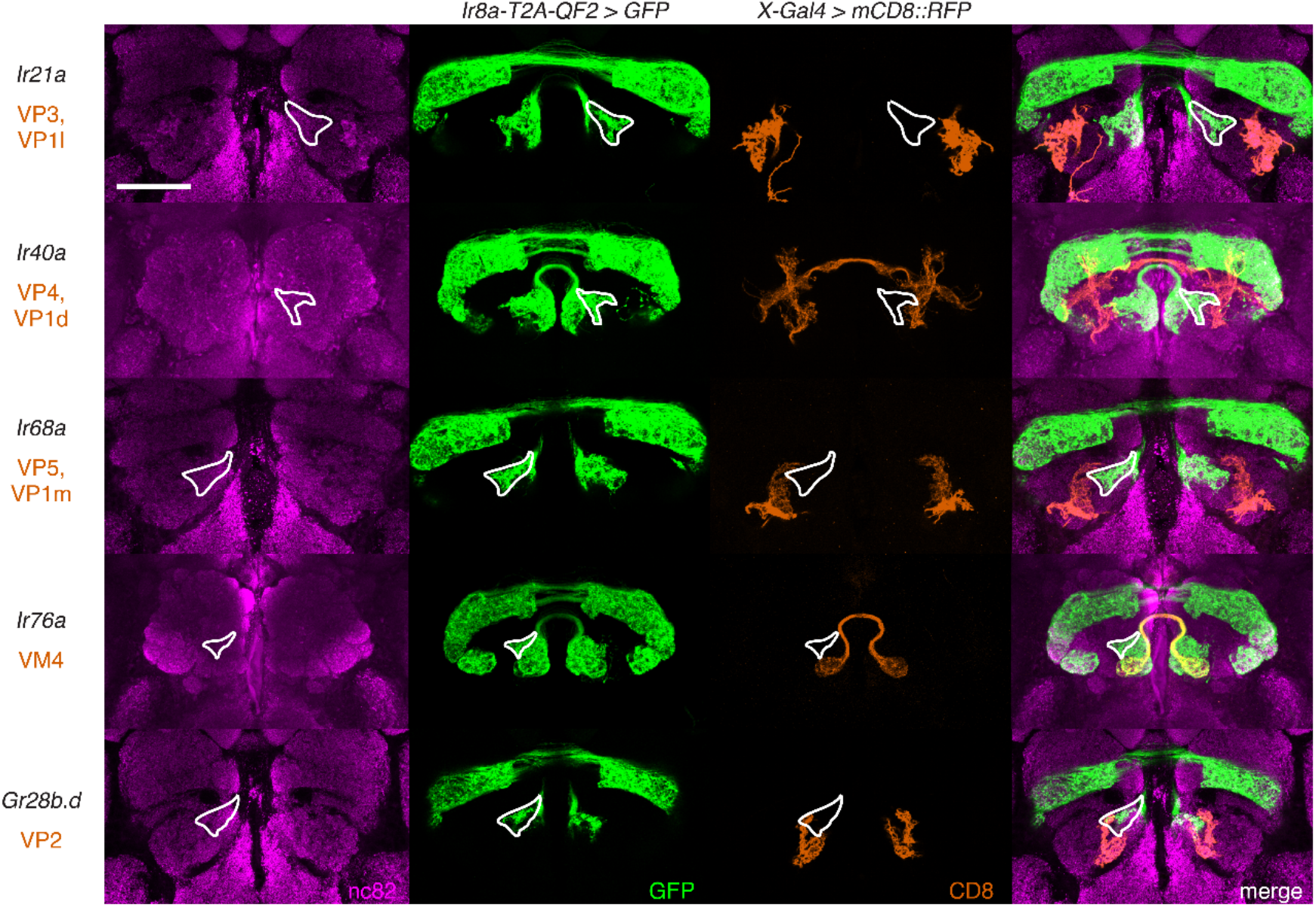
The New Glomerular Region Labeled by the *Ir8a* Knock-in Does Not Correspond to Previously Identified Posterior Glomeruli. Co-labeling experiments with *Ir8a-T2A-QF2 > GFP* (green) and various *Gal4* lines driving *mCD8::RFP* (orange) confirm that the outlined region does not correspond to any of the VP glomeruli nor to VM4. Scale bar: 50 µm.

**Figure 6-Figure Supplement 1.**
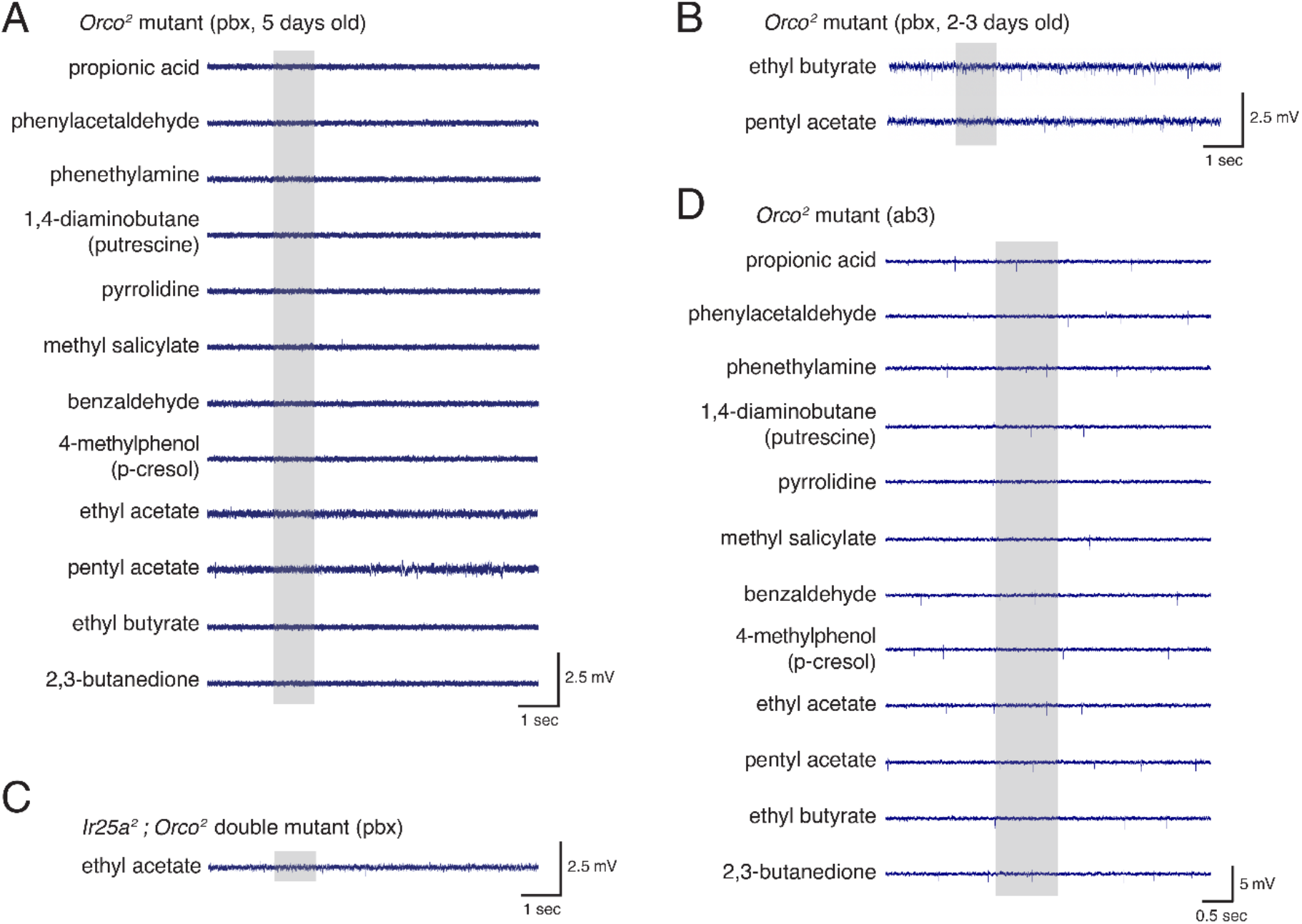
Electrophysiological Experiments to Examine *Ir25a* Function in Orco+ Neurons. **A.** SSR recordings in *Orco^2^* mutant palpal sensilla show neither baseline nor odor-evoked activity in response to a panel of odorants in 5-day old flies. N = 20 sensilla, 4 animals (3 male, 1 female). **B.** Young *Orco^2^* mutant palpal neurons also do not have spontaneous or odor-evoked activity. **C.** *Ir25a^2^; Orco^2^* double mutant palpal sensilla show neither baseline nor odor-evoked activity. N = 42 sensilla from 5 animals (3 male, 2 female). **D.** Antennal ab3 sensilla do occasionally show baseline activity in the *Orco^2^* mutant, but generally do not show odor-evoked activity (N = 5 flies; see also Figure 6). Grey box indicates the time of stimulus delivery. See also Figure 6-Source Data 1.

**Figure 6-Figure Supplement 2.**
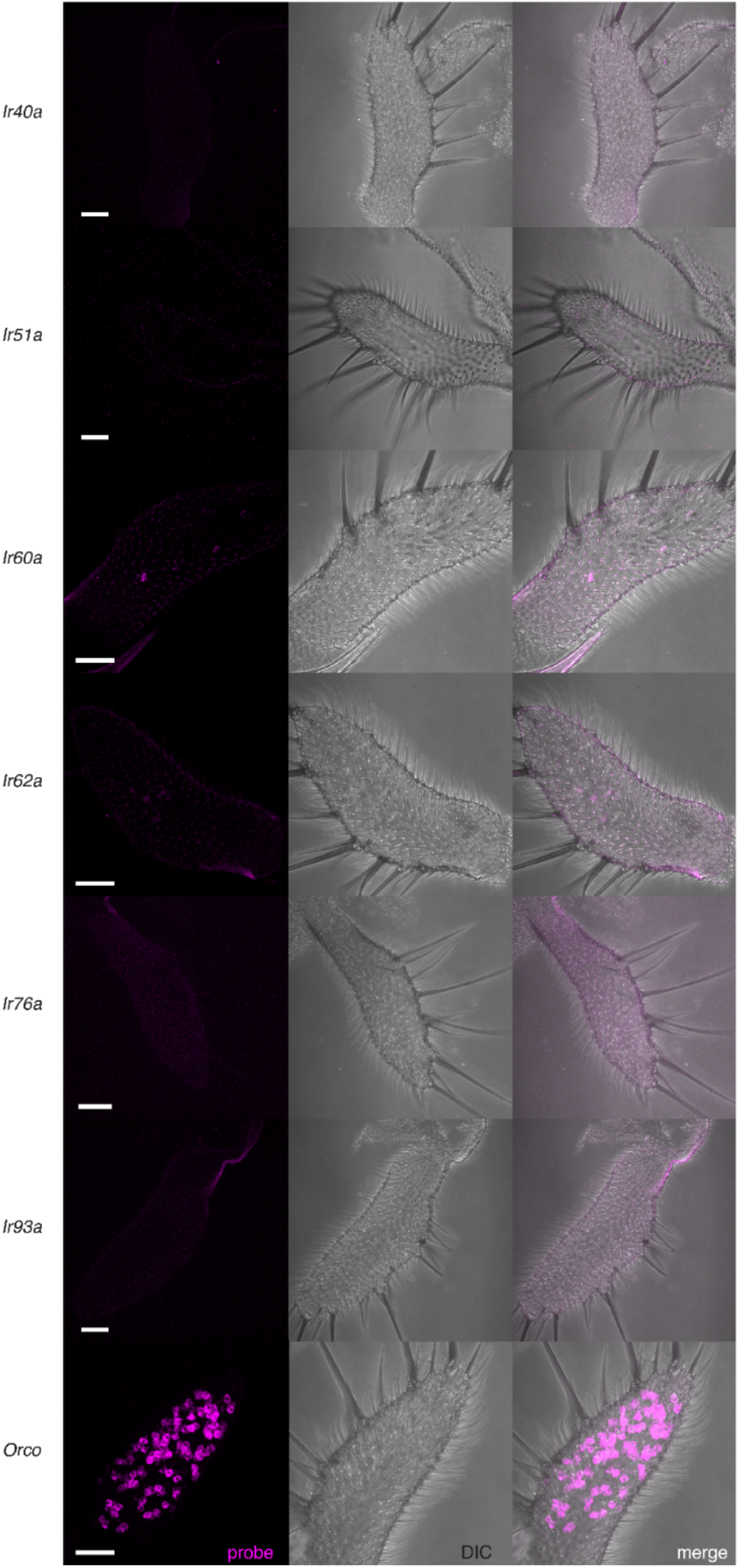
No IrX Expression of the Top Candidates in the Maxillary Palps. The recent fly cell atlas (Li et al., 2021) was used to identify potential IrX tuning receptors in the palps. *In situ* experiments did not reveal strong expression of any of these target genes. Orco (bottom) was used as a positive control. See also Figure 6-Source Data 2. Scale bars = 25 µm.

